# Climate warming weakens local adaptation

**DOI:** 10.1101/2020.11.01.364349

**Authors:** M. Bontrager, C. D. Muir, C. Mahony, D. E. Gamble, R. M. Germain, A. L. Hargreaves, E. J. Kleynhans, K. A. Thompson, A. L. Angert

**Affiliations:** Department of Botany and Biodiversity Research Centre, University of British Columbia, Vancouver, BC, Canada; School of Life Sciences, University of Hawai’i, Honolulu, HI, United States; British Columbia Ministry of Forests, Lands, Natural Resource Operations and Rural Development, Victoria, BC, Canada; Department of Ecology, Evolution, and Marine Biology, University of California, Santa Barbara, United States; Department of Zoology and Biodiversity Research Centre, University of British Columbia, Vancouver, BC, Canada; Department of Biology, McGill University, Montreal, Quebec, Canada; Departments of Botany and Zoology and the Biodiversity Research Centre, University of British Columbia, Vancouver, BC, Canada

## Abstract

Anthropogenic climate change is generating mismatches between the environmental conditions that populations historically experienced and those in which they reside. Understanding how climate change affects population performance is a critical scientific challenge. We combine a quantitative synthesis of field transplant experiments with a novel statistical approach based in evolutionary theory to quantify the effects of temperature and precipitation variability on population performance. We find that species’ average performance is affected by both temperature and precipitation, but populations show signs of local adaptation to temperature only. Contemporary responses to temperature are strongly shaped by the local climates under which populations evolved, resulting in performance declines when temperatures deviate from historic conditions. Adaptation to other local environmental factors is strong, but temperature deviations as small as 2°C erode the advantage that these non-climatic adaptations historically gave populations in their home sites.

**One sentence summary:** Climate change is pulling the thermal rug out from under populations, reducing average performance and eroding their historical home-site advantage.

---

Anthropogenic climate change is increasing the likelihood of extreme weather conditions that exceed current biological tolerances, posing severe threats to biodiversity [1]. While plasticity can help individuals maintain performance in the face of climatic variability, there are limits to its buffering capacity [2]. As a result, populations frequently adapt to local conditions, leading to differences among populations in climatic optima [3]. Conditions that deviate strongly from these optima can decrease organismal performance and threaten population persistence [4, 5]. To anticipate and mitigate the effects of global change on biodiversity, we must identify which climatic variables most affect performance and quantify the contribution of genetic differences among populations to species-level responses.

If populations within a species respond similarly to climatic variation, then performance effects will manifest regardless of geographic origin—for example, all populations of a species may perform well under moderate temperatures and poorly under extreme temperatures (Fig. 1A). If adaptation to local environments has occurred, population responses to climatic variation may also correspond to how strongly conditions deviate from those that they have historically experienced [6] (Fig. 1B). Local adaptation offers a pathway to species persistence as climate changes: declining populations may be rescued by migration of genotypes that are better adapted to contemporary conditions [7, 8]. However, adaptation to local environments also poses a challenge. If populations are strongly adapted to non-climatic components of the landscape (such as soils or competitors), then climate-tracking migration may disrupt these adaptations. Here, we paired a comprehensive quantitative synthesis of field transplant experiments with a new theoretical framework to evaluate whether responses to temperature and precipitation are locally adapted, quantify the magnitude of adaptation to these climatic variables vs. other components of the environment, and investigate the extent to which climate change is disrupting local adaptation.

**Fig. 1.**
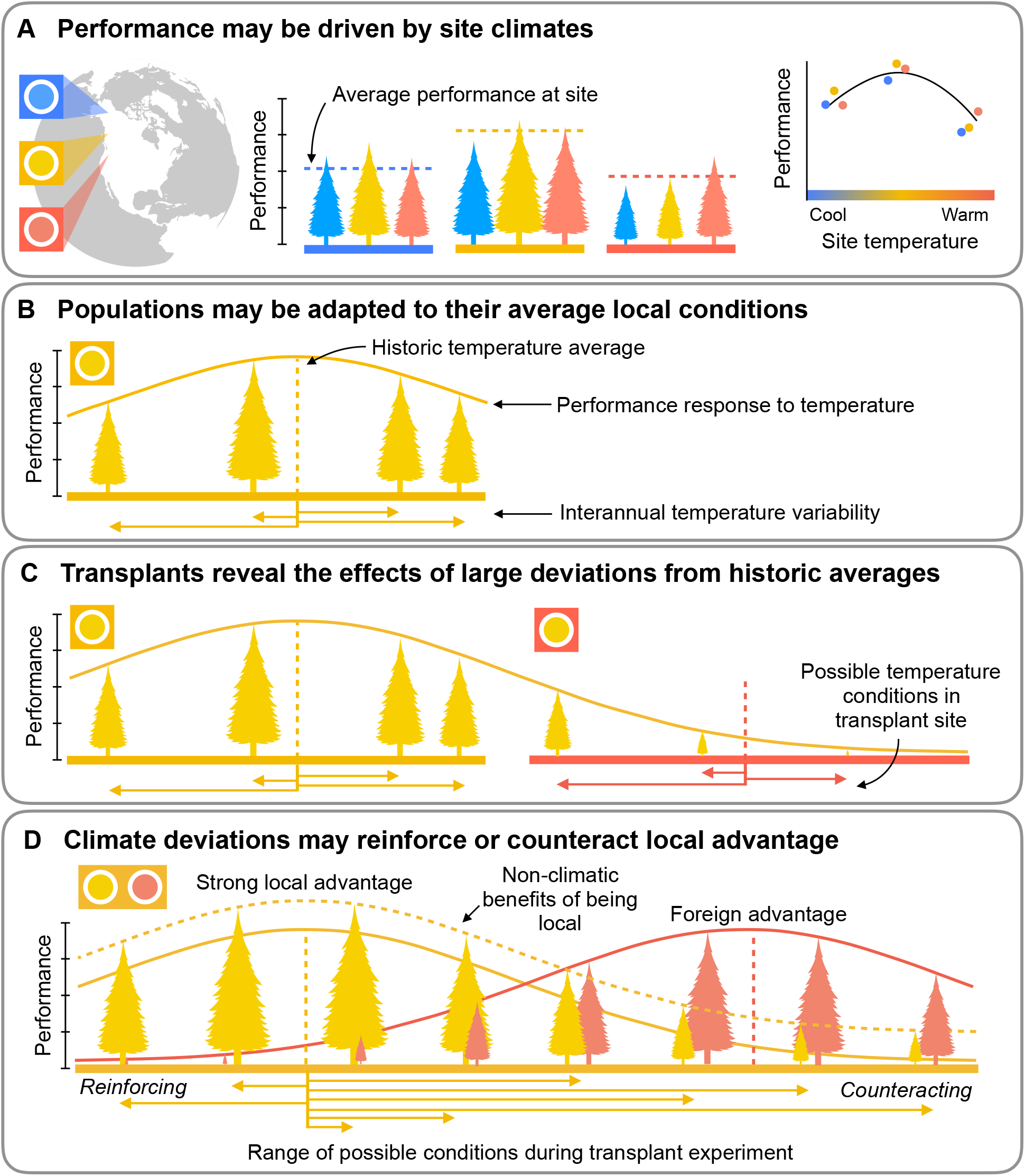
Transplant experiments, where individuals of different populations are moved among sites, are a powerful resource for understanding responses to climatic variation. (**A**) Site conditions may affect the performance of individuals, regardless of where they are from. (**B**) Populations may be adapted to the average climate in their home site, and their performance may decline when conditions deviate from these optima. (**C**) Transplant experiments allow us to observe population responses to a dramatic range of climate deviations. (**D**) The strength of local advantage may depend on the conditions of that site relative to the populations’ optima (the *relative experimental environment*) and the magnitude of adaptation to unmeasured factors.

Transplant experiments—in which individuals are moved between locations and their performance is measured—are a classic tool for parsing contributions of plasticity and genetic differentiation to variation in performance [9, 10]. Transplants offer an opportunity to examine population responses to an expanded range of climatic variation, including climates far outside the range of what populations have experienced in recent history (Fig. 1C). We compiled data from 1787 populations moved among 541 sites between 1967 and 2015, drawn from 147 published transplant studies within the native ranges of 164 species, the vast majority of which were plants (Materials and Methods 1.1–1.2, Table S1, Fig. 2AB). We used monthly temperature and precipitation records for each source population and transplant site to compare experimental conditions among transplant sites and to calculate how much these conditions deviated from the historic baselines (Materials and Methods 1.3, Fig. 2CD). Consistent with global trends, conditions during the transplant experiments in our dataset have become increasingly warm (Materials and Methods 3.1, Fig. 2C, Table S2A).

**Fig. 2.**
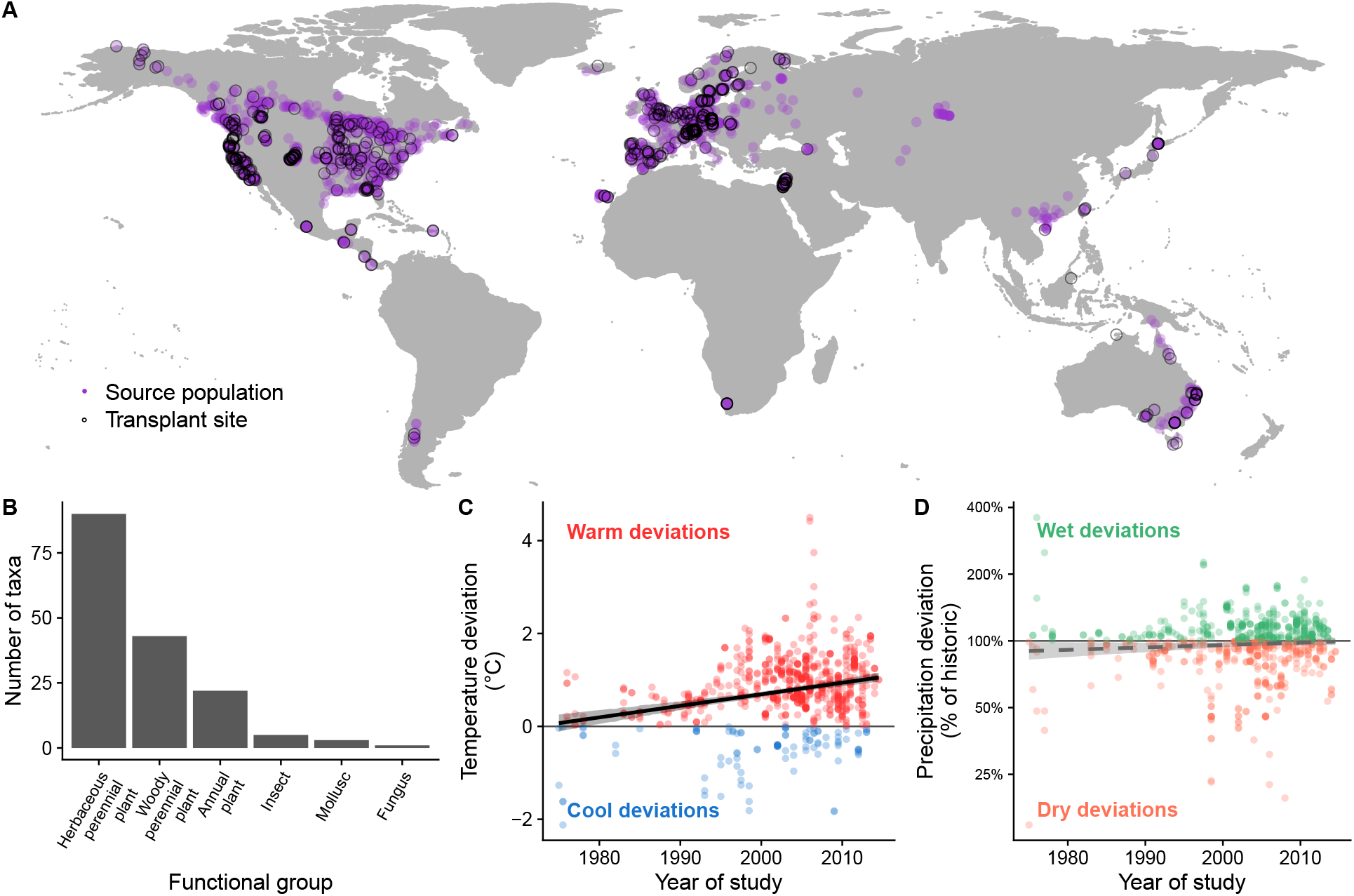
Experiments in the dataset encompass a wide range of climate deviations from historic averages. (**A**) Geographic locations of transplant sites (black circles) and source populations (purple points) were mostly in the northern hemisphere. (**B**) Most experimental taxa were plants. Across study sites, warm temperature deviations during transplant experiments became more prevalent over time (**C**), while precipitation conditions during experiments did not become either wetter or drier (**D**, nor did they become more variable, Table S2A). Shading represents 95% HDIs.

To characterize how sensitive performance was to the conditions at experimental sites, we centered temperature and precipitation around the average experimental conditions within each study, and quantified species’ average performance responses to this range of conditions (Materials and Methods 3.2). Performance was highest in transplant sites with relatively moderate temperatures (Fig. 3A, Table S3), and declined in sites that were more thermally extreme than experiment averages. For example, at a site 5°C warmer than a study’s average, performance suffered a 20% decline (95% highest density interval (HDI): 12 – 29%). Performance also declined in sites that were drier than experiment averages (Fig. 3B, Table S3).

**Fig. 3.**
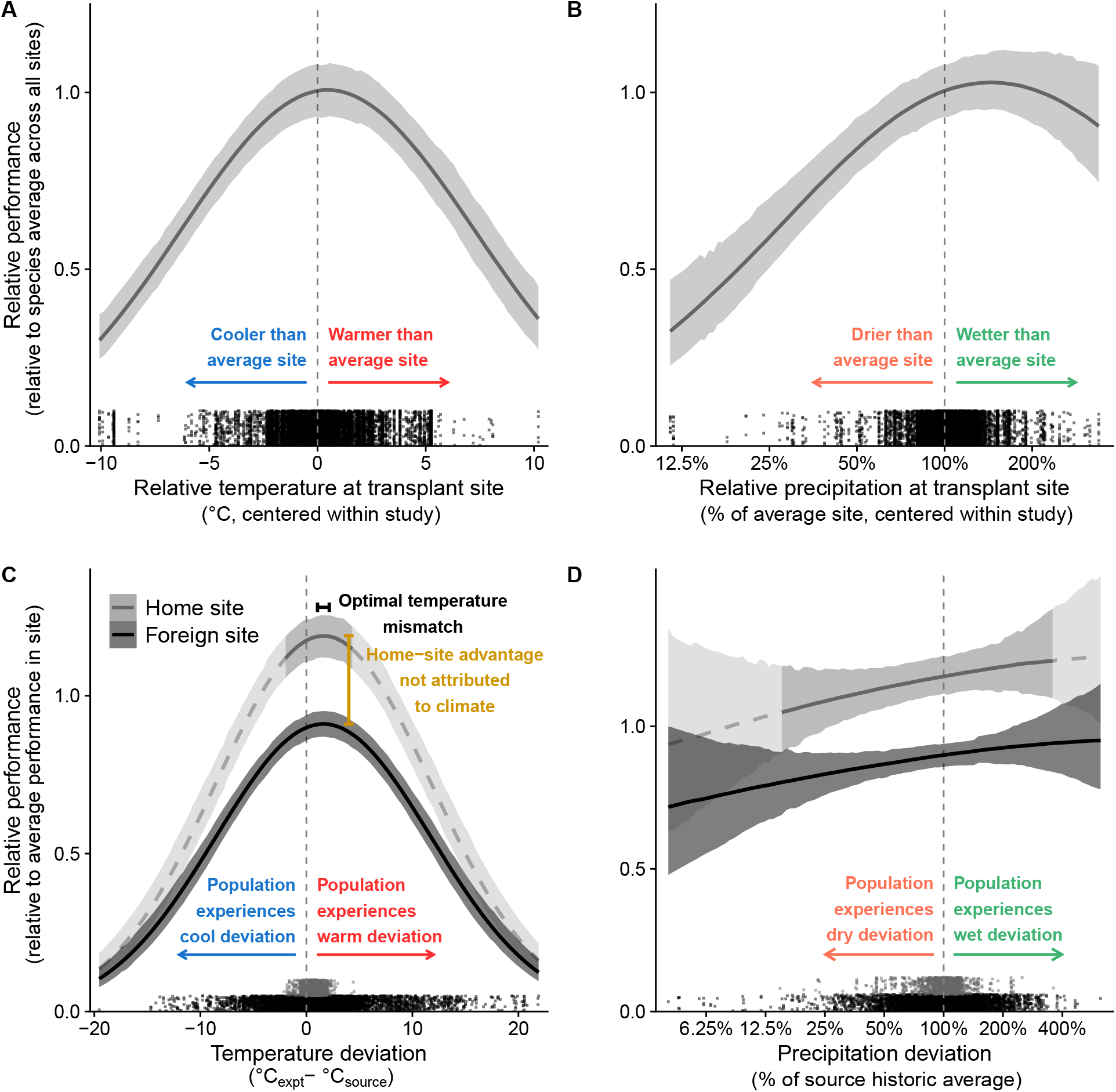
Populations exhibit both species-wide and locally adapted responses to deviations between experimental conditions and their historical climate. (**A** and **B**) Species’ mean performance is lower in sites with extreme temperature or precipitation conditions relative to other sites in the experiment. Temperature and precipitation values were centered around the average value within each study. Performance was averaged across all populations transplanted to a site and centered around the average performance of each species. (**C**) Relative performance declines when populations experience deviations from the historic average temperatures (1951-1980) in their home sites, but (**D**) does not decline with precipitation deviations. After controlling for climate deviations, local populations have a home-site advantage over foreign populations. The range of deviations captured in our database is indicated by the rugs of points. The darker portion of the local curve indicates the range of deviations that local populations experienced during transplants (i.e., due to interannual rather than spatial variation); the lighter portion is inferred from the model. In (**C**) and (**D**), performance is relative to other populations in the same site, removing any site-specific effects, such as those shown in (**A**) and (**B**). Shading represents 95% HDIs.

We then tested whether responses to climatic variation are locally adapted within species (Materials and Methods 3.2). The presence of local adaptation is frequently evaluated by comparing the performance of local and foreign populations across sites [11]. This quantifies the extent to which populations have adapted to the unique selection pressures of their home environments and reveals the effects of novel environments on populations. Among-site transplants allowed us to investigate performance responses to a broad range of deviations from the historic climates that populations have experienced: up to 21°C temperature differences and 6-fold differences in precipitation. We found that performance peaked when populations experienced temperatures slightly warmer than their historic average (95% HDI: 1.0 – 2.2°C, Fig. 3C, Table S4). This may reflect asymmetric effects of warm vs. cool temperatures, thermal safety margins, or recent adaptation to contemporary climate in some systems. Performance then declined away from this temperature optimum, consistent with local adaptation to historic temperature regimes. Transplanting populations to sites that deviated from their historic precipitation regimes did not affect their performance (Fig. 3D, Table S4).

As experiments are conducted under increasingly warm conditions, populations from historically warmer sites have been shown to outperform local populations at historically cool sites [12–14]. However, it is also plausible (though under-recognized) that at warm sites, the home-site advantage of warm-adapted populations could be enhanced. Thus, contemporary deviations from historic climate regimes may increase or decrease the strength of local advantage, depending on the populations being compared and the direction of the climatic deviation (Fig. 1D). To make quantitative predictions for the effects of experimental conditions on local advantage, we developed a new metric based in evolutionary theory (Materials and Methods 2). This metric, which we call the *relative experimental environment,* incorporates three variables—the historic climate of the *local* population, the historic climate of the *foreign* population, and the *experiment conditions—*into a single predictor (Materials and Methods 3.3). Higher values predict that the local population is better matched to the experimental climatic conditions, and therefore, local advantage should be strong (Materials and Methods 2.3). We tested whether these predictions were supported by experimental data, and found that the strength of local advantage is indeed positively correlated with the relative experimental temperature (Fig. 4A, Table S5).

**Fig. 4.**
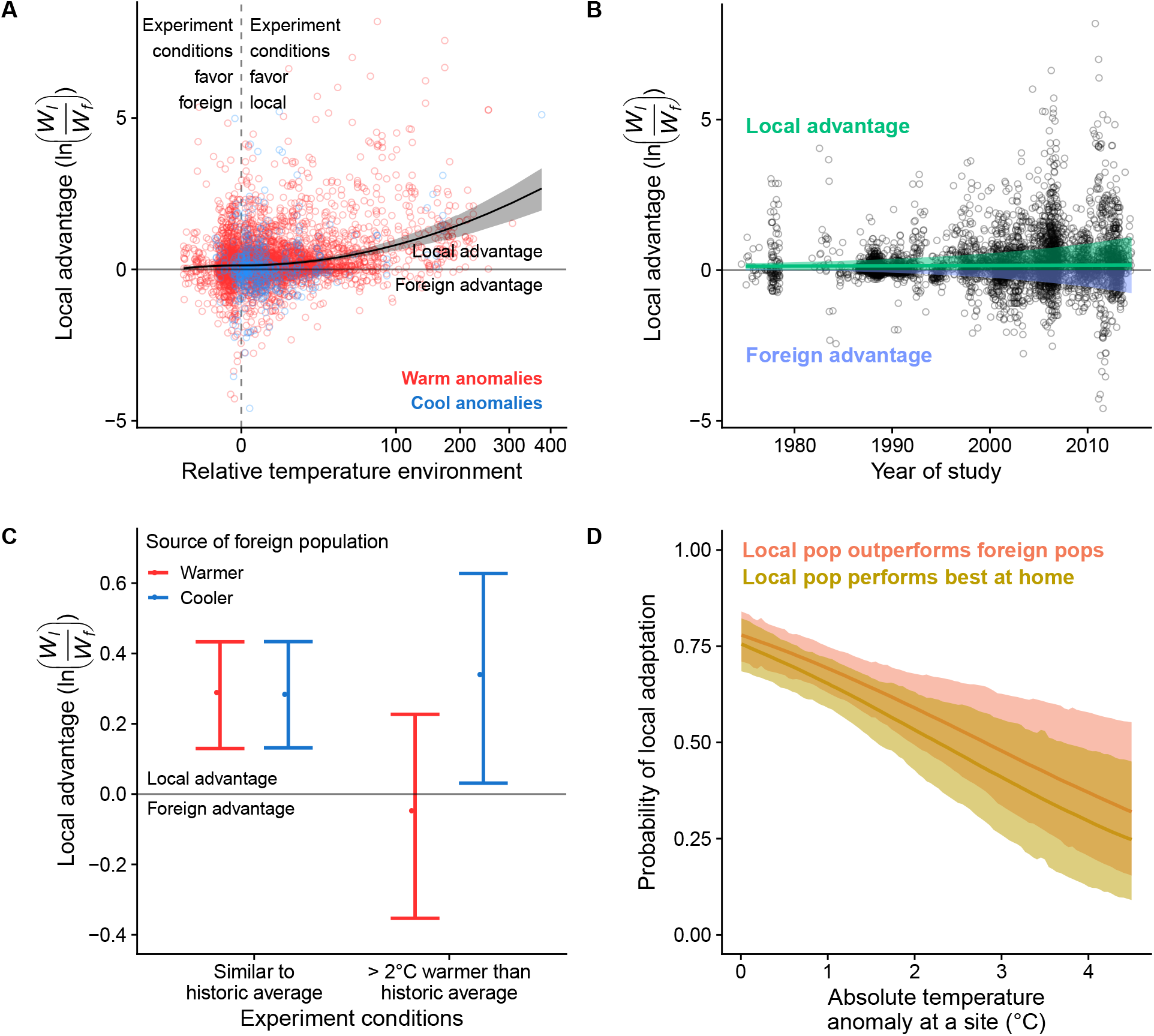
Experimental conditions affect the strength and prevalence of local advantage in transplant experiments. (**A**) Temperature deviations that favor the local population strengthen home-site advantage, while those that favor the foreign population reduce it. Local advantage is calculated as the natural log of the ratio of local to foreign performance. The relative temperature environment is calculated based on each population’s historic average temperature and the temperature during the transplant experiment (Materials and Methods 2 and 3.3). (**B**) The average strength of local advantage detected (solid green line) has not changed over time, but its variance has increased (shaded region). (**C**) When temperatures during transplant experiments are similar to historic averages, local advantage does not depend on the source temperature of the foreign population. When experiments are conducted under temperature deviations 2°C warmer than historic averages, local populations lose their advantage when compared to warmer foreign populations but maintain their advantage when compared to cooler foreign populations. (**D**) Temperature anomalies decrease the probability that the local population will outperform all foreign populations in a site (red line) and that a population moved to multiple sites will perform best in its home site (gold line). Shading represents 95% HDIs.

Because climate deviations sometimes weaken and sometimes strengthen local advantage, our theory shows that their net effect should increase the variance in the strength of local adaptation (Materials and Methods 2.5). Consistent with this prediction, we detected a nearly ten-fold increase in the variance of local advantage, from 0.049 in 1975 (95% HDI: 0.044 – 0.055) to 0.47 in 2015 (95% HDI: 0.44 – 0.51) (Fig. 4B, Materials and Methods 3.3). However, very little of the increased variance in local advantage can be attributed directly to temperature or precipitation deviations *per se* (Materials and Methods 3.4, Supplementary Results 4.4). Nevertheless, we found striking evidence that climate warming is reducing local adaptation: 1) when temperatures deviated more than 2°C above historic averages, the advantage of local populations over warmer foreign populations vanished (Fig. 4C, Table S6); 2) the probability that the local population outperforms all foreign populations in a site decreased as temperatures deviated from their historic averages (Materials and Methods 3.5, Supplementary Results 4.5, Fig. 4D, Table S7A); and 3) temperature deviations decreased the probability that a population did best at its home site compared to other sites to which it was moved (Fig. 4D, Table S7B).

Such a strong signature of decreased local adaptation in the face of temperature deviations is remarkable because adaptation to local environments is multidimensional, encompassing not just climate but also microhabitat characteristics [15] and biotic factors such as competitors [16, 17], pathogens [18], and predators [19, 20]. After accounting for the effects of temperature and precipitation, we found that foreign populations were still on average 24% less fit than local populations (Fig. 3C). This indicates that populations that move to track their climate optimum may suffer from leaving their historic biotic or edaphic environments. Nonetheless, analyses that account for environmental and spatial differences between local and foreign populations (Fig. 4CD, Table S6, S7) suggest that as little as 2°C warming can overwhelm all historical non-climatic and climatic adaptations. To date, syntheses of local adaptation have focused on evaluating its ubiquity and quantifying its average strength [21, 22] (but see [23, 24]). In contrast, our work partitions the effects of temperature and precipitation from other environmental drivers of local adaptation to assess the impacts of a changing environment. Whether population persistence will be robust to these changes depends on both the relative importance of different drivers [25] and how these drivers co-vary in space and time.

We found that both temperature and precipitation affected performance, but detected local adaptation to temperature only (Fig. 3CD). Precipitation has been shown to be an important driver of local adaptation in some systems [26–29], but the absence of an overall affect in our analyses indicates that there is more to learn about how and when it is important. Adaptation to precipitation may be mediated in ways that our analyses did not capture. For example, populations may be adapted to the seasonal timing of precipitation [30] or how precipitation conditions amplify other abiotic or biotic pressures, such as competition [31] or wildfire [32]. Alternatively, if precipitation has been historically more variable than temperature, populations may have evolved broader tolerance to precipitation deviations than to temperature deviations [33, 34].

Local adaptation is a key element of biodiversity that often smooths out ecological dynamics in space [35]. However, our results show that as the climatic rug is being pulled out from under populations by anthropogenic warming, the stabilizing effect of local adaptation is being eroded. A critical next step for revealing which populations and species will be most vulnerable or resilient to ongoing rapid warming is to evaluate how local adaptation to temperature varies across species and geographic regions [36]. Additionally, further work is required to infer how changes in relative performance impact population persistence. Warming-driven changes in populations’ rank order of relative performance do not necessarily imply that populations are falling below a persistence threshold. They do, however, mean that populations are experiencing strong natural selection. Given sufficient genetic variation, such strong selection could drive rapid adaptation and favor the redistribution of lineages across the landscape. When natural migration is insufficient to keep pace with rates of climate change, our results can provide a starting point for determining which foreign populations will become best-matched to future conditions.

## Supporting information

Supplemental Area and Trace plots

## Acknowledgments

The authors would like to thank J. Elleouet, T. Richards, and C. Leven for assistance with initial data collection. D. Schluter, M. Whitlock, M. Pennell, M. Osmond, and D. Schemske provided critical feedback. We would like to thank all the authors that provided data that was not originally published in the form we required.

## Author contributions

MB and ALA conceived of the study. MB performed the literature search and managed data collection. MB, ALA, DEG, RMG, ALH, EJK, and KAT collected data. CRM advised on and prepared climate data. CDM developed theoretical and statistical models and conducted statistical analyses with assistance from MB. MB, CDM, and ALA wrote the main text of the manuscript with feedback and edits from all authors. CDM wrote the theory and statistical methods sections of the Materials and Methods with assistance from MB. MB and CDM made figures.

## Data and materials availability

All data and code required to recreate these analyses will be provided during review and will be archived in an public repository upon acceptance.

## Supplementary Materials for

**Other Supplementary Materials for this manuscript include the following:**

Data and code to fit models (compressed folder).

Trace and area plots of model estimates (separate pdf).

### 1 Material and methods: Data collection

#### 1.1 Literature search and criteria for inclusion

We searched for papers through 19 March 2017 using the Web of Science interface. We searched for: ((“reciprocal transplant*” OR “egg transfer experiment”) OR “local adaptation” AND ‘‘transplant*”) OR “provenance trial” OR “local maladapt*” OR ((“common garden*”) AND (“fitness” OR “surviv*” OR “reproduc*” OR “mortality” OR “intrinsic growth rate” OR “population growth rate”) AND (adapt*)) OR ((“common garden*” OR “reciprocal* transplant*” OR “transplant experiment” OR “assisted migration”) AND (temperature OR climat* OR latitud* OR elevation* OR altitud*) AND (“fitness” OR “surviv*” OR “reproduc*” OR “mortality” OR “intrinsic growth rate” OR “population growth rate” OR “establish*” OR “success*” OR “perform*”)) NOT invas* NOT marine NOT microb*). This returned 2111 results.

For inclusion in our analyses, we required that studies either moved ? 1 population to multiple locations, or moved ? 2 populations to a single location, and subsequently measured at least one component of fitness (germination, survival, reproduction, or a composite fitness metric such as population growth rate). We only included studies that moved populations at a geographic scale at which climatic differences could be detected between populations (>1 km or >200 m elevation between populations), therefore studies testing performance between fine-scale microhabitats were not included. We excluded studies in which the test environment was outside the species’ natural range or in a lab, growth chamber, or greenhouse. We also excluded studies of marine organisms because the climate variables used in our analyses are not likely to capture the climates experienced by these organisms [37]. We excluded studies that transplanted only hybrids or inbred lines and studies that performed reciprocal transplants between subspecies that are reproductively isolated.

Our initial search results were first screened based on their titles and abstracts. Studies that were obviously unsuitable were excluded, yielding 741 potential studies. After a more in-depth screening using the criteria described above, we retained 196 studies. Additionally, we checked the reference lists of other published meta-analyses and reviews [21, 38–41] and added any appropriate studies that were not returned by our initial search. We also added additional suitable studies that we encountered while gathering data. This resulted in the inclusion of 25 studies that were not returned in our initial database search. In total, we attempted to include 219 studies in our meta-analysis. Of the 219 studies, 72 were excluded once data collection had begun: 58 because we were unable to obtain adequate supplemental data from the authors and 14 because they were duplicates of data analyzed in other papers. This resulted in a final dataset of 147 studies of 164 species, the vast majority of which were plants (Fig. 2B, Table S1). Slightly different versions of this database have been used to examine the effects of biotic interactions on local adaptation [24] and how local adaptation and population quality vary across species geographic ranges [36].

When data were not published at the resolution that our study required, we requested data from authors. When data were presented in a figure, we used image analysis tools (WebPlotDigitizer, automeris.io/WebPlotDigitizer; GraphClick, www.arizona-software.ch/graphclick) to extract values as needed.

#### 1.2 Fitness data

Within a study, we collected fitness data for each unique combination of species, site, source, fitness metric, temporal replicate, and lifestage. A “site” refers to the test location (e.g., transplant garden), and a “source” refers to the population’s location of origin. For each data point, we categorized the source population as local or not local to the site. Sometimes this was obvious, for example in cases where a population was collected at or very near the test site. In some common garden studies (49 of 149) the authors did not explicitly designate a local population, so we used geographic or elevational proximity to assign one. When multiple fitness metrics were presented, we collected data from one representative metric in each of five possible categories (germination, recruitment (germination and survival combined), survival, reproduction, or a composite fitness metric). Reproductive estimates that account for mortality or failure to reproduce (i.e., population means that included zeros for non-reproductive or dead plants) were considered to be composite fitness estimates. When multiple measurements could be used for a single fitness metric (i.e., both flower counts and total seed weight were available for an estimate of reproduction) we selected the one that seemed most representative of fitness, at the discretion of the data collector. For any fitness estimates that were reported at multiple time points for the same cohort, we recorded a cumulative estimate (e.g., last reported survival as a proportion of starting sample size; summed reproduction from multiple seasons). When possible, we calculated cumulative fitness metrics from underlying components (e.g., composite fitness from germination x survival x reproduction). Sometimes transplants were conducted with starting material from different lifestages (i.e., seeds and seedlings), and we collected results from each type of planting. When studies included manipulative treatments (e.g., water addition or herbivore exclusion), we collected data from the treatment that most closely represented natural conditions.

In some cases, experiments were repeated in multiple years and we collected data from each temporal replicate. There was one exception to this: Wilczek et al. [12] was a uniquely large study, and a single site in that study with three temporal replicates contributed 690 lines of data. To avoid excessive influence of a single site on our results, we used only the fall temporal replicate of the Norwich site from this study. Across all studies, we obtained a total of 9414 fitness component measurements, each representing a unique combination of species, site, source, fitness metric, temporal replicate, and lifestage transplanted. These measurements included 1787 source populations at 541 sites.

We recorded latitude, longitude, and elevation for each source and site. We required precision of geographic coordinates to at least the nearest minute or 0.01 degrees in latitude/longitude; when data were presented at a coarser resolution, we contacted authors to request more precise coordinates. In some cases, latitudes and longitudes weren’t reported, but we were able to estimate locations from a map figure or other landmarks. In cases where elevations were not provided by authors, we used Google Earth and landmarks to estimate them.

#### 1.3 Climate data

We extracted time series of climatic records for each source and site using ClimateXY products ([42, 43]; ClimateNA for North America, ClimateSA for South America, ClimateEU for Europe, and ClimateAP for the Asia-Pacific region). These products downscale coarsely gridded (0.5°) historical monthly time series (CRU TS 3.22; [44]) using elevation-adjusted kilometer-scale climatological surfaces (PRISM for contiguous US, Western Canada, and Alaska; WorldClim2 elsewhere). For the 411 locations not covered by these databases, either because they were outside the spatial range or more recent than the latest available years in ClimateXY, we downscaled CRU TS 4.00 1901-2015 time series to the WorldClim2 climatology using the same change-factor (delta) method employed in the ClimateXY products.

We then used these data to estimate climatic deviations from normal conditions at a site (hereafter “climate deviations”) and climatic differences between a source population’s normal climate and the experimental conditions of test sites (hereafter “climate mismatch”). We estimated normal climates within a baseline period from 1951-1980. This time window was chosen because prior to 1950 the density of weather stations is too sparse for reliable estimations, while after 1980 the signal of climate change is already detectable [45].

We calculated deviations and mismatch specific to the duration and seasonal time window of data collection. For example, if a study recorded survival of seedlings transplanted in May of 2010 until August of 2010, we calculated deviations and mismatch based on the historical (1951-1980) monthly averages of May-August and the experimental monthly average of May 2010-August 2010. We expressed precipitation deviations and mismatches on the log scale so that they reflect proportional changes rather than absolute changes. We reasoned that a raw precipitation deviation of 100 mm has a different biological impact at a site with 100 mm mean annual precipitation versus one with 1000 mm.

#### 1.4 Covariates

The magnitude of local adaptation detected in transplant studies is likely to be correlated with the magnitude of environmental difference between the populations selected for the study. To account for this, we assumed that average geographic and elevation distances were rough proxies for environmental differences. We calculated a metric of study scale by putting geographic and elevation differences between sites into comparable units. We based this conversion on the amount of elevation or latitudinal distance required to generate an equivalent change in temperature. A change in elevation of 1000 m is, on average, equivalent to a 6°C change in temperature [46], while a change in latitude of 145 km is equivalent to a 1°C change in temperature [47]. Using these lapse rates as a conversion factor, we generated “composite” distances between sites and sources that account for both geographic distances and elevation differences between sites and sources.

### 2 Material and methods: Theory

Before using the data we compiled, we theoretically derive the expected fitness of foreign populations relative to that of local populations in a transplant experiment. More specifically, we derive this expectation as a function of the environmental distance (temperature or precipitation difference) between the foreign source population’s native site and the local experimental site. In Section 3 (Materials and Methods: Statistical analyses), we describe the statistical methods we used to fit this theory with data.

#### 2.1 Fitness along an environmental gradient

Many ecological and evolutionary models assume that fitness peaks under the conditions to which a population is best adapted and declines in environments that are further from the optimum. For a single environmental gradient *x*, we model absolute fitness (W) as a Gaussian function:

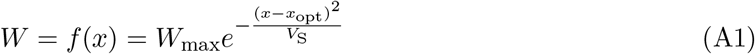

Gaussian fitness functions are commonly used in ecological and evolutionary models [48]. Here, *W*_max_ is the maximum absolute fitness in the optimum environment where x = *x*_opt_. *V*_s_ determines how quickly absolute fitness declines as *x* departs from *x*_opt_. This term is sometimes referred to as niche breadth in ecological models (e.g., [49]) and “selective variance” or “strength of stabilizing selection” in quantitative genetic models (e.g., [50]).

#### 2.2 Local and foreign fitness along an environmental gradient

Next, we extend the model to two populations, local and foreign, adapted to different environmental optima, *x*_opt,L_ and *x*_opt,F_, respectively. The absolute fitness of each population along environmental gradient *x* is:

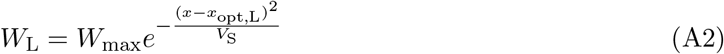

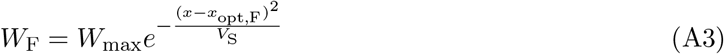

Note that we assume for simplicity that maximum fitness (*W*_max_) and niche breadth (*V*_s_) are the same for both local and foreign populations. However, their environmental optima (*x*_opt,L_ and *x*_opt,F_) can differ.

Here is a simple worked example that illustrates our underlying assumptions. Suppose that the average historical environment of the local population is 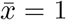 and that of the foreign population is 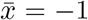. Further, suppose that both are perfectly adapted to these environments, so *x*_opt,L_ = 1 and *x*_opt,F_ = −1. During a transplant experiment, the site environment is *x* = 0.9, which is very close to the historical average for the local population. If *W*_max_ = 1 and V_S_ = 1, then *W*_L_ = 0.99 and *W*_F_ = 0.03. The local population has much higher fitness than the foreign population because of local adaptation to its environment.

#### 2.3 The strength of local adaptation

In general, populations are considered to be locally adapted if they have higher fitness than a foreign population transplanted to their native environment, as shown in the example above. But the strength of local adaptation (how much higher local fitness is than foreign) varies considerably [21]. We define the strength of local adaptation as the log-ratio of *W*_L_ to *W*_F_. For brevity, we refer to this quantity as ω. Along a single environmental gradient, the model above predicts that the strength of local adaptation is:

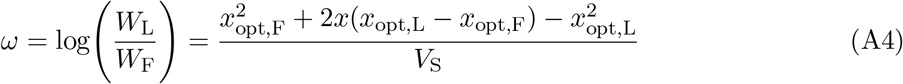

This formulation has several desirable properties. First, populations are locally adapted when *W*_L_ > *W*_F_ (which means that *ω* > 0). Second, the strength of local adaptation does not depend on the absolute environment *x*, but on the position of *x* relative to the difference in population optima. For example, *ω* = 3.6 when *x* = 0.9, *x*_opt,L_ = 1, *x*_opt,F_ = −1, and *V*_S_ = 1. Symmetrically, *ω* = 3.6 also when *x* = −0.9, *x*_opt,L_ = −1, *x*_opt,F_ = 1, and *V*_S_ = 1. The switch from local adaptation (*ω* > 0) to foreign advantage (*ω* < 0) occurs when *x* is closer to *x*_opt,F_ than it is to *x*_opt,L_. Finally, this formulation shows that, all else being equal, greater difference between local and foreign optima 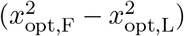 and narrower niche breadth (smaller *V*_S_) increase the strength of local adaptation (or foreign advantage, depending on whether the experimental environment favors the local or the foreign population).

#### 2.4 The effect of climate mismatch on relative fitness of local and foreign populations

In this section, we apply the theory introduced above for two populations (local and foreign) to derive expectations for how climate mismatch affects the relative fitness of local and foreign populations. Climate mismatch for population *i* at site *j* (*m_ij_*) is defined as the difference between the experimental conditions at site *j* (*x_j_*) and the average climate of population *i* in its home site 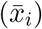:

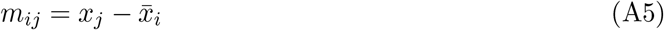

Relative fitness of population *i* at site *j* is the absolute fitness of population *i* at site *j* divided by the average absolute fitness in site *j*:

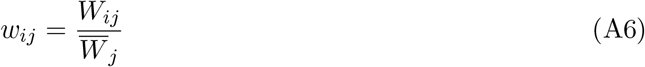

As above, we model population fitness with a Gaussian function (Equation A1) assuming that optimal environment is population-specific, but that maximum absolute fitness (*W*_max_) and niche breadth (*V_S_*) are the same within a species in a given experiment.

The average fitness in an experiment at site *j* with environment *x_j_* is the mean fitness of all *n* foreign populations:

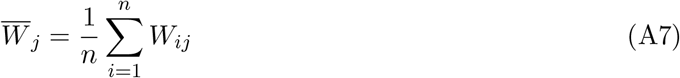

We estimate average fitness from foreign populations only to control for differing numbers of foreign populations in each experiment. When populations are locally adapted, including the local population in 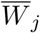 will cause the relative fitness of local populations to appear lower in smaller experiments because local fitness contributes more to the average fitness. Assuming that foreign populations are chosen randomly with respect to experiment size, the number of foreign populations in an experiment will not bias the estimate of 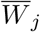, though the variance will be larger among smaller experiments.

The relative fitness of the local population at site *j* (*w_L,j_*) is its absolute fitness (*W_L,j_*) divided the average fitness of foreign populations:

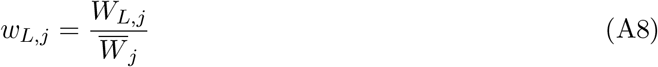

Substituting Equation A2 in the numerator, we obtain:

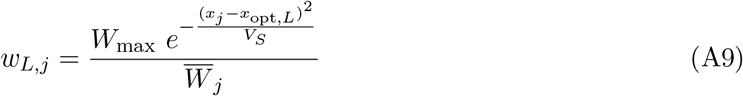

For one of the foreign populations, substituting Equation A3 yields the analogous equation:

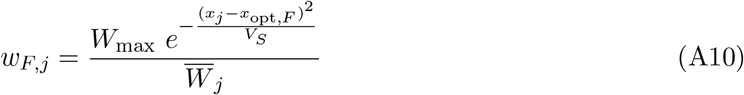

The relative fitness of the local population will be greater than a foreign population as long as the experimental environment is closer to the local than the foreign optimum, *x_j_* — *x*_opt,*L*_ < *x_j_* – *x*_opt,*F*_. Even when the experimental environment is not in between to the two populations’ optima, it will still be nearer to one than the other. Note that Wmax cancels out of both equations because it is a scalar in the denominator as well.

Climate mismatches should decrease relative fitness of local and foreign populations in the same manner if the expected mean fitness of foreign populations 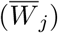 is constant for all values of climate mismatch. This assumption is valid as long as a change in climate mismatch for a focal population (local or foreign) increases the fitness of some foreign populations as much as it decreases fitness of other foreign populations, keeping 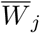 unchanged.

If populations are locally adapted to their environment, then the optimal environment is close to the average environment, 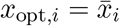. Substituting this into Equation A5, we can equate climate mismatches to deviations of the experimental environment from the optimal environment, *m_ij_* = *x_j_* – *x*_opt,*i*_. Therefore a climate mismatch will lower absolute fitness, all else being equal.

If both the local population and a given foreign population have the same optimal environment, then this model predicts they have equivalent relative fitness at a given climate mismatch. However, since local populations are likely adapted to multiple environmental factors, we expect that local populations will have higher relative fitness than foreign populations even with the same amount of climate mismatch. In Section 3 (Materials and Methods: Statistical analyses), we account for local adaptation to other unmeasured environmental factors.

#### 2.5 Climate deviations increase the variation in local adaptation

We anticipate that climate deviations will not, on average, favor or disfavor local populations, but will increase the variation in experimental outcomes. We define deviations as the difference between the environment at time i and the average environment at a site:

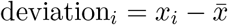

We also assume that the local population is adapted to the average environment at its home site:

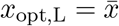

Hence, deviation, *i*. = *x_i_* – *x*_opt,L_. We obtain *ω* at a given time as a function of deviations by substituting *x* = *x_i_*, = deviation_*i*_, + *x*_opt,L_ in Equation A4:

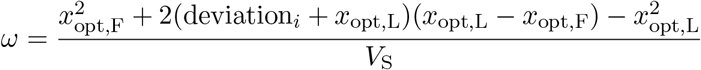

This expression simplifies to:

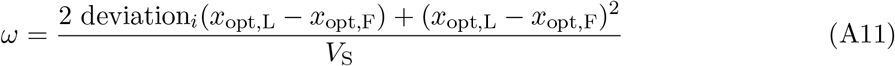

In an “average” time period, when deviation_*i*_, = 0, the strength of local adaptation is the squared difference in environmental optima divided by the niche breadth, *ω* = (*x*_opt,L_ – *x*_opt,F_)^2^/V_S_. Deviations strengthen local adaptation when they reinforce the difference in environmental optima, deviation_*i*_, (*x*_opt,L_ – *x*_opt,F_) > 0; deviations weaken local adaptation when they counteract the difference in optima, deviation_*i*_, (*x*_opt,L_ – *x*_opt,F_) < 0:

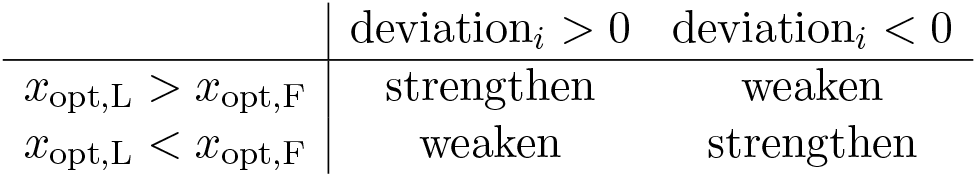

|  | deviation<sub>i</sub> > 0 | deviation<sub>i</sub> < 0 |
| --- | --- | --- |
| $x_{\text{opt,L}} > x_{\text{opt,F}}$ | strengthen | weaken |
| $x_{\text{opt,L}} < x_{\text{opt,F}}$ | weaken | strengthen |

The net result is that climate deviations increase the variance in experimental outcomes. For example, as warm deviations increase in frequency, local populations adapted to warmer climates than foreign populations experience an even stronger local advantage. Conversely, warm deviations will cause foreign populations adapted to warmer climates to have higher fitness than local populations.

Here we illustrate this verbal argument mathematically. For simplicity, we assume that niche breadth is the same for local and foreign populations. We treat both deviations and the difference between local and foreign optima (*d*_clim_ = *x*_opt,L_ — *x*_opt,F_) as random variables. Prior to climate change, we assume that deviations are equally likely to be warm or cold, wet or dry (*μ*_dev;*t*0_ = 0) with some variance (*V*_dev_). Likewise, differences in climatic optima are equally likely to be positive or negative (*μ*_diff_ = 0) with some variance (*V*_diff_). With climate change, there is a directional shift (e.g. warmer average temperature), so *μ*_dev,*t*1_ = 0, but the variation around the mean remains the same as in the past. Although we assume it is constant for this derivation, if variation has increased because of global climate change, this would further increase variance in experimental outcomes.

To keep the notation simpler, let *x* = 2 deviation and *y* = *d*_clim_. Substituting *x* and *y* into Equation A11 yields:

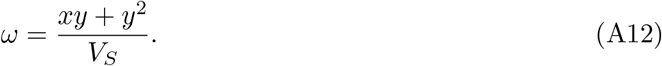

Based on standard random variable algebra, the variance in *ω*, denoted *V_ω_*, is:

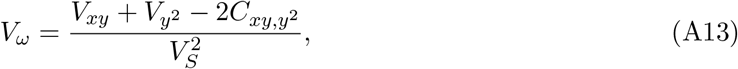

where *V_xy_* is the variance of *xy*, *V*_*y*^2^_ is the variance of *y*^2^, and *C*_*xy,y*^2^_ is the covariance between *xy* and *y*^2^. We can further decompose *V_xy_* and *V*_*y*^2^_. The variance of the product of two random variables (*V_xy_*) is defined as:

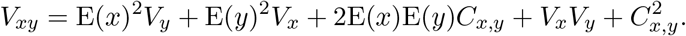

E(*x*) is the expected value of *x*, i.e. the average value of a climate deviation. If we assume that there is no correlation between climate deviations and the difference between local and foreign optima, then *C_x,y_* = 0. The above equation simplifies to:

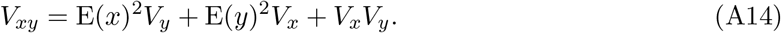

The variance in the square of population climatic differences (*V*_*y*^2^_) is:

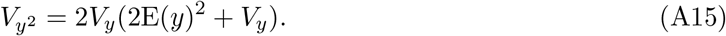

Now, substituting Equations A14 and A15 into Equation A13, we obtain an expression for the variance *ω* as function of the mean and variance in climate deviations and the difference in climatic optima:

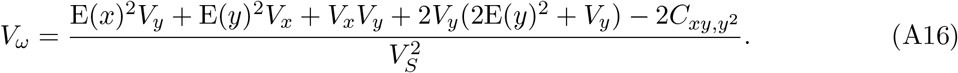

This equation is complex, but it reveals some key insights. First of all, the variance in local adaptation increases when there is greater temporal variation in climate (increased *V_x_*, the variance in deviations) and greater variation in the difference between local and foreign optima (*V_y_*). If either of these variances change over time because of climate change and/or experimental practices, it will affect the variance in local adaptation. For example, greater interannual variation in temperature would increase the variance in local adaptation. We would see a similar effect if scientists were comparing local fitness to a wider variety of foreign populations. The variance in local adaptation will decrease if the niche breadth is wider.

The variance in local adaptation will also increase with a directional shift in climate, even if climatic variability remains the same. Directional climate change shifts the expected value of deviations, E(*x*), from 0 to some new value (e.g. 1° warmer). All else being equal, the variance in *ω* increases with the square of E(*x*). Therefore, if climate deviations are changing directionally (e.g., more warm deviations and fewer cold deviations), experimental outcomes should become more variable after accounting for other factors, such as niche breadth and differences in climatic optima, that determine the strength of local adaptation.

### 3 Materials and methods: Statistical analyses

We first examined whether we observe larger climate deviations over time, consistent with global climate trends, in our database of transplant experiments (Model A). Next, we tested whether the fitness of populations (relative to the average foreign fitness in a site) is affected by experimental conditions and deviations from their climates of origin (Model B).

If climate deviations are becoming more frequent and populations are locally adapted to climate, then it follows that our measurements of the strength of local adaptation may be changing. Our theoretical model predicts that climate deviations strengthen local adaptation when the local population is better adapted to anomalous conditions than the foreign population; for example, warm deviations will strengthen local adaptation when the local population is adapted to warmer conditions than the foreign population. Conversely, warm deviations are predicted to weaken or even overturn local adaptation when the local population is adapted to cooler climates than the foreign population. We tested whether deviations have these predicted effects on local adaptation and whether these effects are leading to increased variance in local adaptation over time in Model C.

We fit these models in Stan [51]; all Stan code and data required to fit these models has been submitted for review and will be published in an appropriate repository upon acceptance. We provide further information on model fitting and posterior predictive checks in Section 3.6.

When estimating effect sizes from many different datasets, it is best practice to account for variance in estimates, as studies may differ in their sample sizes and power to detect significant effects. However, our data set includes data on many different scales (i.e., survival proportions vs. counts of seed numbers), which means that uncertainty is not reported using a consistent metric across the original studies. In some cases, uncertainty in estimates is not reported at all. Rather than lose data by omitting those that did not report variance, we accounted for other factors that might affect the magnitude local adaptation detected, such as the number of populations being compared and the geographic extent from which populations are drawn.

#### 3.1 Model A: Are climate deviations becoming more frequent?

We tested whether temperature and/or precipitation deviations are becoming more frequent using a Bayesian mixed effects model. Table A2 summarizes model parameters. Temperature deviations were expressed as °C_experiment_ – °C_site normal_. We calculated these deviations over each temporal window in which fitness was measured at each site, for a total of 809 deviation observations. For each site, we treated mid-year (average of start and end years for studies that spanned multiple years) of the experiment as a fixed effect regressed against temperature deviation. If warm deviations are becoming more common, then the coefficient *β*_year_,A should be significantly greater than zero. We rescaled year by subtracting 1975 (the earliest mid-year in our dataset) so that the y-intercept (*β*_0,A_) is the average temperature deviation in 1975. We also included site as a random effect. We also tested whether the variance in temperature deviations is increasing through time by estimating the effect of year (scaled) on the residual scale term, assuming residuals are normally distributed. We repeated this process with precipitation deviations, which were expressed as log_10_mm_experiment_ – log_10_mm_site normal_. Because we did not have an *a priori* expectation about whether dry deviations or wet deviations might be becoming more frequent among the sites in our data set, we also tested whether the absolute values of precipitation deviations were changing over time. Parameter estimates are presented in Table S2.

**Table A1.** Glossary of symbols for Theory section

| Symbol | Description |
| --- | --- |
| $\text{deviation}_i$ | Environmental deviation at time $i$ |
| $x$ | Environmental gradient such as average temperature or precipitation |
| $x_{\text{opt}}$ | Optimal environment where absolute fitness is maximized |
| $x_{\text{opt,L}}$ | Optimal environment ($x_{\text{opt}}$) for **local** population |
| $x_{\text{opt,F}}$ | Optimal environment ($x_{\text{opt}}$) for **foreign** population |
| $V_s$ | Niche breadth or strength of stabilizing selection |
| $W$ | Absolute fitness |
| $W_{\text{max}}$ | Maximum absolute fitness when $x = x_{\text{opt}}$ |
| $W_L$ | Absolute fitness of **local** population |
| $W_F$ | Absolute fitness of **foreign** population |

**Table A2.** Model A parameters and predictions. Each parameter has a mathematical Symbol used in the text, a Stan character string used in the model code, a brief Description of the parameters, and associated *a priori* Predictions where applicable. Parameter estimates from fitted models are presented in Table S2 and in Fig. 2CD.

| Symbol | Stan | Description | Prediction<br>(where applicable) |
| --- | --- | --- | --- |
| Regression on climate deviations |  |  |  |
| $\beta_{0,T,A}$ | b0_t_A | Average temperature deviation in year 0 (= 1975) | |
| $\beta_{T,\text{year},A}$ | b_t_A_year | Effect of year on average temperature deviation | $\hat{\beta}_{T,\text{year},A} > 0$ |
| $\beta_{T,\text{site},A_i}$ | b_t_A_site[i] | Effect of site $i$ on average temperature deviation | |
| $\sigma_{T,\text{site},A}$ | sigma_t_A_site | Standard deviation of $\beta_{T,\text{site}_i,A}$ | |
| $\beta_{0,P,A}$ | b0_p_A | Average precipitation deviation in year 0 (= 1975) | |
| $\beta_{P,\text{year},A}$ | b_p_A_year | Effect of year on average precipitation deviation | |
| $\beta_{P,\text{site},A_i}$ | b_p_A_site[i] | Effect of site $i$ on average precipitation deviation | |
| $\sigma_{P,\text{site},A}$ | sigma_p_A_site | Standard deviation of $\beta_{P,\text{site}_i,A}$ | |
| $\beta_{0,AP,A}$ | b0_ap_A | Average absolute precipitation deviation in year 0 (= 1975) | |
| $\beta_{P,\text{year},A}$ | b_p_A_year | Effect of year on average absolute precipitation deviation | |
| $\beta_{P,\text{site},A_i}$ | b_p_A_site[i] | Effect of site $i$ on average absolute precipitation deviation | |
| $\sigma_{AP,\text{site},A}$ | sigma_ap_A_site | Standard deviation of $\beta_{AP,\text{site}_i,A}$ | |
| Regression on $\sigma_{\epsilon,A}$ | | | |
| $\beta_{\epsilon,T,A,\text{year}}$ | b_t_A_resid_year | Effect of study year on $\log(\sigma_{\epsilon,T,A,0})$ | |
| $\log(\sigma_{\epsilon,T,A,0})$ | log_sigma0_t_A_resid | Standard deviation of temperature deviations (log-scale) in study year 0 (= 1975) | |
| $\beta_{\epsilon,P,A,\text{year}}$ | b_p_A_resid_year | Effect of study year on $\log(\sigma_{\epsilon,P,A,0})$ | |
| $\log(\sigma_{\epsilon,P,A,0})$ | log_sigma0_p_A_resid | Standard deviation of precipitation deviations (log-scale) in study year 0 (= 1975) | |
| $\beta_{\epsilon,AP,A,\text{year}}$ | b_ap_A_resid_year | Effect of study year on $\log(\sigma_{\epsilon,AP,A,0})$ | |
| $\log(\sigma_{\epsilon,AP,A,0})$ | log_sigma0_ap_A_resid | Standard deviation of absolute precipitation deviations (log-scale) in study year 0 (= 1975) | |

#### 3.2 Model B: Do site conditions and climate deviations affect relative fitness?

We first tested whether relative fitness is lower in sites that are hotter, cooler, wetter, or drier than other sites in an experiment, regardless of how conditions match the source climates of populations (Model B1, Fig. 1A). This could occur if performance responds plastically to weather conditions in a similar manner across populations, instead of or in addition to locally adapted responses. To estimate this, we centered the temperature and precipitation conditions at each site j around the average within each study (*T_j_*, *P_j_*). Site temperature conditions were expressed in °C difference from the experiment average and precipitation conditions as the difference between log_10_(experiment average precipitation) and log_10_(site precipitation).

We treated centered temperature and precipitation conditions as fixed effects regressed against relative fitness. We calculated relative fitness for each population *i* at site *j* (*w_ij_*) by dividing the fitness of that population by the average fitness of all foreign populations at all sites (within each temporal window and for each fitness metric). This put diverse fitness metrics (e.g., proportion survival, seed number) on a common scale. Our results should be interpreted accordingly; we cannot make inferences about absolute fitness or population growth, only relative performance. We estimated model parameters using quadratic regression of site temperature and precipitation against log-transformed relative fitness (log *w_ij_*) to normalize the residual variance. This model allowed us to evaluate the shape of the relationship between site temperature/precipitation conditions and fitness.

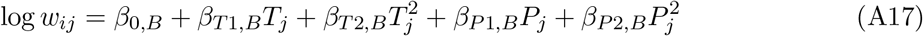

Table A3 describes model parameters. We included a covariate of the composite distance over which a population was moved, which may coarsely account for the displacement a population experiences along ecological axes. We also included a covariate of whether a population was local or foreign in a given site, as local adaptation to any aspect of the environment might result in populations performing better in their home site. We included a random intercept for each combination of study and taxon.

**Table A3.** Model B1 parameters and predictions. Each parameter has a mathematical Symbol used in the text, a Stan character string used in the model code, a brief Description of the parameters, and associated *a priori* Predictions where applicable. Parameter estimates from fitted models are presented in Table S3 and Fig. 3AB.

| Symbol | Stan | Description | Prediction<br>(where<br>applicable) |
| --- | --- | --- | --- |
| Regression on $\log(w_{ij})$ | | | |
| $\beta_{0,B}$ | b0_B | Average $\log(w_{ij})$ under average temperature and precipitation conditions | $\beta_{0,B} = 0$ |
| $\beta_{T1,B}$ | b1_ta_B | Linear effect of $T$ on $\log(w_{ij})$ | |
| $\beta_{T2,B}$ | b2_ta_B | Quadratic effect of $T$ on $\log(w_{ij})$ | |
| $\beta_{P1,B}$ | b1_pa_B | Linear effect of $P$ on $\log(w_{ij})$ | |
| $\beta_{P2,B}$ | b2_pa_B | Quadratic effect of $P$ on $\log(w_{ij})$ | |
| $\beta_{\text{local},B}$ | b_local_B | Effect of being local | $\hat{\beta}_{\text{local},B} > 0$ |
| $\beta_{\text{dist},B}$ | b_dist_B | Effect of composite distance moved | $\hat{\beta}_{\text{dist},B} < 0$ |
| $\beta_{\text{study\_taxon},B_i}$ | b_study_taxon_B[k] | Effect of study-taxon $k$ on $\log(w_{ij})$ | |
| $\sigma_{\text{study\_taxon},B}$ | sigma_B_study_taxon | Standard deviation of $\beta_{\text{study\_taxon},B_i}$ | |
| Regression on $\sigma_{\epsilon,B}$ | | | |
| $\beta_{\epsilon,B,n}$ | b_B_n | Effect of $\sqrt{n}$ on $\log(\sigma_{\epsilon,B})$ | |
| $\log(\sigma_{\epsilon,B,0})$ | log_sigma0_B_resid | Standard deviation (log-scale) of residual $\log(w_{ij})$ when $n = 0$ | |

When *w_ij_* for a given population was measured as 0 (8.2% of observations), our calculation of log *w_ij_* is −∞. We censored these observations as described for Model C (see section 3.3.6 below) using the minimum non-zero relative fitness for a given combination of study, taxon, and fitness type as a lower censor. We excluded 6 observations of performance for which fitness was measured as 0 at all sites, and 288 observations in which populations were transplanted to only one site, resulting in 9120 observations used to fit this model.

We assumed that residual variation is normally distributed but decreases linearly with the square root of *n*, the number of populations used to calculate average fitness. If *n* is high and foreign populations are sampled randomly, the variability in *w_ij_* is primarily due to absolute fitness of the focal population *W_ij_*; when *n* is low, sampling variance in the foreign populations also contributes, but this additional variance should decrease with the square-root of sample size. We estimated this parameter (*β_n,σ,B_*) on a log-link scale and predicted it should be less than 0, meaning that the residual variance decreases in proportion to 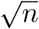. Parameter estimates are presented in Table S3.

We next tested whether a population’s fitness declines as they experience more extreme temperature or precipitation mismatch (Model B2). Our transplant dataset allows us to examine a broad range of climate mismatch between experimental conditions at a site and the climate a population has historically experienced. This is because transplanted populations experience deviations caused by interannual variability within a site but also climatic mismatch introduced by moving across space (Fig. 1C). We calculated overall climatic mismatch for each population at each site (see Equation A5): temperature mismatch for each population-site combination *ij* (*m_T,ij_*) was expressed as the difference between temperature during the experiment and average temperature at the source (in °C) and precipitation mismatch for each population-site combination *ij* (*m_p,ij_*) was expressed as the difference between log_10_ (precipitation during the experiment) and log_10_(average precipitation at the source). For local populations, this mismatch is simply the result of interannual variability in their home sites; for foreign populations it incorporates both experimental conditions and spatial differences in climate.

For each site, we treated temperature and precipitation mismatch as fixed effects regressed against relative fitness. We calculated relative fitness for each population *i* at site *j* (*w_ij_*) by dividing the fitness of each population by the average fitness of all foreign populations (see Section 2.4, Equation A6) at that site within each temporal window, for each fitness metric. In contrast to our relativization for Model B1, here we relativize within sites, which removes the effect of differences in site quality. Following the assumptions in Section 2 (Materials and Methods: Theory), we predicted that relative fitness would peak when mismatch is near zero and decline as mismatch becomes larger.

Based on the theory developed in Section 2.4 (Materials and Methods: Theory), we predicted a Gaussian relationship between climate mismatch for both local and foreign populations (Equation A9–A10). We estimated model parameters using quadratic regression of temperature and precipitation mismatch against log-transformed relative fitness (log *w_ij_*) to normalize the residual variance. Exponentiating a quadratic equation yields a Gaussian curve when the coefficient on the squared term is negative.

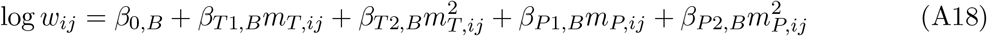

Table A4 describes model parameters. First, this model allowed us to evaluate whether a hump-shaped relationship between climate mismatch and fitness was present in our data if *β*_*T*2,*B*_ < 0 or *β*_*P*2,*B*_ < 0. We predicted that linear parameters (*β*_*T*1,*B*_ and *β*_*P*1,*B*_) should be approximately 0 if the optimal climate mismatch is 0. If the linear parameters *β*_*T*1,*B*_ or *β*_*P*1,*B*_ = 0, this either indicates that the fitness response peaks at a non-zero climate mismatch (if the quadratic term is significant) or an exponential relationship between relative fitness and climate (if the quadratic term is not significant). Temperature and precipitation mismatch were not highly correlated in our dataset (*r* = 0.034, correlation between absolute value of the two mismatch types: *r* = 0.23).

We also predicted that the average relative fitness of foreign populations should be 1 when climate mismatch is 0 (*β*_0,*B*_ ≈ 0 because log(1) = 0), but that local populations would have higher relative fitness than foreign populations for a given degree of climate mismatch. We tested this latter prediction by including a fixed effect of local origin (*β*_local,*B*_). When *w_ij_* for a given population was measured as 0 (8.2% of observations), our calculation of log *w_ij_* is –∞. We censored these observations as described for Model C (see section 3.3.6 below) using the minimum non-zero relative fitness for a given combination of study, taxon, and fitness type as a lower censor. Fitness type and functional group did not significantly alter the relationship between mismatch and relative fitness (results not shown), so these were not included here. We excluded 58 observations for which there was no non-zero fitness estimate at the site, resulting in a dataset of 9356 observations. As described for Model B1, we assumed that residual variation is normally distributed but decreases linearly with the square root of the number of foreign populations used to calculate average fitness, and included terms to account for this.

When the quadratic term was significantly negative for temperature or precipitation mismatch, we estimated from these quadratic fits:

- The optimal temperature and precipitation mismatch (the location of the peak of the curves) as –*β*_*T*1,*B*^/2*β*^*T*2,*B*_ and –*β*_*P*1,*B*_/2*β*_*P*2,*B*_;
- The Gaussian variance in response to temperature and precipitation (the breadth of the curves) as – 1/*β*_*T*2,*B*_ and – 1/*β*_*P*2,*B*_.

#### 3.3 Model C: Are populations locally adapted to climate?

Next, we hypothesized that populations are adapted to the historic climate at their site of origin and, hence, maladapted to the historic climate at foreign sites. Based on this hypothesis, we developed a theoretical model of local adaptation to derive quantitative predictions to test with our dataset (see Section 2, Materials and Methods: Theory). This model predicts that the strength of local adaptation (*ω*) should be a function of the experimental site environment (*x*), the environmental optima of local and foreign populations (*x*_opt,L_ and *x*_opt,F_), and the niche breadth (*V*_s_):

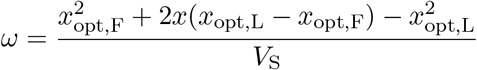

We fit this model using only data from transplant experiments with designs that allowed for the contrast of a local population and at least one foreign population in the same site. There is no information about the fitness ratio when both local and foreign fitness (*W*_L_ and *W*_F_) are 0, so we removed these observations. After applying these filters, our dataset contained 5705 local-foreign contrasts.

The simple model above does not account for local adaptation to unmeasured environmental axes, differences in experimental design, or variation in niche breadth across taxa. In later sections, we explain how we incorporated these additional factors into a Bayesian linear mixed effects model.

##### 3.3.1 Relative experimental environment (EE_rel_)

Equation A4 shows that *ω* is proportional to the quantity 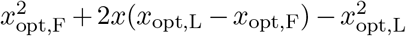. This quantity scales the site environment during the experiment relative to the optimal environments for each of the populations. For brevity, we refer to this quantity as the <u>Rel</u>ative <u>E</u>xperimental <u>E</u>nvironment (EE_rel_):

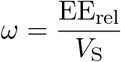

If populations are locally adapted, then the optimal environment should be close to the historical average environment at that site. Hence, we estimated *x*_opt,L_ and *x*_opt,F_ for temperature or precipitation as the average historical values in the respective home sites of the populations. We also obtained data for the site climate (*x*) during the experiment (see Section 1.3 for further detail). When the experimental environment favors the local population over the foreign population, EE_rel_ will be large. This doesn’t necessarily mean that the experimental environment is close to the optimal environment for the local population, only that the experimental environment is closer to the local optimum than it is to the foreign optimum. We estimated the average niche breadth from the inverse of the slope of EE_rel_ against *ω*:

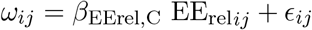

*β*_EErel,C_ is therefore an estimate of 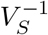. The variables *ω_i,j_*, EE_rel*ij*_, and *ϵ_i,j_* are the relative experimental environment, log-fitness ratio, and residuals of the *j*th observation from the *i*th study, site, taxon, and fitness type combination (see explanation below). If populations are locally adapted such that the population that is best matched to the environment performs better, then our niche breadth term should be positive, meaning the slope (*β*_EErel,C_) should also be positive. Estimated slopes near or less than 0 would indicate no adaptation or maladaptation, respectively. For fitting purposes, the slope parameter had better statistical properties, but we transformed posterior estimates to niche breadth, which is more biologically meaningful.

##### 3.3.2 Local adaptation to other environmental factors

In this study, we focused on climate because human activity has increased average temperature globally and altered precipitation regimes. However, populations could simultaneously be locally adapted to many other biotic and abiotic factors. If temperature or precipitation were strongly correlated with these other unmeasured factors, this could make local adaptation to climate appear much stronger than it is. We accounted for local adaptation to unmeasured environmental factors in three ways:

- An intercept term (*β*_0,C_) to estimate average local adaptation to unmeasured factors;
- A composite distance metric combining geographic and elevational distance between a population’s original location and the experimental site (*β*_dist,C_), assuming that this may be a proxy for overall environmental distance (see Section 1.4 for calculation of this metric).

After incorporating these effects, the statistical model becomes:

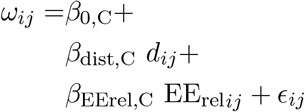

*d_ij_* is the geographic distance between the experimental site and focal population. If there is local adaptation to unmeasured factors, then 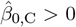. If geographic distance is a good proxy for differences in these selection pressures, then 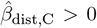. A negative value for 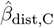 would indicate that greater distance weakens local adaptation, after accounting temperature or precipitation differences. This could occur if, for example, more distant populations are less susceptible to local pathogens or herbivores.

##### 3.3.3 Variation in niche breath

Niche breadth, which measures the sensitivity of fitness to deviations from the optimal climate, likely varies among taxa and populations. To account for this additional variation, we included a random effect of each combination of study, taxon, source population, and fitness type on the slope:

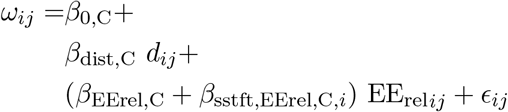

##### 3.3.4 Experimental differences in measuring fitness

As described in Section 1.2, the studies in our database measured a variety of fitness metrics. Studies that measured a single fitness component may have captured less of the total fitness difference than studies that measured multiple fitness components. The overall intercept is the expected ω for the composite fitness type including all components and the effect of less-complete fitness types were estimated as fixed effects.

With these effects, the final statistical model becomes:

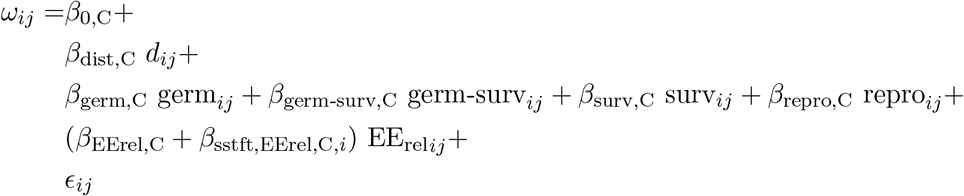

##### 3.3.5 Is the variance in local adaptation changing over time?

Our theoretical model predicts that if deviations are changing directionally over time (e.g. warmer or drier, see Section 3.1), then the variance in the strength of local adaptation should be increasing through time (see Section 2.5). The effect is caused by the fact that more extreme deviations can both 1) overturn local adaptation, leading to foreign advantage where it otherwise would not have occurred, and 2) reinforce local advantage, leading to stronger local adaptation than would have otherwise occurred. We tested whether overall variance in local adaptation increased through time by regressing study year against the scale parameter of residuals (*ϵ_ij_*), using a log link because the variance must be positive. We modeled *ϵ_ij_* using a Student *t* distribution, a robust regression approach which is less influenced by extreme data points [52]. As the “degrees of freedom” parameter *ν* approaches ∞, the t distribution converges to a normal distribution with mean *μ* and standard deviation *σ*. We modeled the effect of time on the scale parameter to test if the variance changes over time:

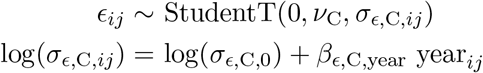

We predicted that the slope of study year against log(*σ_ϵ,C,ij_*) will be positive.

Because climate deviations can both reinforce and counteract historical climatic differences between populations, we did not predict that the average strength of local adaptation would change over time, only the variance. We tested whether the average strength of local adaptation changed through time by including a fixed effect of study year (*β*_C;year_), which we did not expect to be significantly different from 0. With these parameters, the final model becomes:

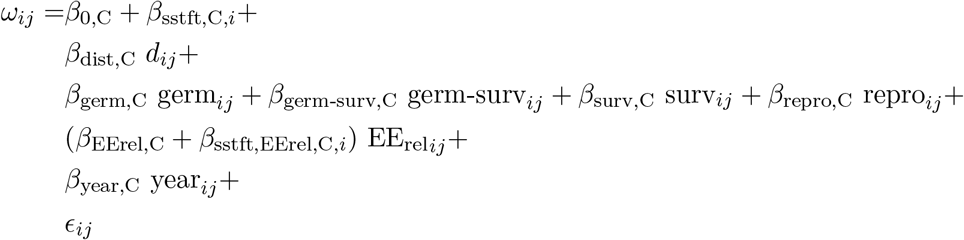

Increased variance in deviations, increased variance in EE_rel_, or decreased variance in niche breadth (*V_S_*) over time should also cause the variation in ω to increase (see Equation A16). Statistical Model A (Section 3.1) tests whether the variance in deviations is increasing; statistical Model C accounts for the latter effects. For example, if the variation in EE_rel_ increased through time because scientists were choosing foreign populations with a wider variety of temperature optima, this variation would be explained by the EE_rel_ coefficient. Random effects also allow us to account for variance in niche breadth among taxa. Hence, if the variance in local adaptation increases after accounting for these factors, it strongly implies that unmeasured changes in climate and/or experimental practice are responsible.

##### 3.3.6 Censoring nonfinite observations

Some observations (6.8%) of log-relative fitness were not finite because either the local or foreign fitness were measured as 0 (log(0/*x*) = −∞; log(*x*/0) = ∞). When the expected absolute fitness is very low, fitness will often be 0 because of finite sample sizes. For example, if the expected number of fruits in a sample is Poisson-distributed with a mean of 2, there is a ~13.5% chance of measuring 0 fruits. Values of −/ + ∞ have 0 likelihood in the statistical model and cannot be included. However, excluding these instances is not ideal, since they likely represent cases of very strong local adaptation or foreign advantage.

Therefore, we treated nonfinite observations of *ω* as censored because researchers were unable to observe below a certain value (the censor). For example, assume the lowest observable fitness value is *W*_min_. If the local fitness is *W*_L_, then the upper (a.k.a. right) censor is 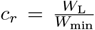. If *W*_L_ = 0, then lower (a.k.a. left) censor is 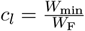.

We estimated *W*_min_ for every combination of study, site, taxon, and fitness type as the minimum non-zero fitness value observed in that experiment. This can be considered an upper (lower) bound for *c_r_* (*c_l_*) since there are likely non-zero fitness values that could have been measured but were not. The site-specific minimum fitness for site *j* is denoted *W*_min,j_. When *W*_F_ = 0 and *W*_L_ > *W*_min,*j*_, we right-censored the data at 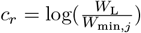. When *W*_L_ = 0 and *W*_F_ > *W*_min,*j*_), we left-censored the data at 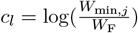.

Uncensored data (*ω* > *c_l_*, or *ω* < *c_r_*) have a likelihood of 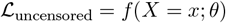 where *f* (*X; θ*) is the probability density function with parameters *θ*. To calculate the likelihood of censored data, we use the associated cumulative density function *F*(*x; θ*). The likelihood of censored data is the cumulative probability density from −∞ to *c_l_*, or *c_r_* to −∞ for left- and right-censored observations, respectively:

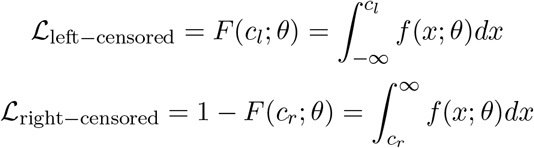

#### 3.4 Extending Model C: Have temperature deviations increased the variance in local adaptation?

As we report in the main text, warm temperature, but not precipitation, deviations are increasing (Fig. 2CD) and local adaptation is becoming more variable (Fig. 4B). We explore the effects of climate on the magnitude of local advantage in three additional analyses. First, we examine whether warm climate deviations reinforce and counteract local adaptation in predictable ways depending on whether the foreign population is expected to be better or worse suited to a warm climate. Second, we modified statistical Model C (Section 3.3) to quantify how much temperature deviations *per se* are contributing to increased variance in local adaptation. This approach asks whether increased variance associated with temperature deviations contributes significantly to the overall increase in variance.

In the first approach, we extracted two subsets of observations from our dataset for Model C: one in which sites resembled their historic normals (< 0.5° absolute temperature deviations, 1820 observations) and the other in which sites had > 2° warm temperature deviations (515 observations). We then classified each contrast as either one in which we expect local adaptation to be reinforced (when the foreign population is from a cooler location than the local population, 1367 observations) or one in which we expect local adaptation to be counteracted (when the foreign population is from a warmer location than the local population, 968 observations). We then fit a model that estimated the effect of site conditions (normal or warm), foreign temperature origin (warmer or cooler) and the interaction between these two factors on the magnitude of local adaptation. We included covariates of composite distance and fitness component, and fit the model using the robust regression approach described for Model C.

In a second approach, we estimated the effect of study year on variance in the magnitude of local advantage *after* removing the model-predicted effect of temperature and precipitation deviations. For each observation, the average climate deviation in that study year, estimated from model A, was used to modify the observed value of *ω_ij_*. We refer to the modified, baseline value as *ω*_baseline;*ij*_. Rearranging Equation A11, we removed the estimated effect of temperature and precipitation deviations as:

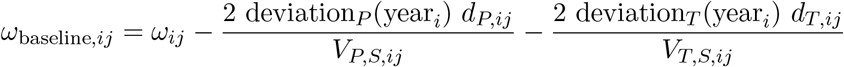

Estimated deviations in the year of each SSTFT combination *i* are calculated from parameters estimated from model A, denoted with the 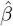:

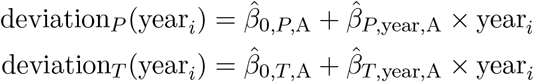

*d_P,ij_* and *d_T,ij_* are the historic differences between average source and site precipitation and temperature, respectively, for the *j*th observation at SSTFT combination *i*. *V_P,S,ij_* and *V_T,S,ij_* are the niche breadths for precipitation and temperature, calculated as:

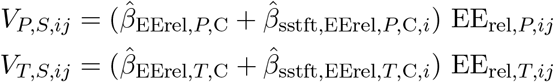

If temperature deviations make a large contribution to the increased variance in local adaptation, then the rate at which the variance increases through time should be reduced once the effect of deviations are statistically removed. We compared the posterior distributions of *β*_*ϵ*,C,year_ in models with and without the effect of deviations.

#### 3.5 Effects of temperature deviations on the prevalence of local adaptation

In the preceding analyses, we evaluated whether temperature and precipitation deviations alter the strength of local advantage—that is, the ratio of local to foreign fitness for any pair of populations compared. Shifting from this pairwise framework, local adaptation can also be conceptualized more generally as 1) the local population outperforming all foreign populations moved to its home site or 2) a population performing better in its home site than any other site that it is transplanted to.

If temperature deviations weaken local advantage and disrupt the fit of populations to their home environments, it follows that they may decrease the prevalence of local adaptation as defined above, even if they do not alter the average strength of local advantage. To test this, we regressed the binary outcome of whether a population outperforms all other populations in its home site on the absolute temperature deviation of that site during the experiment. We included covariates of the average composite distance that populations were moved, as well as the number of foreign populations that were moved to that site. We expected that the probability of the local population outperforming all foreign populations would decline under more anomalous thermal conditions. We also predicted that the probability of the local population outperforming all foreign populations would be higher if foreign populations were moved over greater distances, but might decrease as more foreign populations are included in the experiment, since that increases the probability that a foreign population might outperform the local population due to random chance and that one of the foreign populations might be better suited to the site environment.

We ran a similar regression in which the response variable was the binary outcome of whether a population performed best in its home site compared to other sites (after standardizing by the average performance of all populations in a site). We expected that the probability of a population performing best at home would decline under more anomalous thermal conditions. In this model we included covariates of the average distance that a population was moved and the number of sites a population was tested at. We expected greater distances to increase the probability a population would do best at home and more test sites to decrease the probability a population would do best at home.

These models were fit in Stan using the package **brms** [53], using a Bernoulli response distribution, a logit link function, and weakly informative priors. The number of populations or number of sites were included in the model as 1/(the number of populations or number of sites), because. We included a random intercept for each combination of study and taxon. Convergence was assessed as described below.

#### 3.6 Fitting models with Stan

We fit Bayesian mixed effects Models A-C using Stan [54], a probabilistic programming language, via the R package **rstan** version 2.18.2 [51]. In all models, we assigned diffuse normal priors (*μ* = 0, *σ* = 10) to fixed effect parameters and weakly informative half-cauchy priors (*μ* = 0, *σ* =1) to variance parameters (random effect and residual variances). Because some parameters are shared across models, all models were run simultaneously on two chains for 10^4^ warm-up and 10^4^ sampling, each, with a thinning interval of 10^1^. All key parameters (Tables A2 to A5) converged when the Gelman-Rubin 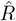 < 1.01 [55] and the effective sample size (ESS) was greater than 10^3^. 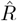 and ESS statistics were calculated using the ‘summarize_draws’ function in the R package **posterior** version 0.0.3 [56]. We tested *a priori* predictions using 95% highest posterior density intervals. For example, if we predicted a parameter should be greater than 0, then the 95% HPDI for that parameter should include only positive values.

**Table A4.** Model B2 parameters and predictions. Each parameter has a mathematical Symbol used in the text, a Stan character string used in the model code, a brief Description of the parameters, and associated *a priori* Predictions where applicable. Parameter estimates from fitted models are presented in Table S4 and Fig. 3CD.

| Symbol | Stan | Description | Prediction<br>(where<br>applicable) |
| --- | --- | --- | --- |
| Regression on $\log(w_{ij})$ | | | |
| $\beta_{0,B}$ | b0_B | Average $\log(w_{ij})$ when $m_T = 0$ and $m_P = 0$ | $\beta_{0,B} = 0$ |
| $\beta_{T1,B}$ | b1_t_B | Linear effect of $m_T$ on $\log(w_{ij})$ | $\beta_{T1,B} = 0$ |
| $\beta_{T2,B}$ | b2_t_B | Quadratic effect of $m_T$ on $\log(w_{ij})$ | $\hat{\beta}_{T2,B} < 0$ |
| $\beta_{P1,B}$ | b1_p_B | Linear effect of $m_P$ on $\log(w_{ij})$ | $\beta_{P1,B} = 0$ |
| $\beta_{P2,B}$ | b2_p_B | Quadratic effect of $m_P$ on $\log(w_{ij})$ | $\hat{\beta}_{P2,B} < 0$ |
| $\beta_{\text{local},B}$ | b_B_local | Effect of being local | $\hat{\beta}_{\text{local},B} > 0$ |
| $\beta_{\text{study\_taxon},B_i}$ | b_B_study_taxon[k] | Effect of study-taxon $k$ on $\log(w_{ij})$ | |
| $\sigma_{\text{study\_taxon},B}$ | sigma_B_study_taxon | Standard deviation of $\beta_{\text{study\_taxon},B_i}$ | |
| Regression on $\sigma_{\epsilon,B}$ | | | |
| $\beta_{\epsilon,B,n}$ | b_B_n | Effect of $\sqrt{n}$ on $\log(\sigma_{\epsilon,B})$ | |
| $\log(\sigma_{\epsilon,B,0})$ | log_sigma0_B_resid | Standard deviation (log-scale) of residual $\log(w_{ij})$ when $n = 0$ | |

**Table A5.** Model C parameters and predictions. Each parameter has a mathematical Symbol used in the text, a Stan character string used in the model code, a brief Description of the parameters, and associated *a priori* predictions where applicable.

| Symbol | Stan | Description | Prediction<br>(where<br>applicable) |
| --- | --- | --- | --- |
| Regression on $\omega$ | | | |
| $\beta_{0,C}$ | `b0_C` | Average strength of local adaptation ($\omega$) | $\hat{\beta}_{0,C} > 0$ |
| $\beta_{\text{dist},C}$ | `b_C_dist` | Effect of geographic distance on $\omega$ | $\hat{\beta}_{\text{dist},C} < 0$ |
| $\beta_{\text{germ},C}$ | `b_C_ft_germ` | Effect of germination fitness type on $\omega$ | |
| $\beta_{\text{germ\_surv},C}$ | `b_C_ft_germ_surv` | Effect of germination-survival fitness type on $\omega$ | |
| $\beta_{\text{surv},C}$ | `b_C_ft_surv` | Effect of survival fitness type on $\omega$ | |
| $\beta_{\text{repro},C}$ | `b_C_ft_repro` | Effect of reproduction fitness type on $\omega$ | |
| $\beta_{\text{year},C}$ | `b_C_year` | Effect of study year on $\omega$ | $\hat{\beta}_{\text{year},C} = 0$ |
| Precipitation-specific parameters |  |  |  |
| $\beta_{\text{EErel},P,C}$ | `b_p_C_eerel` | Effect of precipitation $\text{EE}_{\text{rel}}$ on local adaptation, inverse of niche breadth | $\hat{V}_{S,P}^{-1} = \hat{\beta}_{\text{EErel},P,C} > 0$ |
| $\beta_{\text{sstft},\text{EErel},P,C,i}$ | `b_p_C_eerel_sstft[i]` | Effect of $\text{SSTFT}_i$ on $\beta_{\text{EErel},P,C}$ | |
| $\sigma_{\text{sstft},\text{EErel},P,C}$ | `sigma_p_C_eerel_sstft` | Standard deviation of $\beta_{\text{sstft},\text{EErel},P,C,i}$ | |
| Temperature-specific parameters |  |  |  |
| $\beta_{\text{EErel},T,C}$ | `b_t_C_eerel` | Effect of precipitation $\text{EE}_{\text{rel}}$ on local adaptation, inverse of niche breadth | $\hat{V}_{S,T}^{-1} = \hat{\beta}_{\text{EErel},T,C} > 0$ |
| $\beta_{\text{sstft},\text{EErel},T,C,i}$ | `b_t_C_eerel_sstft[i]` | Effect of $\text{SSTFT}_i$ on $\beta_{\text{EErel},T,C}$ | |
| $\sigma_{\text{sstft},\text{EErel},T,C}$ | `sigma_t_C_eerel_sstft` | Standard deviation of $\beta_{\text{sstft},\text{EErel},T,C,i}$ | |
| Regression on $\sigma_{\epsilon,C}$ | | | |
| $\beta_{\epsilon,C,\text{year}}$ | `b_C_resid_year` | Effect of study year on $\log(\sigma_{\epsilon,C})$ | $\hat{\beta}_{\epsilon,C,\text{year}} > 0$ |
| $\log(\sigma_{\epsilon,C,0})$ | `log_sigma0_C_resid` | Scale parameter (log-scale) of residual $\omega$ in study year 0 (= 1975) | |
| $\nu_C$ | `nu_C_resid` | Degrees of freedom parameter of residual $\omega$ | |

### 4 Supplemental results

#### 4.1 Model performance and adequacy

Trace plots and posterior densities for all key parameters also indicated good mixing and stationarity (Fig. S1 and. ‘mcmc_areas’, ‘mcmc_trace’, and ‘ppc_dens_overlay’ functions from the R package **bayesplot** version 1.7.2 [57] were used for area, trace, and posterior predictive check plots, respectively.

Area and trace plots are included in an additional pdf. The parameter is listed at the top. In area plots, the plotting region shows the posterior density of the parameter, the point estimate (median of the posterior, dark blue line), and 80% probability mass interval (shaded blue area). In trace plots, the plotting region show lines plots of the parameter value during sampling for both chains.

**Fig. S1.**
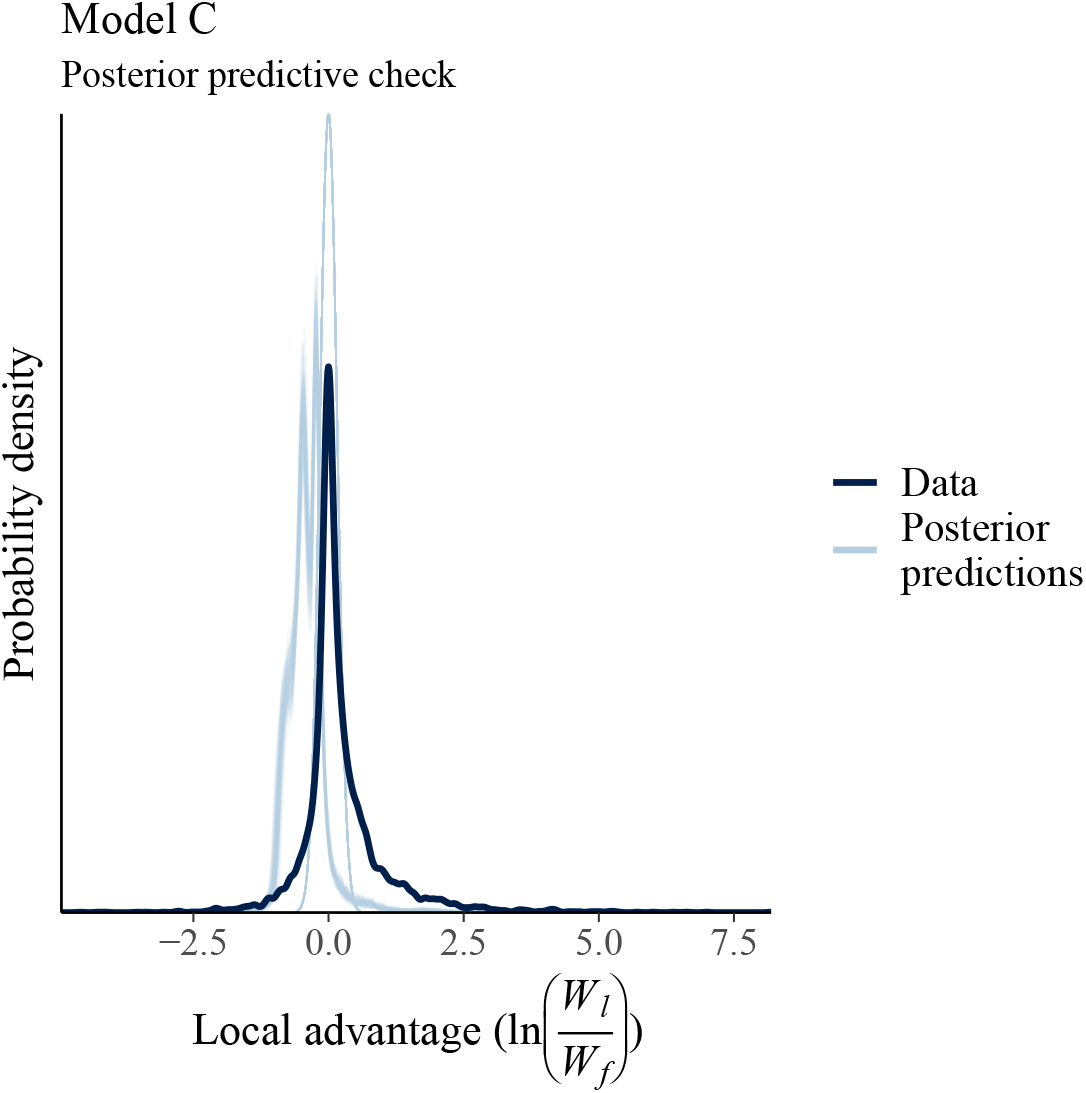
Posterior predictive check of the response variable (local advantage) from Model C. Agreement between the data (dark blue line) and simulated responses from 1000 draws of model posterior (light blue lines) suggested the model adequately fits the data.

#### 4.2 Comparing parameters from Models B and C

We estimate local adaptation to factors other than average temperature and precipitation in both Model B and Model C. In Model B, this is the effect of being local (vs. foreign), and in Model C it is the intercept, i.e., the magnitude of local adaptation when the experimental climate conditions favor both local and foreign populations equally. Estimates of the costs of being foreign from Model C are less severe than those of Model B (Fig. S2A, Table S4, Table S5). This may be because Model C accounts for composite (geographic and elevational) distance between the foreign population and the transplant site, and this parameter likely accounts for the magnitude of local adaptation to biotic and abiotic factors other than average temperature and precipitation that are collinear with distances between sites.

We estimate the sensitivity of fitness to variation in climate (*V_S_*) in both Model B and Model C. We find that our estimate of VS is larger in model B (Fig. S2, Table 3.2) than in model C (Table 3.3). This may occur because because our estimate from Model B includes both the sensitivity to climate as well as variability in the optima of the foreign populations that we have used to relativize fitness. The optima of foreign populations are included in our estimate of EE_rel_, and as a result, the estimate of *V_S_* from Model C is likely to more accurately reflect the sensitivity of populations to variation in climate.

**Fig. S2.**
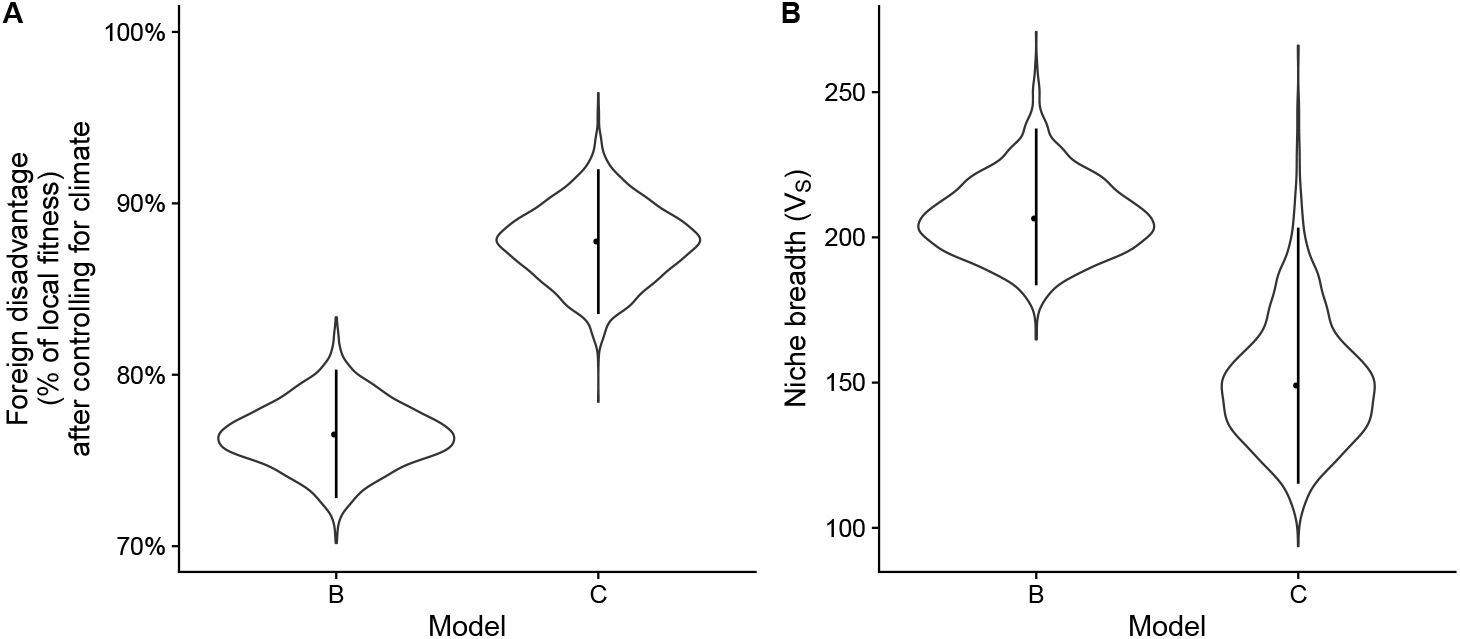
(**A**) The magnitude of local adaptation to non-climatic factors estimated using two approaches, with the parameter *β*_local,B_ in Model B and *β*_0;C_ in Model C. (**B**) We also estimate temperature niche breadth using these two models, with the Gaussian variance parameter in Model B and the inverse of the estimate of temperature EE_rel_ in Model C.

#### 4.3 Increasing variance may reduce the overall frequency of local advantage

Even in the absence of a change in the average strength of local advantage over time, an increase in the variance in experimental outcomes should lead to an increase in the proportion of pairwise contrasts in which the foreign population outperforms the local population (i.e., the relative width of the blue vs. green shaded areas in Fig. 4B, which is plotted as the orange line and interval in Fig. S3). This decrease in the frequency of local advantage over time does not emerge from our data (grey line and interval in Fig. S3, *β*_year_ = −0.01, 95% HDI: −0.03 – 0.01). Low data density in early years may reduce our ability to detect temporal trends.

**Fig. S3.**
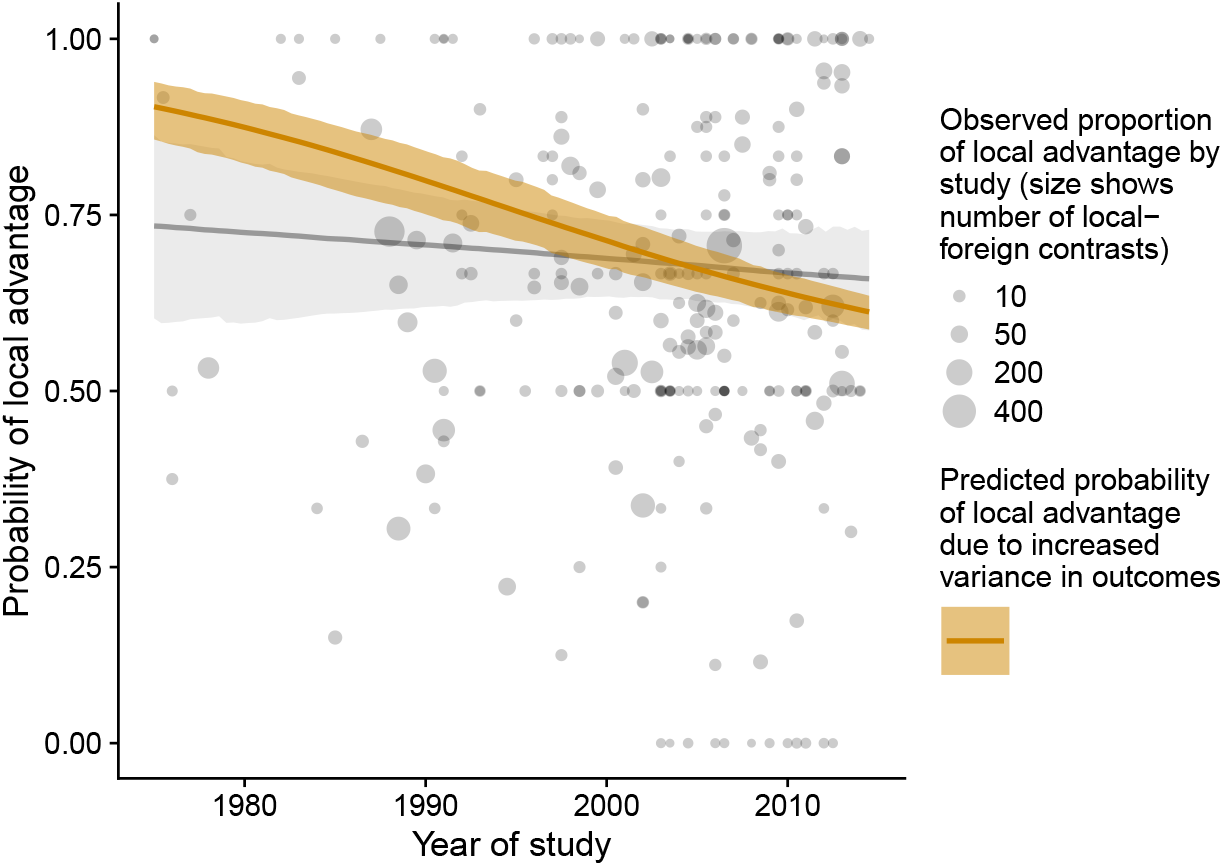
Increased variance in experimental outcomes (Table S5, Fig. 4B) is expected to decrease the probability of detecting local advantage in transplant experiments (orange line). This pattern does not emerge from studies to-date (grey points, the grey line and shaded interval represent estimates from a binomial model).

#### 4.4 Temperature and precipitation deviations cannot explain most of the increased variance in local adaptation

We compared the change in variance through time in Model C to a modified model in which the effect of temperature and precipitation deviations were statistically removed (Section 3.4). This did not substantially reduce the rate at which variance increased (Figure S4), indicating that changes in other environmental parameters and/or experimental practice must account for most of this pattern.

**Fig. S4.**
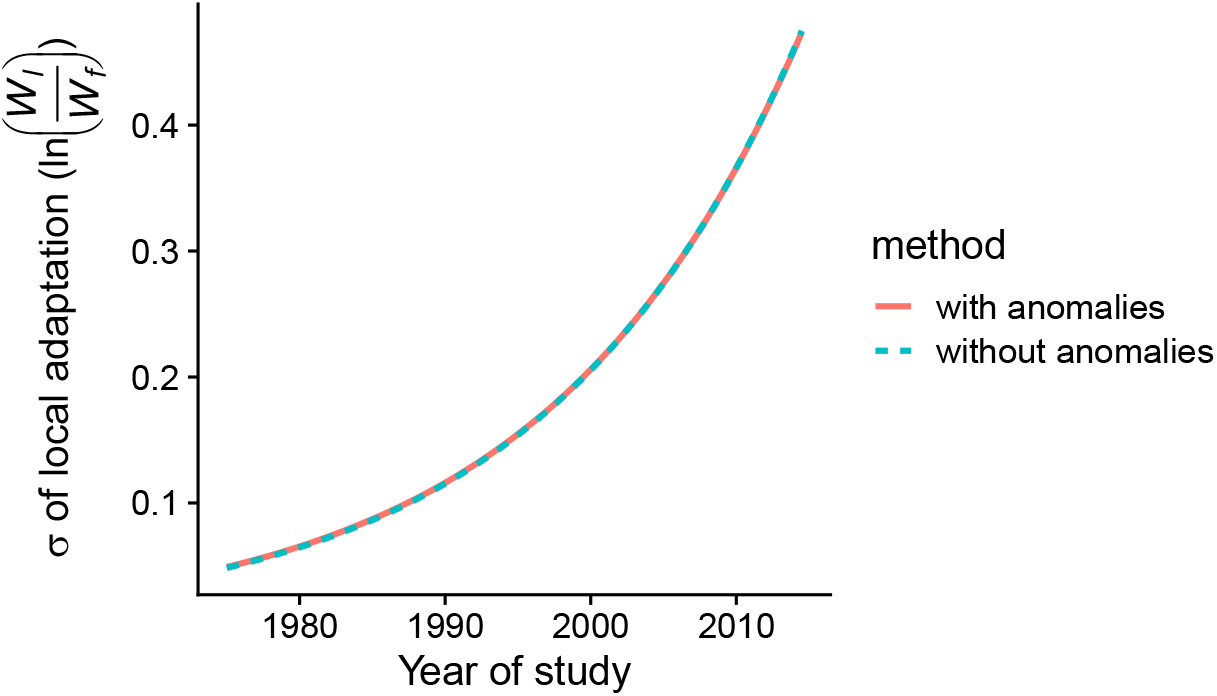
Temperature and precipitation deviations cannot explain most of the increased variance (*σ*^2^) in local adaptation (*y*-axis). The dashed line shows the median estimated residual standard deviation in Model C. The solid line is the same parameter with the effect of temperature and precipitation deviations statistically removed (Section 3.4). The change in variance through time is similar in both models, indicating that these climate deviations do not explain most of this pattern.

#### 4.5 Temperature anomalies decrease the prevalence of local adaptation

Temperature deviations decrease the probability that 1) the local population will outperform all foreign populations in a site, and 2) that a population will perform best in its home site (Fig 4D, Table S7). These probabilities also depend on the number of sites or populations being compared (Fig. S5). When more populations are compared in a site, this reduces the probability that the local population will be the best performer. Similarly, the more sites a population is tested at, the lower the probability that it will do best in its home site. The average composite distance that populations are moved also affects the prevalence of local adaptation: when foreign populations are moved across greater distances, or when a population is tested at sites that are further from its home, the probability of the local population being best or the home site being best is higher (Table S7).

**Fig. S5.**
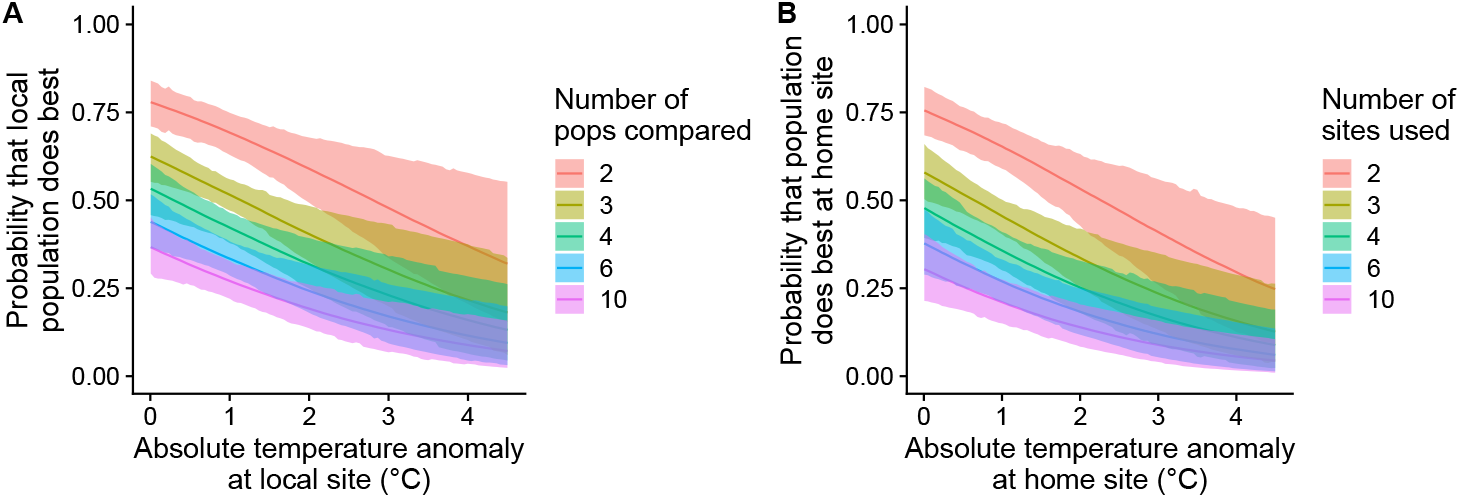
Temperature anomalies decrease (**A**) the probability that the local population outperforms foreign populations in a site or (**B**) the probability that a population performs best in its home site relative to other (“away”) sites. The more populations (**A**) or sites (**B**) that are compared, the lower the probability of the local population or home site being best.

### 5 Supplemental tables

#### 5.1 Table S1: Included studies

**Table S1.**
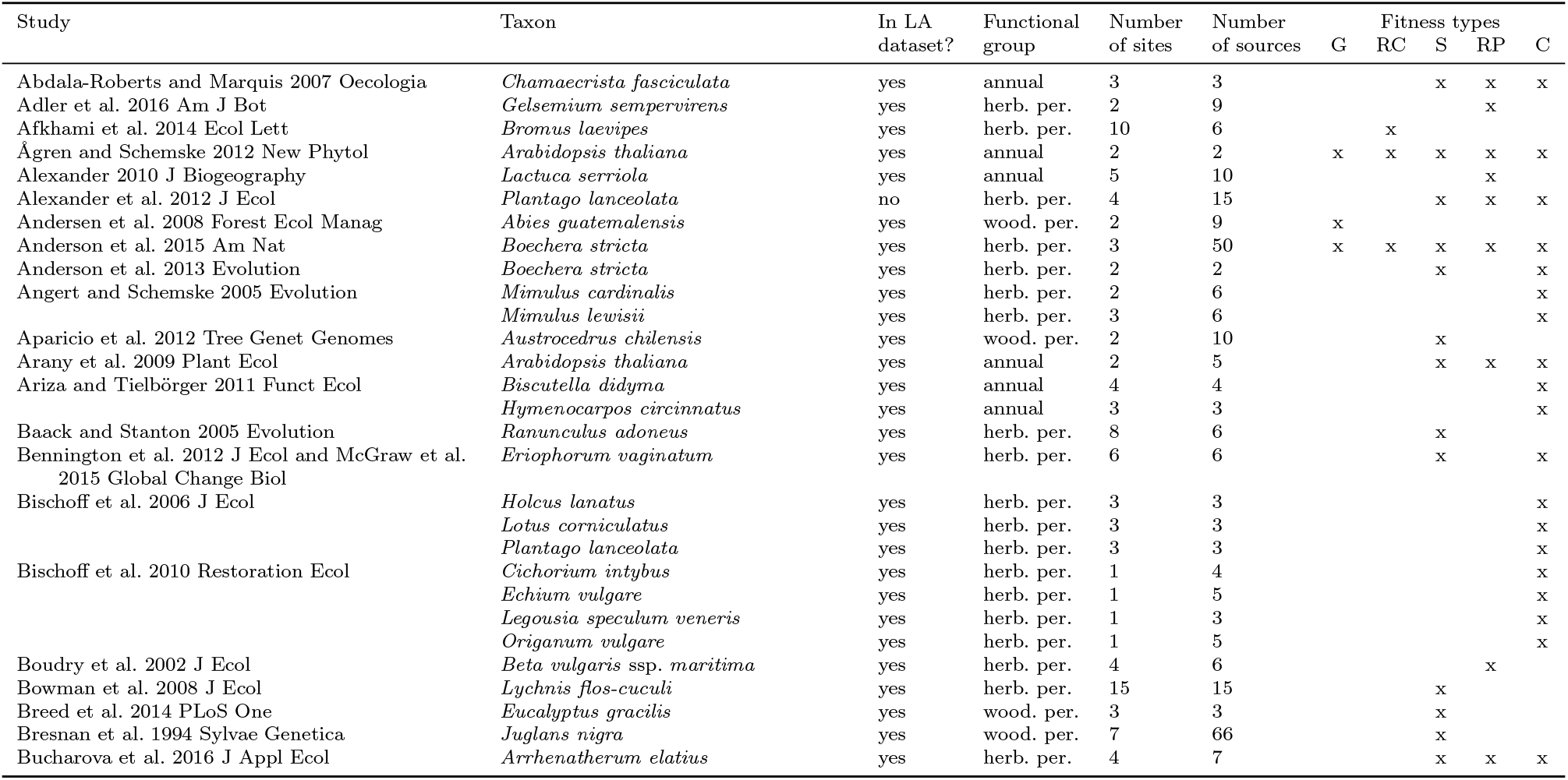
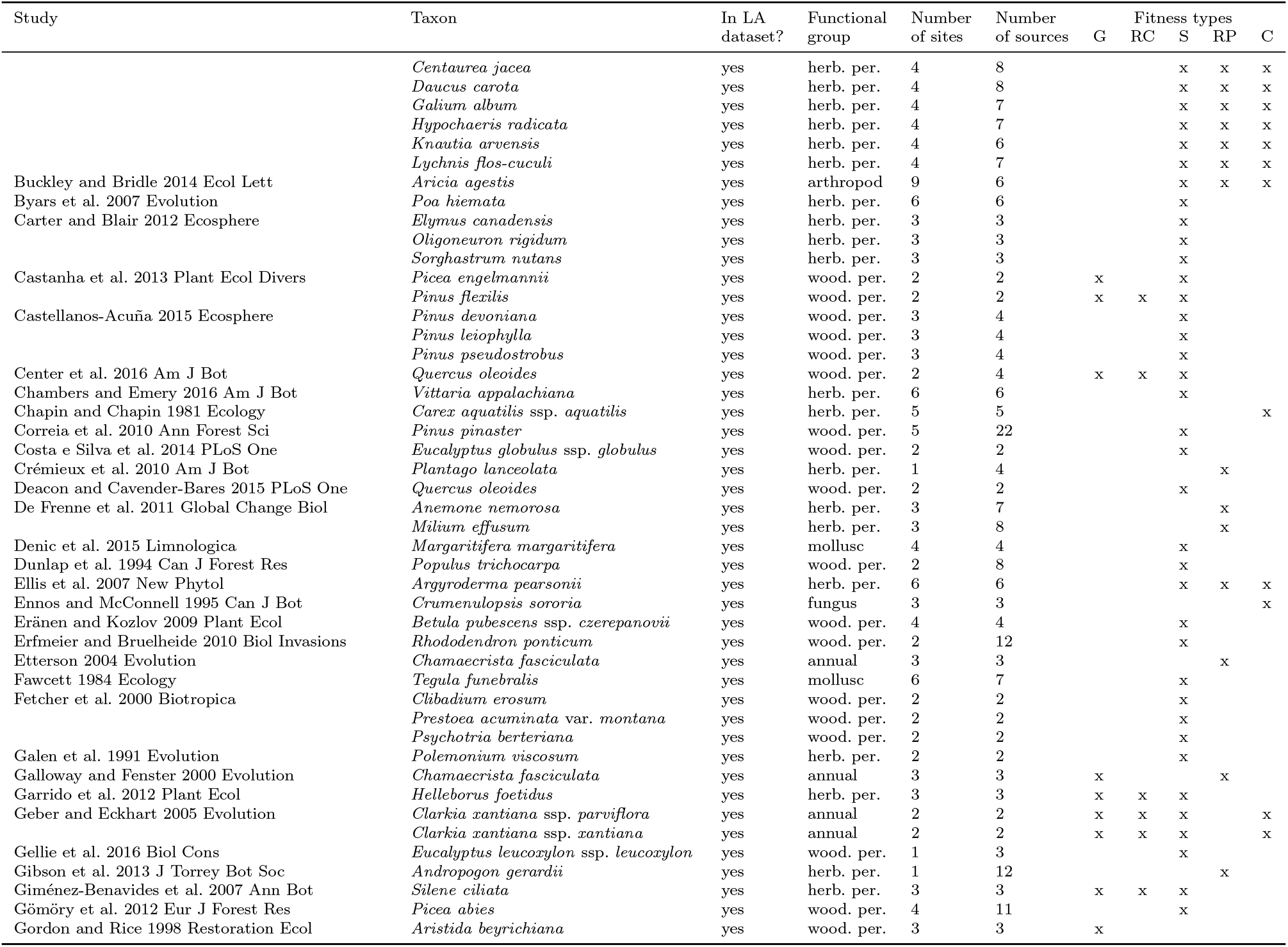
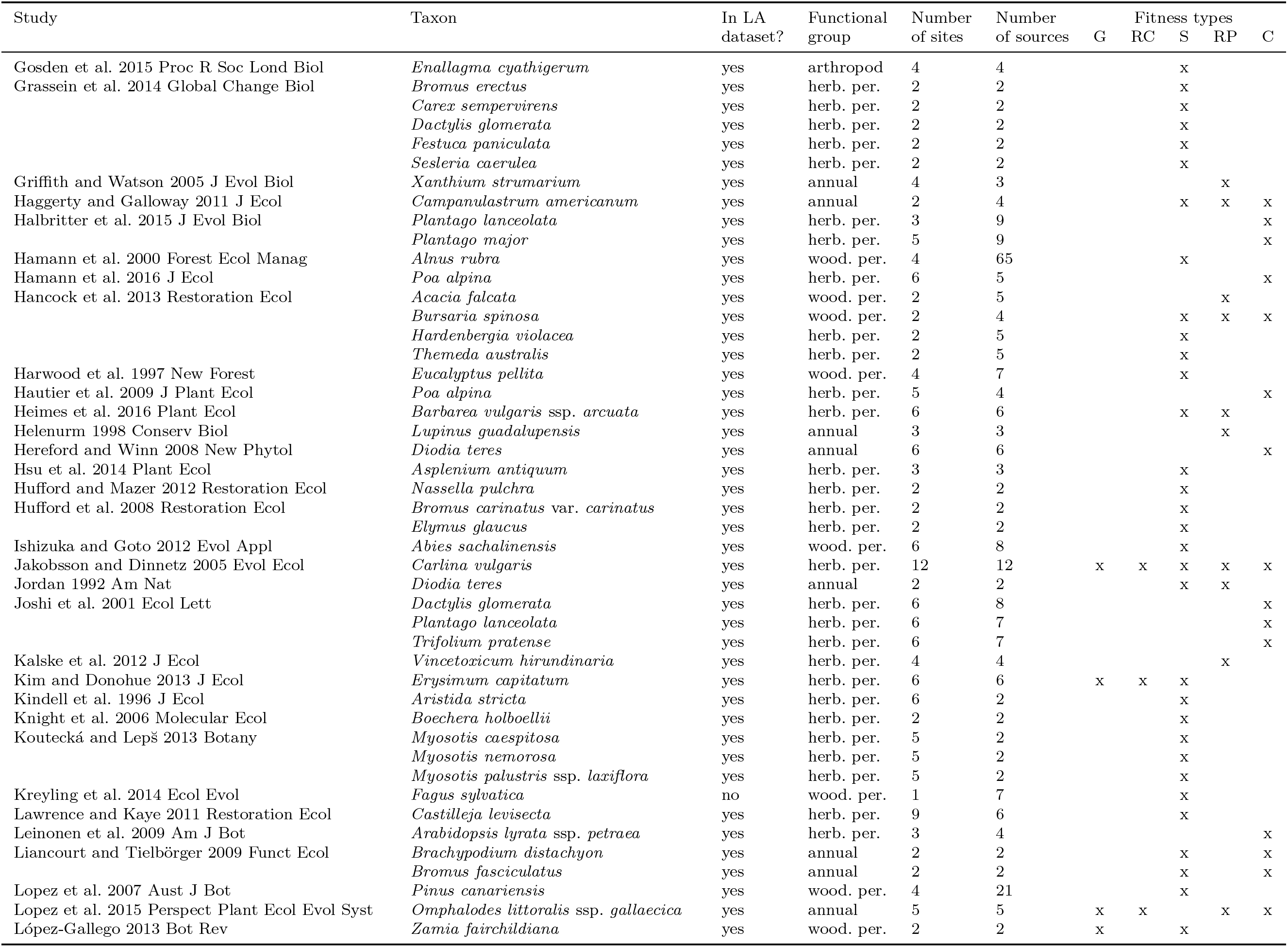
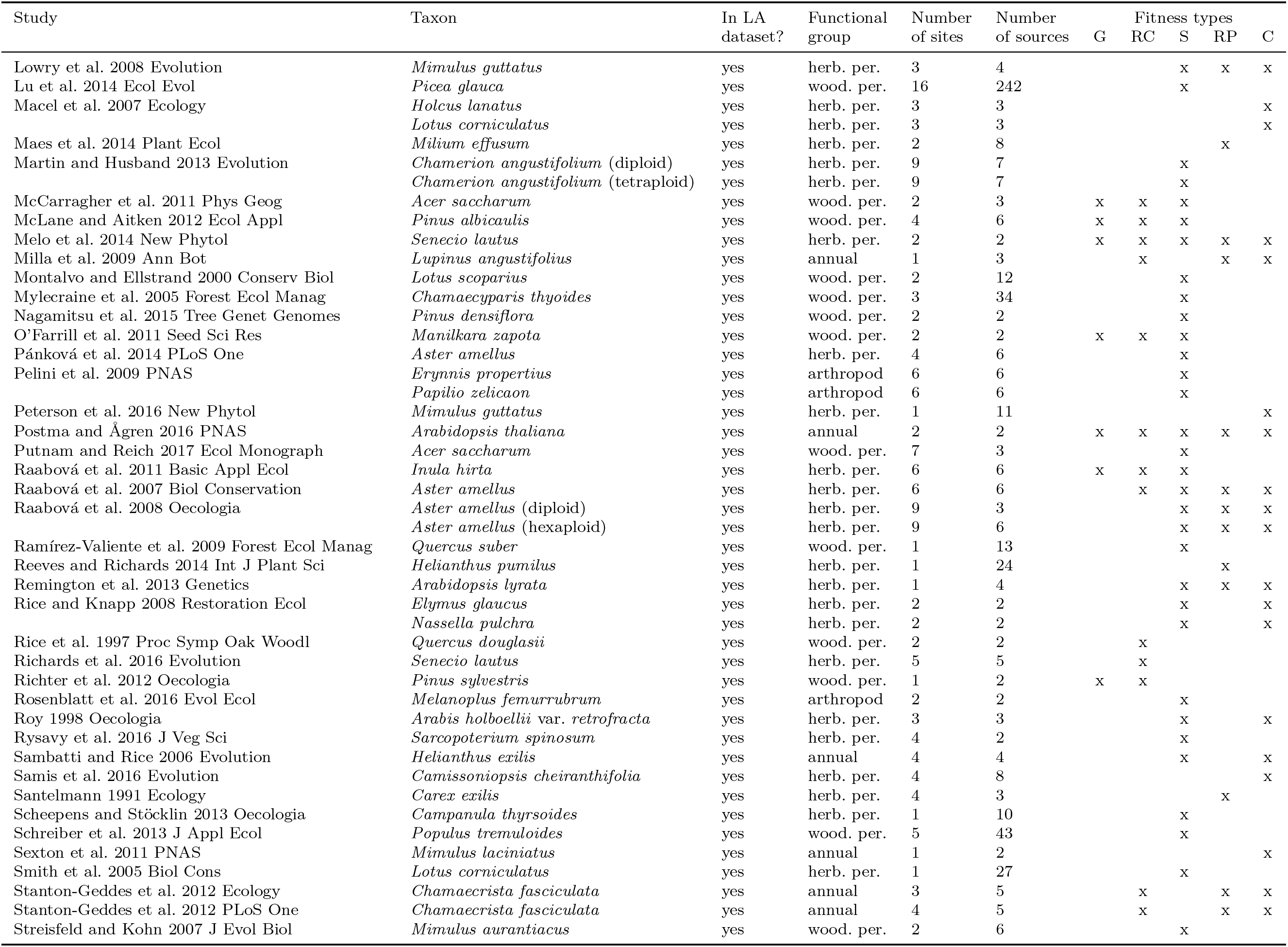
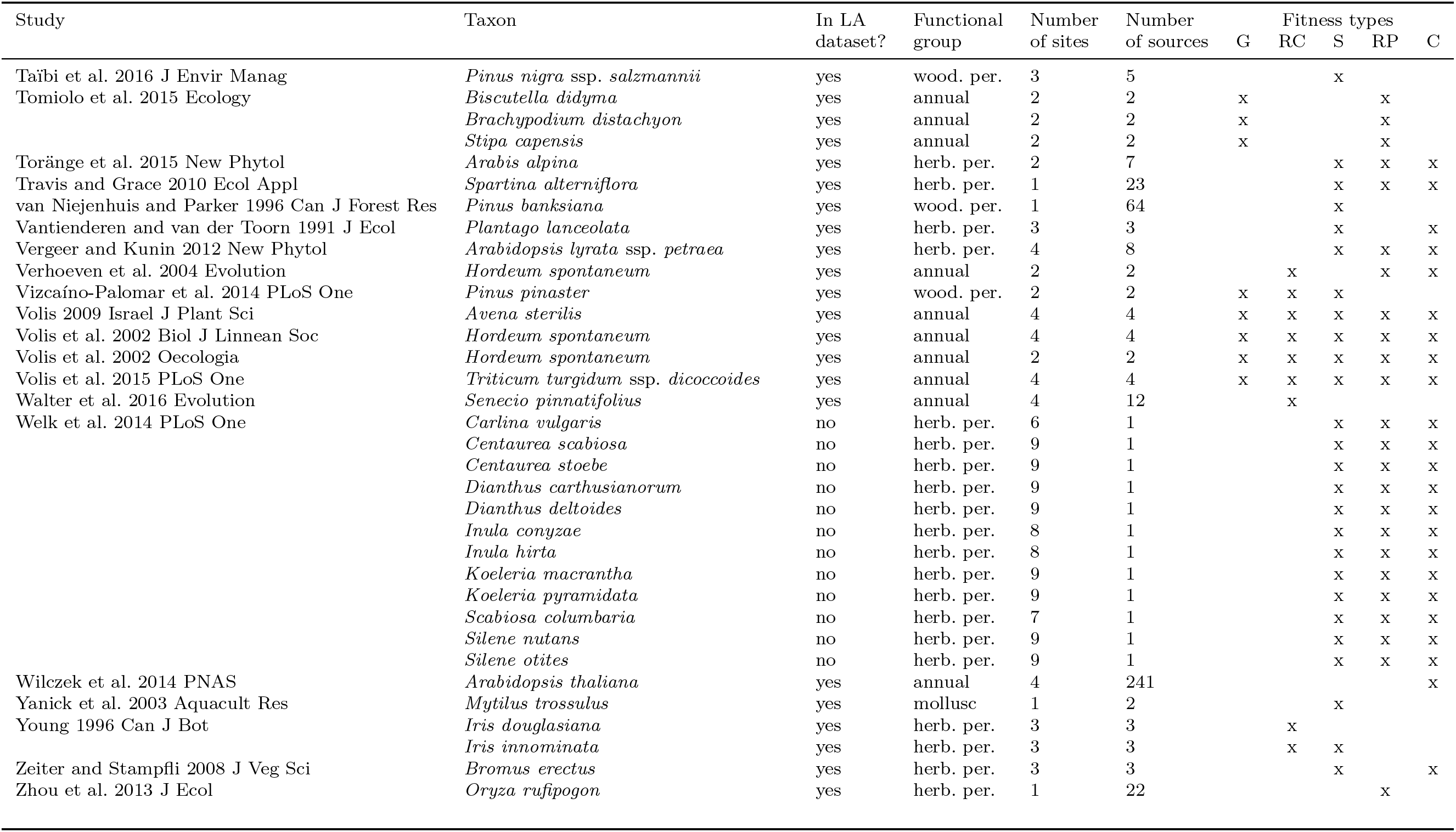
Studies included in our database. All studies were used in analyses of relative fitness, but only a subset of studies that included local populations in their design were included in our analyses of local adaptation. Functional groups include annual plant (annual), herbaceous perennial (herb. per.), woody perennial (wood. per.), fungus, arthropod, and mollusk. Fitness types are germination (G), recruitment (RC; germination and subsequent survival), survival (S), reproduction (RP), composite (C; lifetime fitness estimate incorporating at least survival and reproduction). Note that in some cases some fitness types were only measured for a subset of sites or sources, but the total numbers of sites and sources for each taxon in each study are tallied here.

#### 5.2 Table S2: Model A results (climate deviations over time)

**Table S2.** Bayesian mixed effect model estimates of the change in the frequency and magnitude of (A) temperature, (B) precipitation and (C) absolute precipitation deviations over time (Model A). Estimates of slope with credible intervals that do not include 0 are indicated with bold text. In these models, year is scaled so that the intercept represents the average deviation size in 1975. SE = standard error; Lower/Upper CI = lower/upper 95% highest posterior density interval from the model posterior; 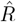 = Gelman-Rubin convergence statistic (1 = convergence); ESS = effective sample size from the posterior. Temperature deviations are expressed as deviations from normal in degrees Celsius. Precipitation deviations are expressed as the log_10_ ratio of experiment precipitation to normal precipitation. Temperature and precipitation results are plotted in Fig. 2CD.

| Term | Estimate | SE | Lower CI | Upper CI | $\hat{R}$ | ESS |
| --- | --- | --- | --- | --- | --- | --- |
| *A. Temperature* |  |  |  |  |  |  |
| Intercept (year 1975 = 0, $\beta_{0,T,A}$) | 0.071 | 0.092 | -0.11 | 0.25 | 1.00 | 1995 |
| **Slope ($\beta_{T,\text{year},A}$)** | **0.025** | **0.0032** | **0.018** | **0.031** | **1.00** | **1966** |
| Effect of year on residual variance ($\beta_{\epsilon,T,A,\text{year}}$) | 0.0044 | 0.0040 | -0.0038 | 0.012 | 1.00 | 1964 |
| Site SD ($\sigma_{T,\text{site},A}$) | 0.43 | 0.033 | 0.36 | 0.49 | 1.00 | 1661 |
| Residual SD in year 1975 (log-scale, $\log(\sigma_{\epsilon T,A,0})$) | -0.72 | 0.13 | -0.97 | -0.47 | 1.00 | 1952 |
| *B. Precipitation* |  |  |  |  |  |  |
| Intercept (year 1975 = 0, $\beta_{0,P,A}$) | -0.046 | 0.021 | -0.085 | -0.0057 | 1.00 | 1790 |
| Slope ($\beta_{P,\text{year},A}$) | 0.0011 | 0.00068 | -0.00025 | 0.0024 | 1.00 | 1877 |
| **Effect of year on residual variance ($\beta_{\epsilon,P,A,\text{year}}$)** | **-0.263** | **0.0038** | **-0.034** | **-0.019** | **1.00** | **1926** |
| Site SD ($\sigma_{P,\text{site},A}$) | 0.064 | 0.0057 | 0.052 | 0.075 | 1.00 | 1484 |
| Residual SD in year 1975 (log-scale, $\log(\sigma_{\epsilon P,A,0})$) | -1.54 | 0.11 | -1.74 | -1.32 | 1.00 | 2017 |
| *C. Absolute precipitation* |  |  |  |  |  |  |
| Intercept (year 1975 = 0, $\beta_{0,AP,A}$) | 0.094 | 0.017 | 0.061 | 0.13 | 1.00 | 1827 |
| Slope ($\beta_{AP,\text{year},A}$) | -0.00059 | 0.00055 | -0.0017 | 0.00050 | 1.00 | 1843 |
| **Effect of year on residual variance ($\beta_{\epsilon,AP,A,\text{year}}$)** | **-0.374** | **0.0044** | **-0.046** | **-0.029** | **0.99** | **1897** |
| Site SD ($\sigma_{AP,\text{site},A}$) | 0.059 | 0.0037 | 0.052 | 0.066 | 1.00 | 1716 |
| Residual SD in year 1975 (log-scale, $\log(\sigma_{\epsilon AP,A,0})$) | -1.62 | 0.122 | -1.85 | -1.37 | 0.99 | 1852 |

#### 5.3 Table S3: Model B1 results (relative fitness vs. site conditions)

**Table S3.**
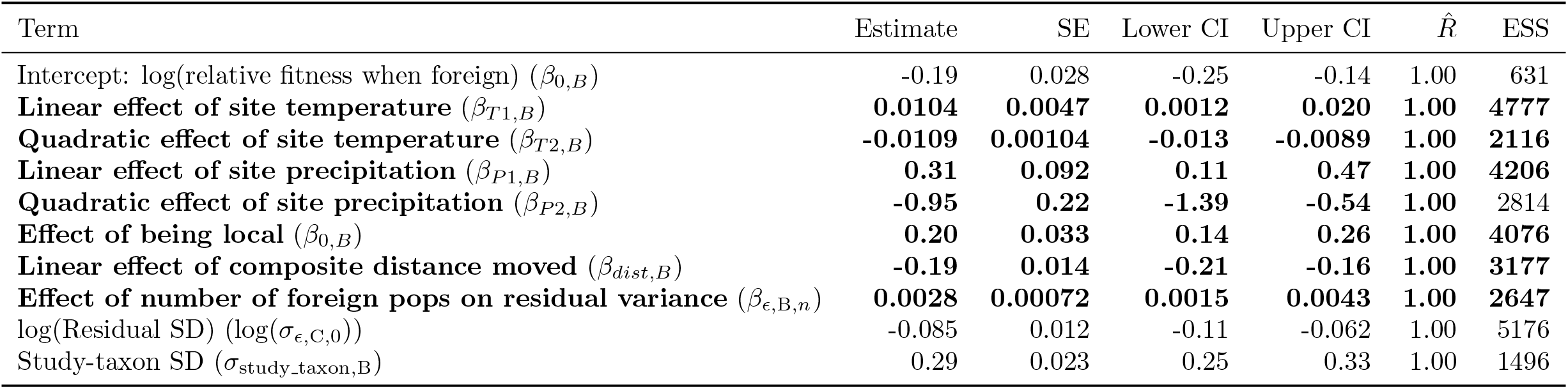
Bayesian mixed effect model estimates of quadratic fits of the effects of temperature and precipitation mismatch on relative fitness (Model B). Parameters of interest with credible intervals that do not include 0 are indicated with bold text. SE = standard error; Lower/Upper CI = lower/upper 95% highest posterior density interval from the model posterior; 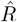 = Gelman-Rubin convergence statistic (1 = convergence); ESS = effective sample size from the posterior. These effects are plotted in Fig. 3AB.

#### 5.4 Table S4: Model B2 results (relative fitness vs. climate mismatch)

**Table S4.** Bayesian mixed effect model estimates of quadratic fits of the effects of temperature and precipitation mismatch on relative fitness (Model B). Parameters of interest with credible intervals that do not include 0 are indicated with bold text. SE = standard error; Lower/Upper CI = lower/upper 95% highest posterior density interval from the model posterior; 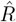 = Gelman-Rubin convergence statistic (1 = convergence); ESS = effective sample size from the posterior. These effects are plotted in Fig. 3CD.

| Term | Estimate | SE | Lower CI | Upper CI | $\hat{R}$ | ESS |
| --- | --- | --- | --- | --- | --- | --- |
| Intercept: log(foreign relative fitness) ($\beta_{0,B}$) | -0.11 | 0.023 | -0.15 | -0.064 | 1.00 | 1970 |
| **Linear effect of temperature mismatch ($\beta_{T1,B}$)** | **0.0160** | **0.0029** | **0.010** | **0.021** | **1.00** | **2071** |
| **Quadratic effect of temperature mismatch ($\beta_{T2,B}$)** | **-0.0048** | **0.00031** | **-0.0054** | **-0.0042** | **1.00** | **1917** |
| Linear effect of precipitation mismatch ($\beta_{P1,B}$) | 0.11 | 0.060 | -0.0090 | 0.23 | 1.00 | 1928 |
| Quadratic effect of precipitation mismatch ($\beta_{P2,B}$) | -0.043 | 0.11 | -0.25 | 0.17 | 1.00 | 1926 |
| **Effect of being local ($\beta_{0,B}$)** | **0.27** | **0.025** | **0.22** | **0.32** | **1.00** | **1888** |
| **Effect of number of foreign pops on residual variance ($\beta_{\epsilon,B,n}$)** | **0.028** | **0.0016** | **0.025** | **0.031** | **1.00** | **1953** |
| log(Residual SD) ($\log(\sigma_{\epsilon,C,0})$) | -0.38 | 0.012 | -0.41 | -0.36 | 1.00 | 2070 |
| Study-taxon SD ($\sigma_{\text{study\_taxon},B}$) | 0.26 | 0.018 | 0.22 | 0.29 | 1.00 | 1797 |
| **Optimal temperature mismatch** | **1.6** | **0.28** | **1.0** | **2.2** | **1.00** | **2132** |
| **Gaussian variance in temperature response** | **210** | **14** | **180** | **230** | **1.00** | **1917** |

#### 5.5 Table S5: Model C results (local adaptation vs. climate deviations)

**Table S5.** Bayesian mixed effect model estimates of the effect of temperature and precipitation deviations on local adaptation (the log ratio of the performance of local populations to the performance of foreign populations). Models were run with (A) temperature and (B) precipitation. Temperature and precipitation deviations are represented by EE_rel_, which relates experimental conditions to the historic conditions of the local and foreign populations based on Gaussian fitness responses to deviations in climate. Also included are the composite geographic distance between the foreign source and the transplant site and the effects of different fitness proxies on local adaptation. Parameters with credible intervals that do not include 0 are indicated with bold text. SE = standard error; Lower/Upper CI = lower/upper 95% highest posterior density interval from the model posterior; 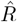 = Gelman-Rubin convergence statistic (1 = convergence); ESS = effective sample size from the posterior. The effects of temperature EE_rel_ are shown in Fig. 4A, the change in variance over time is shown in Fig. 4B, and the expected effects of this change in variance on local advantage is shown in Fig. S3

| Term | Estimate | SE | Lower CI | Upper CI | $\hat{R}$ | ESS |
| --- | --- | --- | --- | --- | --- | --- |
| *Regression on $\omega$* |  |  |  |  |  |  |
| **Intercept (when deviation favors neither local or foreign, $\beta_{0,C}$)** | **0.13** | **0.025** | **0.085** | **0.18** | **1.00** | **1947** |
| Effect of year ($\beta_{year,C}$) | 0.00089 | 0.00059 | -0.00042 | 0.0019 | 1.00 | 1941 |
| **Effect of distance between local and foreign pops ($\beta_{dist,C}$)** | **0.020** | **0.0073** | **0.0053** | **0.034** | **1.00** | **1960** |
| **Effect of fitness type = germination ($\beta_{germ,C}$)** | **-0.12** | **0.032** | **-0.18** | **-0.058** | **1.00** | **1973** |
| **Effect of fitness type = germ + surv ($\beta_{germ\_surv,C}$)** | **-0.071** | **0.032** | **-0.13** | **-0.0057** | **1.00** | **1996** |
| **Effect of fitness type = reproduction ($\beta_{repro,C}$)** | **-0.056** | **0.029** | **-0.11** | **-0.00023** | **1.00** | **1860** |
| **Effect of fitness type = survival ($\beta_{surv,C}$)** | **-0.14** | **0.020** | **-0.18** | **-0.10** | **1.00** | **1887** |
| **Effect of temperature $EE_{rel}$ ($\beta_{EErel,T,C}$)** | **0.0067** | **0.00095** | **0.0049** | **0.0086** | **1.00** | **1795** |
| SD of study-site-taxon-fitness type effects of temp. $EE_{rel}(\sigma_{sstft,EErel,T,C})$ | 0.013 | 0.00098 | 0.011 | 0.015 | 1.00 | 417 |
| Effect of precipitation $EE_{rel}$ ($\beta_{EErel,P,C}$) | -0.016 | 0.24 | -0.51 | 0.46 | 1.00 | 1296 |
| SD of study-site-taxon-fitness type effects of precip. $EE_{rel}(\sigma_{sstft,EErel,P,C})$ | 2.7 | 0.37 | 2.0 | 3.4 | 1.00 | 126 |
| *Regression on $\sigma$* |  |  |  |  |  |  |
| Effect of year on residual variance ($\beta_{\epsilon,C,year}$) | 0.057 | 0.0021 | 0.053 | 0.061 | 1.00 | 1888 |
| Scale parameter of residual $\omega$ in study year 0 (= 1975) ($\log(\sigma_{\epsilon,C,0})$) | -3.0 | 0.061 | -3.1 | -2.9 | 1.00 | 1865 |

#### 5.6 Table S6: Reinforcing vs. counteracting deviations

**Table S6.** Bayesian mixed effect model estimates of the interactive effects of site conditions (normal vs. warm) and the climatic origin of the foreign population (warmer or cooler than the local population) on local adaptation. Parameter estimates with credible intervals that do not include 0 are indicated with bold text. SD = standard deviation; Lower/Upper CI = lower/upper 95% highest posterior density interval from the model posterior; 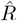 = Gelman-Rubin convergence statistic (1 = convergence); ESS = effective sample size from the posterior. Effects are plotted in Fig. 4C.

| Term | Estimate | SD | Lower CI | Upper CI | $\hat{R}$ | ESS |
| --- | --- | --- | --- | --- | --- | --- |
| **Intercept ($\beta_{0,C}$)** | **0.28** | **0.076** | **0.15** | **0.40** | **248** | **1.00** |
| **Effect of site condition (normal vs. warm)** | **-0.35** | **0.14** | **-0.60** | **-0.12** | **391** | **1.00** |
| Effect of foreign origin (warmer vs. cooler) | -0.0061 | 0.013 | -0.028 | 0.014 | 1888 | 1.00 |
| **Interaction between site condition and foreign origin** | **0.40** | **0.055** | **0.31** | **0.49** | **1616** | **1.00** |
| **Effect of composite distance of foreign source** | **0.076** | **0.0090** | **0.061** | **0.090** | **1707** | **1.00** |
| Effect of fitness type = germination | 0.089 | 0.12 | -0.12 | 0.27 | 536 | 1.00 |
| Effect of fitness type = germination + survival | 0.044 | 0.10 | -0.11 | 0.22 | 602 | 1.00 |
| Effect of fitness type = reproduction | 0.13 | 0.074 | 0.013 | 0.26 | 473 | 1.00 |
| Effect of fitness type = survival | -0.014 | 0.066 | -1.2 | 1.0 | 437 | 1.00 |
| Study-taxon SD | 0.47 | 0.049 | 0.40 | 0.56 | 413 | 1.00 |
| Residual SD | 0.20 | 0.0082 | 0.19 | 0.21 | 1271 | 1.00 |

#### 5.7 Table S7: Prevalence of local adaptation vs. temperature anomalies

**Table S7.** Bayesian mixed effect model estimates of the effect of temperature anomalies on (A) the probability that the local population outperforms foreign populations in a site or (B) the probability that a population performs best in its home site relative to other (“away”) sites. Also included in the models are the number of populations compared or the number of sites tested, and the average composite distance that populations were moved. Parameters of interest with credible intervals that do not include 0 are indicated with bold text. SD = standard deviation; Lower/Upper CI = lower/upper 95% highest posterior density interval from the model posterior; 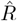 = Gelman-Rubin convergence statistic (1 = convergence); ESS = effective sample size from the posterior. Parameter estimates are on the logit link scale. Effects on the response scale are plotted in Fig. 4D and Fig. S5.

| Term | Estimate | SD | Lower CI | Upper CI | ESS | $\hat{R}$ |
| --- | --- | --- | --- | --- | --- | --- |
| *A. Probability that the local population outperforms foreign populations in a site* |  |  |  |  |  |  |
| Intercept | -0.99 | 0.22 | -1.42 | -0.57 | 2854 | 1.00 |
| **Absolute temperature anomaly in site** | **-0.45** | **0.13** | **-0.70** | **-0.19** | **2293** | **1.00** |
| **1/(Number of populations compared)** | **4.51** | **0.56** | **3.43** | **5.65** | **2319** | **1.00** |
| **Mean composite distance of foreign populations** | **0.27** | **0.09** | **0.09** | **0.47** | **2292** | **1.00** |
| Study-taxon SD | 0.61 | 0.14 | 0.34 | 0.90 | 572 | 1.00 |
| *B. Probability that a population performs best in its home site relative to other sites* |  |  |  |  |  |  |
| Intercept | -1.30 | 0.28 | -1.88 | -0.79 | 2290 | 1.00 |
| **Absolute temperature anomaly in home site** | **-0.50** | **0.15** | **-0.78** | **-0.21** | **2320** | **1.00** |
| **1/(Number of sites tested)** | **4.86** | **0.66** | **3.59** | **6.23** | **2451** | **1.00** |
| **Mean composite distance of away sites** | **0.40** | **0.12** | **0.19** | **0.64** | **2126** | **1.00** |
| Study-taxon SD | 0.68 | 0.15 | 0.41 | 0.98 | 607 | 1.00 |

## References

[1] S. Diaz, et al. (2020).

[2] J. R. Auld, A. A. Agrawal, R. A. Relyea, Proceedings of the Royal Society B: Biological Sciences 277, 503 (2010).

[3] S. Souther, J. B. McGraw, Conservation Biology 25, 922 (2011).

[4] P. J. Van Mantgem, N. L. Stephenson, Ecology Letters 10, 909 (2007).

[5] B. G. Freeman, M. N. Scholer, V. Ruiz-Gutierrez, J. W. Fitzpatrick, Proceedings of the National Academy of Sciences 115, 11982 (2018).

[6] G. E. Rehfeldt, C. C. Ying, D. L. Spittlehouse, D. A. Hamilton Jr, Ecological Monographs 69, 375 (1999).

[7] M. L. Peterson, D. F. Doak, W. F. Morris, Global Change Biology 25, 775 (2019).

[8] C. P. Nadeau, M. C. Urban, Ecography 42, 1280 (2019).

[9] G. Turesson, Hereditas 3, 100 (1922).

[10] J. Clausen, D. Keck, W. Heisey, Experimental studies on the nature of species III: Environmental responses of climatic races of *Achillea* (1948).

[11] T. J. Kawecki, D. Ebert, Ecology Letters 7, 1225 (2004).

[12] A. M. Wilczek, M. D. Cooper, T. M. Korves, J. Schmitt, Proceedings of the National Academy of Sciences 111, 7906 (2014).

[13] J. B. McGraw, et al., Global Change Biology 21, 3827 (2015).

[14] M. Bontrager, A. L. Angert, Evolution Letters 3, 55 (2019).

[15] J. B. Sambatti, K. J. Rice, Evolution 60, 696 (2006).

[16] C. E. Kindell, A. A. Winn, T. E. Miller, Journal of Ecology 84, 745 (1996).

[17] K. J. Rice, E. E. Knapp, Restoration Ecology 16, 12 (2008).

[18] C. C. Wendling, K. M. Wegner, Proceedings of the Royal Society B: Biological Sciences 282, 20142244 (2015).

[19] C. Cousyn, et al., Proceedings of the National Academy of Sciences 98, 6256 (2001).

[20] J. Reger, M. I. Lind, M. R. Robinson, A. P. Beckerman, Nature Ecology & Evolution 2, 100 (2018).

[21] J. Hereford, The American Naturalist 173, 579 (2009).

[22] F. Blanquart, O. Kaltz, S. L. Nuismer, S. Gandon, Ecology Letters 16, 1195 (2013).

[23] A. H. Halbritter, et al., Journal of Evolutionary Biology 31, 784 (2018).

[24] A. L. Hargreaves, R. M. Germain, M. Bontrager, J. Persi, A. L. Angert, The American Naturalist 195, 395 (2020).

[25] N. J. Kooyers, et al., The American Naturalist 194, 541 (2019).

[26] K. E. Barton, C. Jones, K. F. Edwards, A. B. Shiels, T. Knight, Journal of Ecology 108, 1540 (2020).

[27] C. A. Knight, et al., Molecular Ecology 15, 1229 (2006).

[28] E. Allan, J. R. Pannell, Oikos 118, 1053 (2009).

[29] A. Mead, et al., Molecular Ecology 28, 5248 (2019).

[30] J. M. Levine, A. K. McEachern, C. Cowan, Ecology 92, 2236 (2011).

[31] M. L. LaForgia, S. P. Harrison, A. M. Latimer, Ecology 101, e03022 (2020).

[32] D. J. Young, et al., Ecology 100, e02571 (2019).

[33] Z. Munzbergová, V. Hadincová, H. Skálová, V. Vandvik, Journal of Ecology 105, 1358 (2017).

[34] M. M. Kling, S. L. Auer, P. J. Comer, D. D. Ackerly, H. Hamilton, Global Change Biology 26, 2798 (2020).

[35] M. C. Urban, et al., Proceedings of the National Academy of Sciences 117, 17482 (2020).

[36] M. Bontrager, et al., BioRxiv (2020).

[37] L. Tyberghein, et al., Global Ecology and Biogeography 21, 272 (2012).

[38] R. Leimu, M. Fischer, PloS One 3, e4010 (2008).

[39] J. A. Lee-Yaw, et al., Ecology Letters 19, 710 (2016).

[40] A. M. Oduor, R. Leimu, M. Kleunen, Journal of Ecology 104, 957 (2016).

[41] A. L. Gibson, E. K. Espeland, V. Wagner, C. R. Nelson, Evolutionary Applications 9, 1219 (2016).

[42] T. Wang, A. Hamann, D. Spittlehouse, C. Carroll, PLoS One 11, e0156720 (2016).

[43] A. Hamann, T. Wang, D. L. Spittlehouse, T. Q. Murdock, Bulletin of the American Meteorological Society 94, 1307 (2013).

[44] T. D. Mitchell, P. D. Jones, International Journal of Climatology 25, 693 (2005).

[45] I. Harris, P. D. Jones, T. J. Osborn, D. H. Lister, International journal of climatology 34, 623 (2014).

[46] C. Korner, Alpine plant life: functional plant ecology of high mountain ecosystems (Springer Science & Business Media, 2003).

[47] R. K. Colwell, G. Brehm, C. L. Cardeluás, A. C. Gilman, J. T. Longino, Science 322, 258 (2008).

[48] G. Martin, T. Lenormand, Evolution 60, 2413 (2006).

[49] R. Levins, Evolution in changing environments: some theoretical explorations (Princeton University Press, 1968).

[50] M. Lynch, B. Walsh, Genetics and analysis of quantitative traits (Sinauer Sunderland, MA, 1998).

[51] Stan Development Team, RStan: the R interface to Stan (2020). R package version 2.21.2.

[52] A. Gelman, J. B. Carlin, H. S. Stern, D. B. Rubin, Bayesian Data Analysis (Chapman and Hall/CRC, 2004), second edn.

[53] P.-C. Brkner, Journal of Statistical Software 80, 1 (2017).

[54] B. Carpenter, et al., Journal of Statistical Software, Articles 76, 1 (2017).

[55] A. Gelman, D. B. Rubin, Statistical Science 7, 457 (1992).

[56] P.-C. Brkner, J. Gabry, M. Kay, posterior: Tools for Working with Posterior Distributions (2020). R package version 0.0.3.

[57] J. Gabry, T. Mahr, bayesplot: Plotting for Bayesian Models (2020). R package version 1.7.2.

