## Supplemental Area and Trace plots for "Climate warming weakens local adaptation"

### Model A

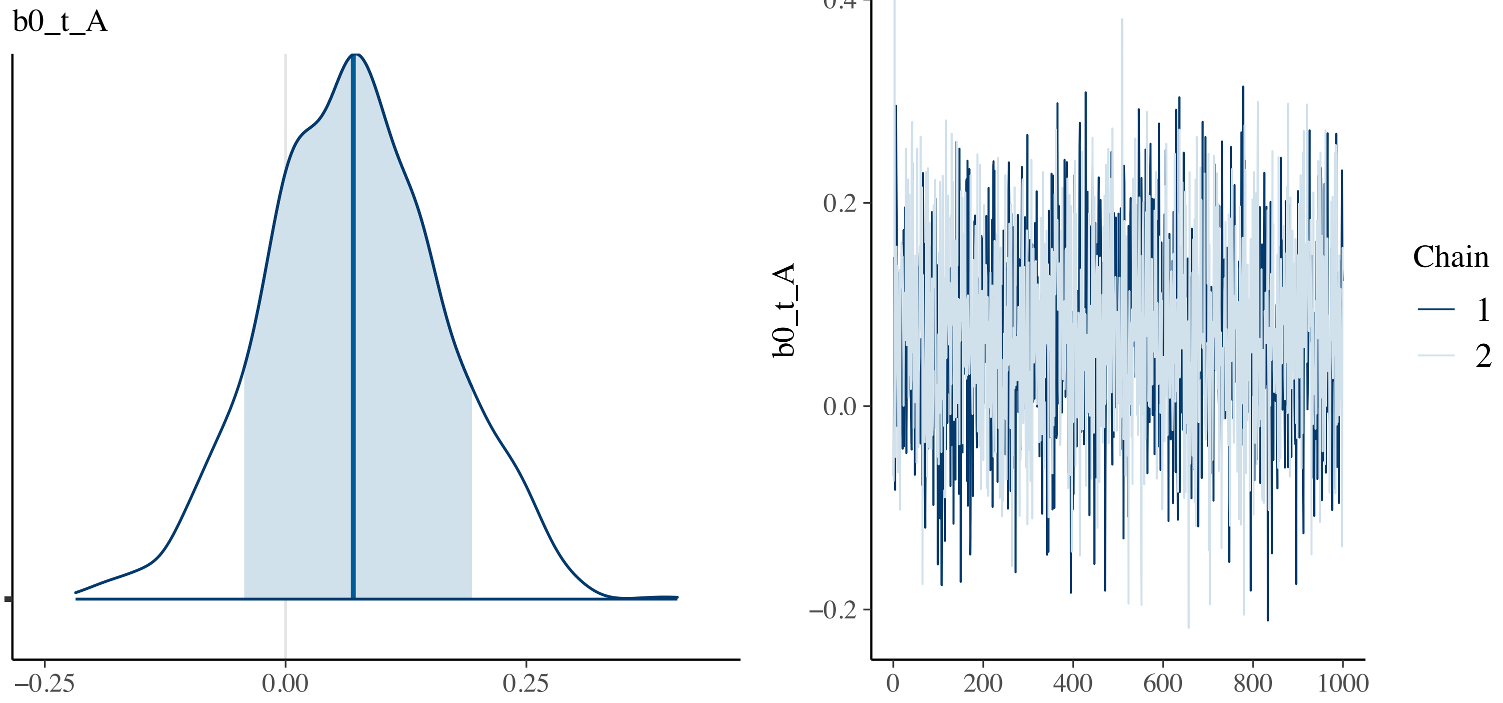

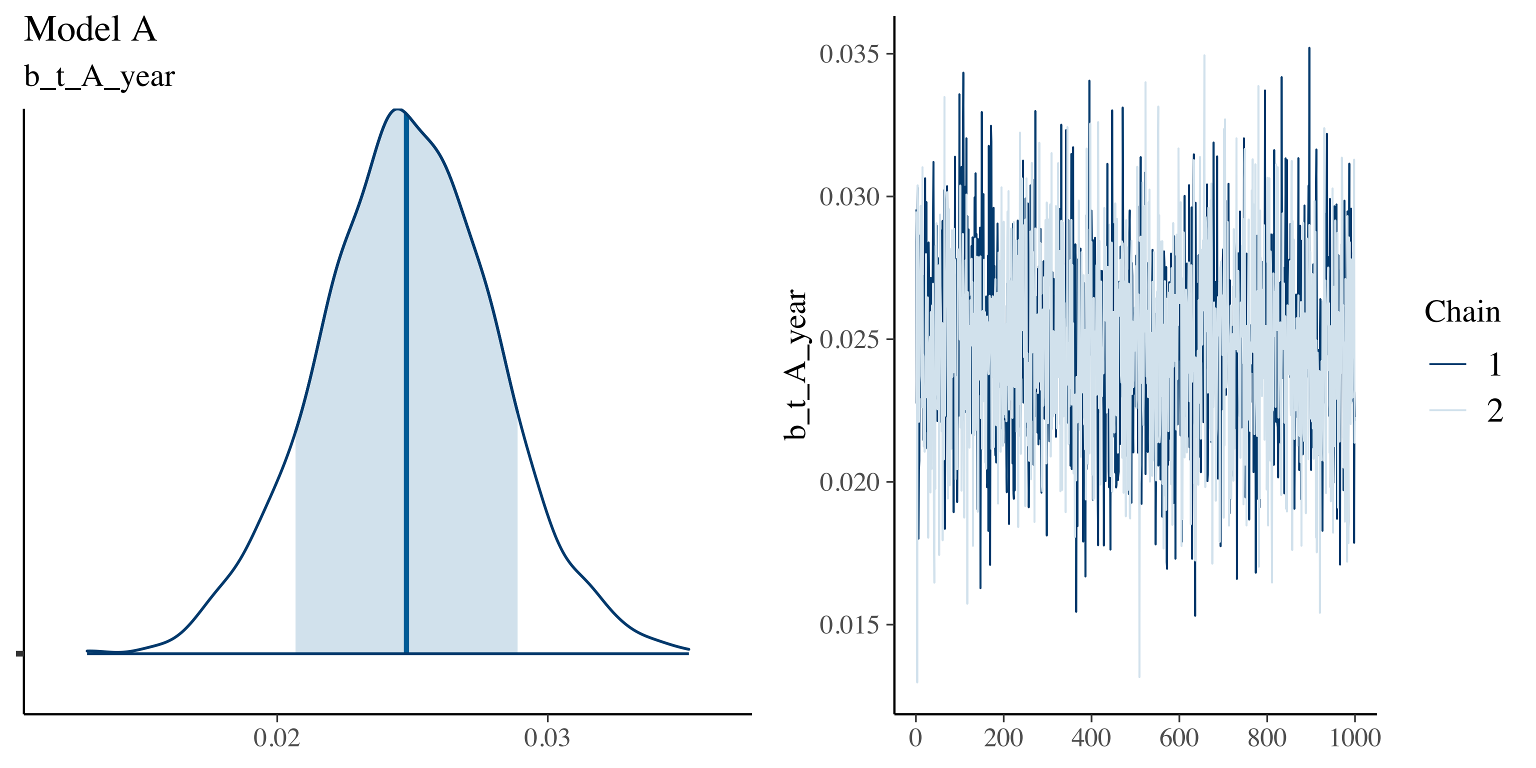
Model A
b_t_A_year
0.02
0.03
0.035
0.030
0.025
0.020
0.015
b_t_A_year
0
200
400
600
800
1000
Chain
1
2

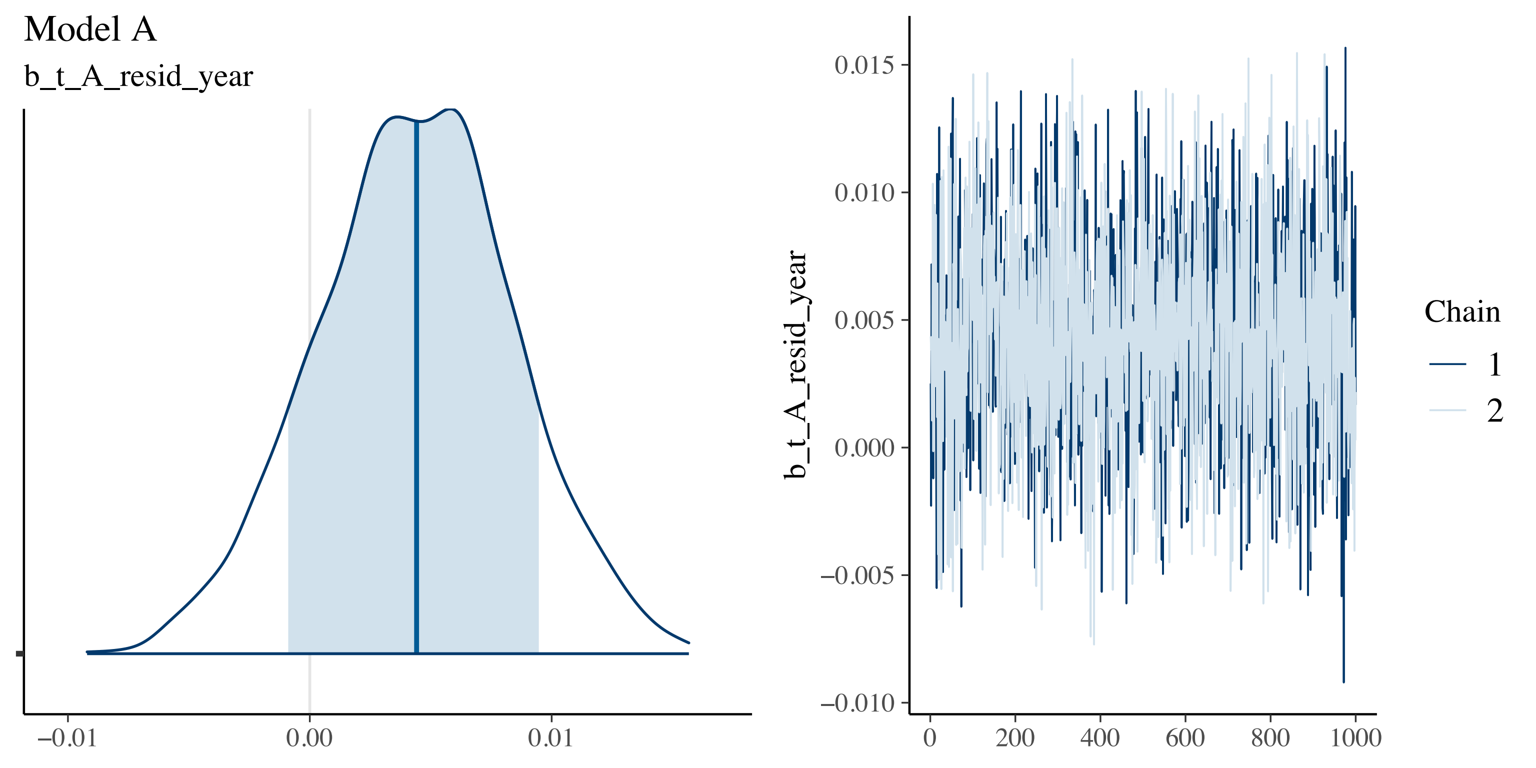
Model A
b_t_A_resid_year
−0.01
0.00
0.01
b_t_A_resid_year
0.015
0.010
0.005
0.000
−0.005
−0.010
0
200
400
600
800
1000
Chain
1
2

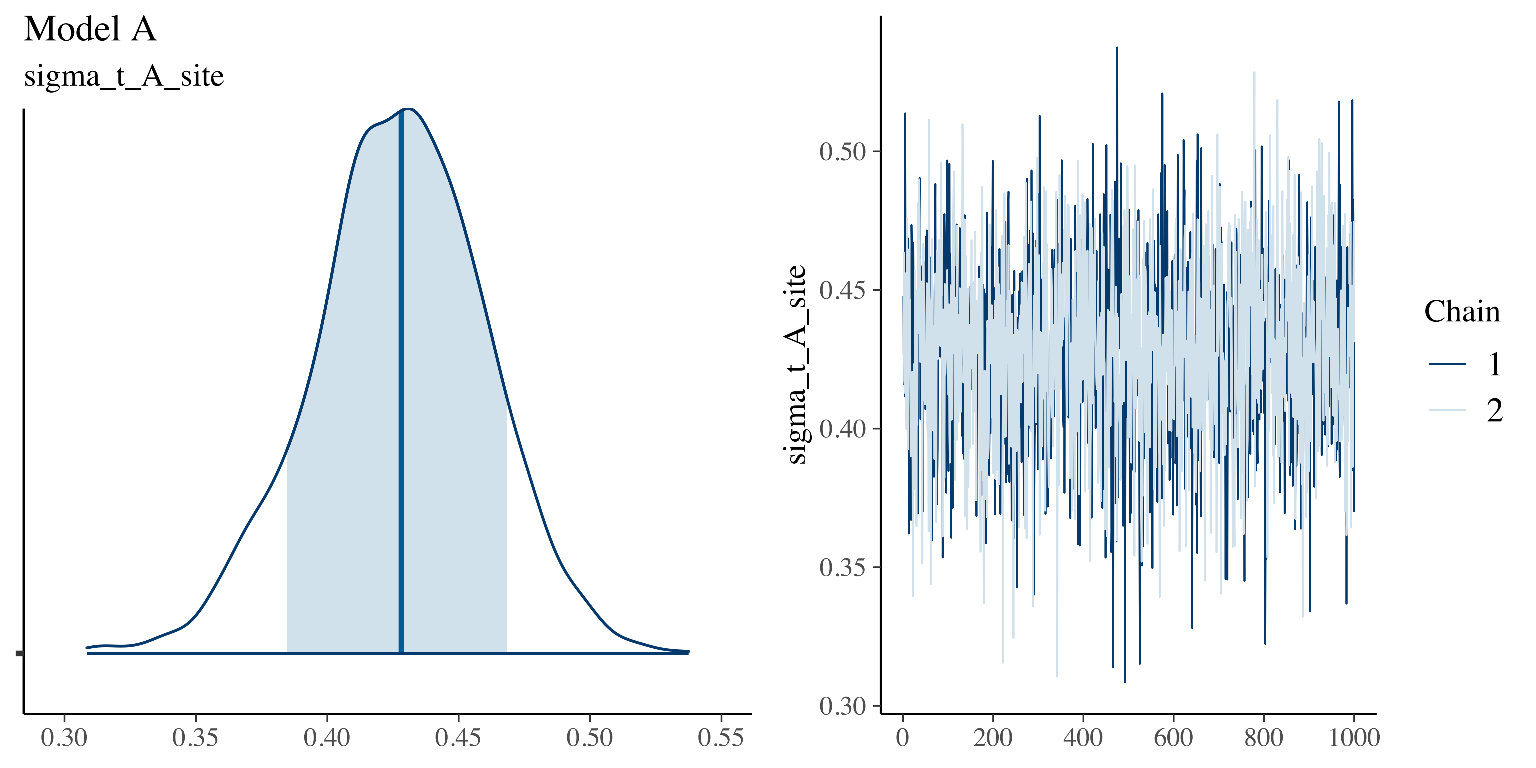

Model A
sigma_t_A_site
0.30
0.35
0.40
0.45
0.50
0.55
sigma_t_A_site
0.30
0.35
0.40
0.45
0.50
0
200
400
600
800
1000
Chain
1
2

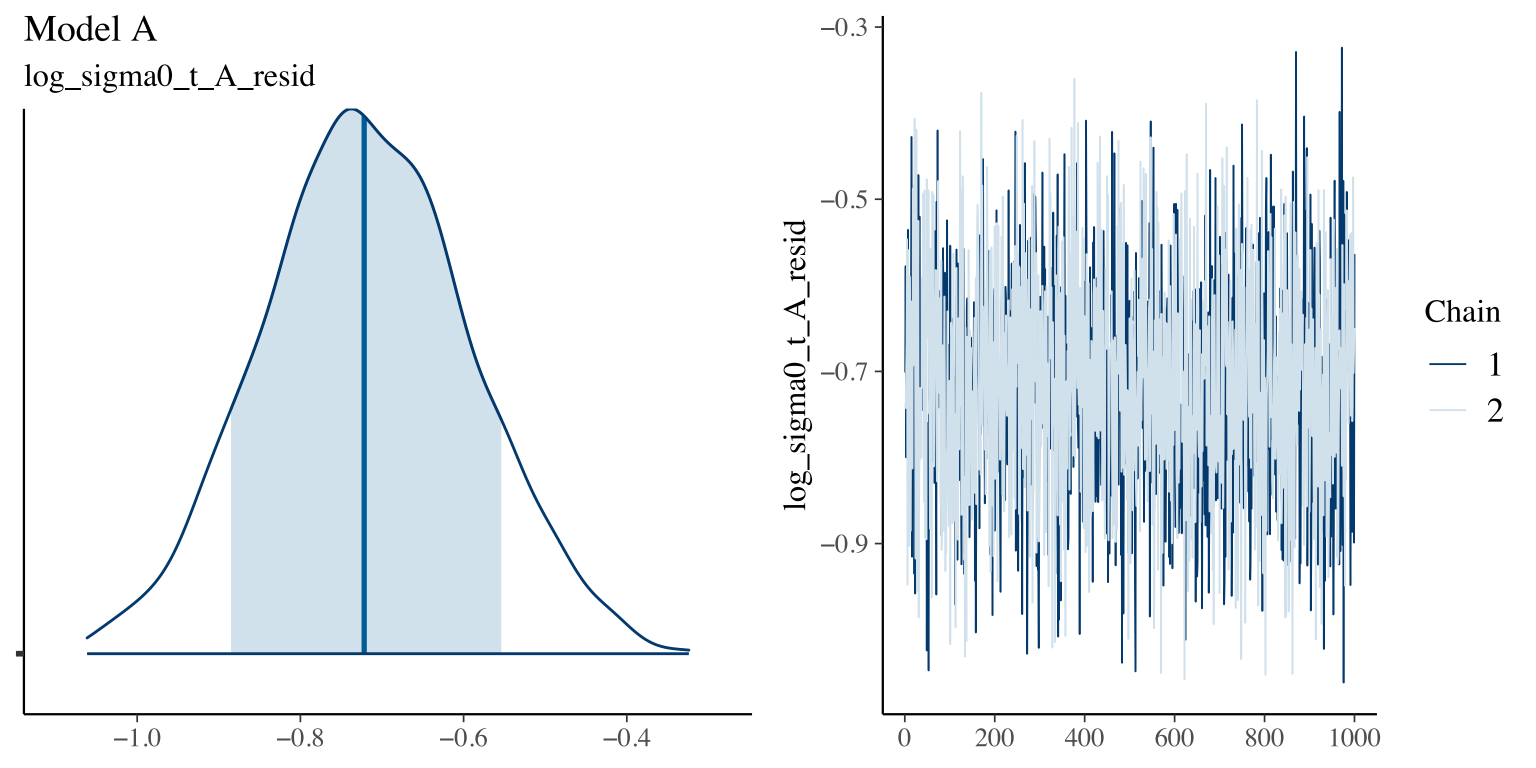

Model A
log_sigma0_t_A_resid
−1.0
−0.8
−0.6
−0.4
−0.3
−0.5
−0.7
−0.9
log_sigma0_t_A_resid
0
200
400
600
800
1000
Chain
1
2

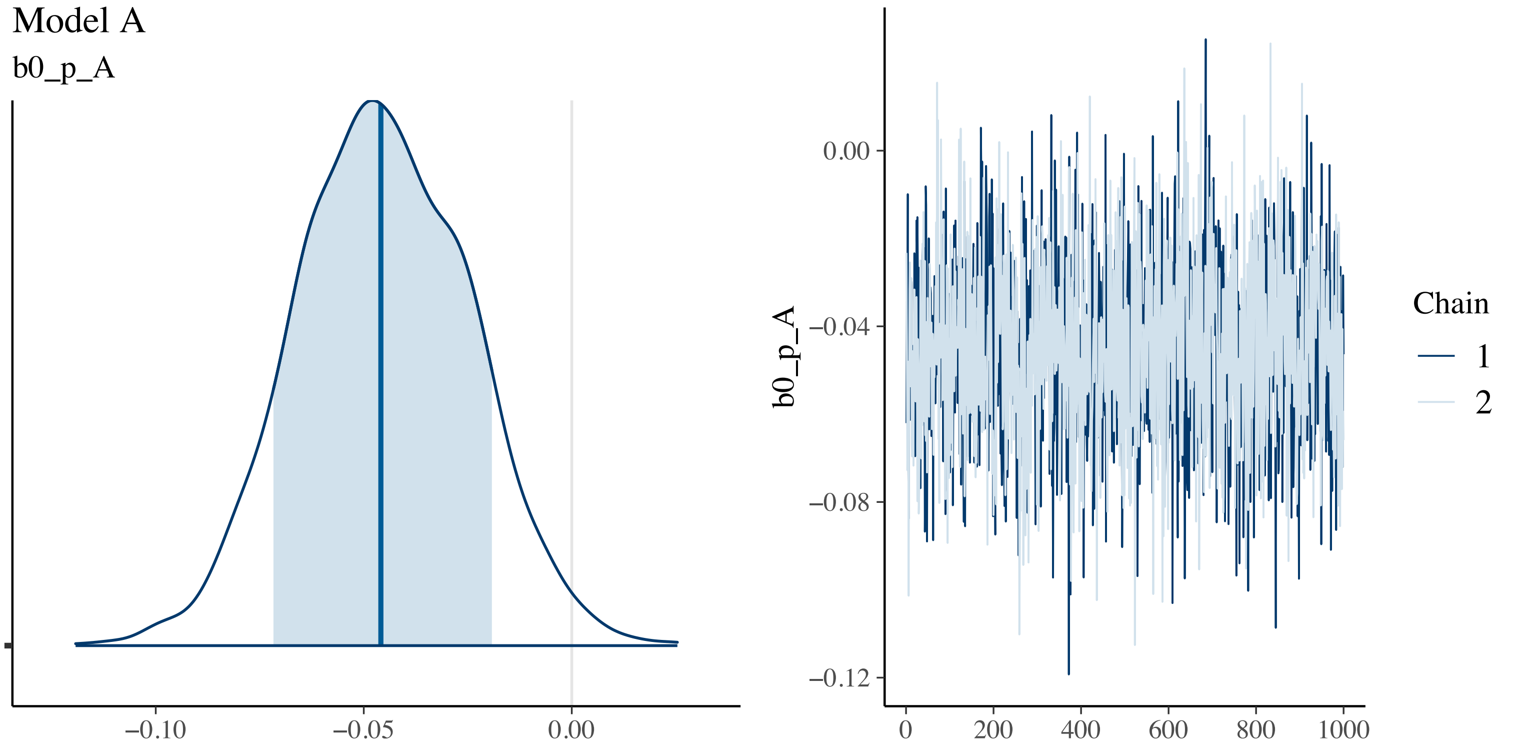

Model A
b0_p_A
−0.10
−0.05
0.00
b0_p_A
0.00
−0.04
−0.08
−0.12
0
200
400
600
800
1000
Chain
1
2

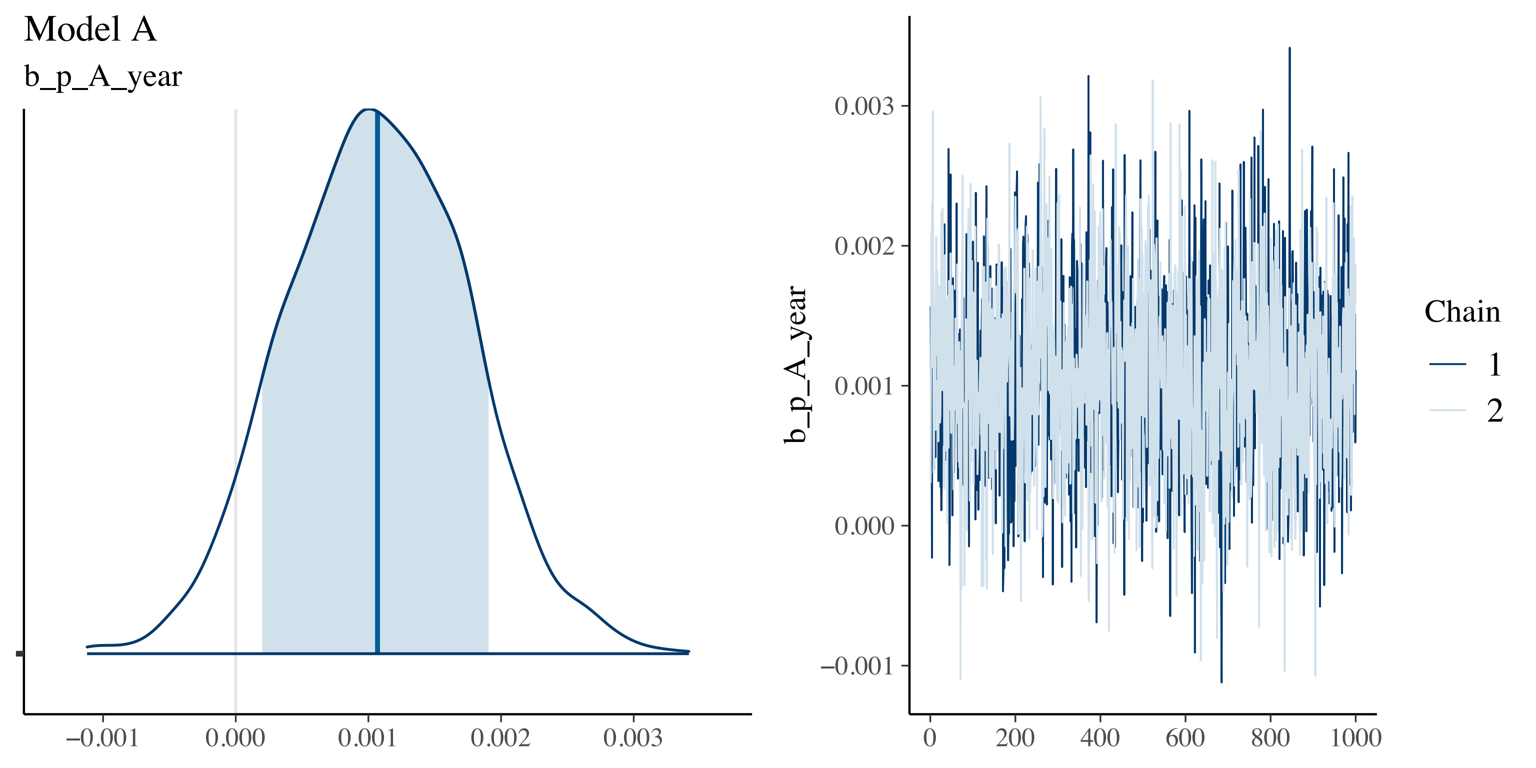

Model A
b_p_A_year
−0.001
0.000
0.001
0.002
0.003
b_p_A_year
0.003
0.002
0.001
0.000
−0.001
0
200
400
600
800
1000
Chain
1
2

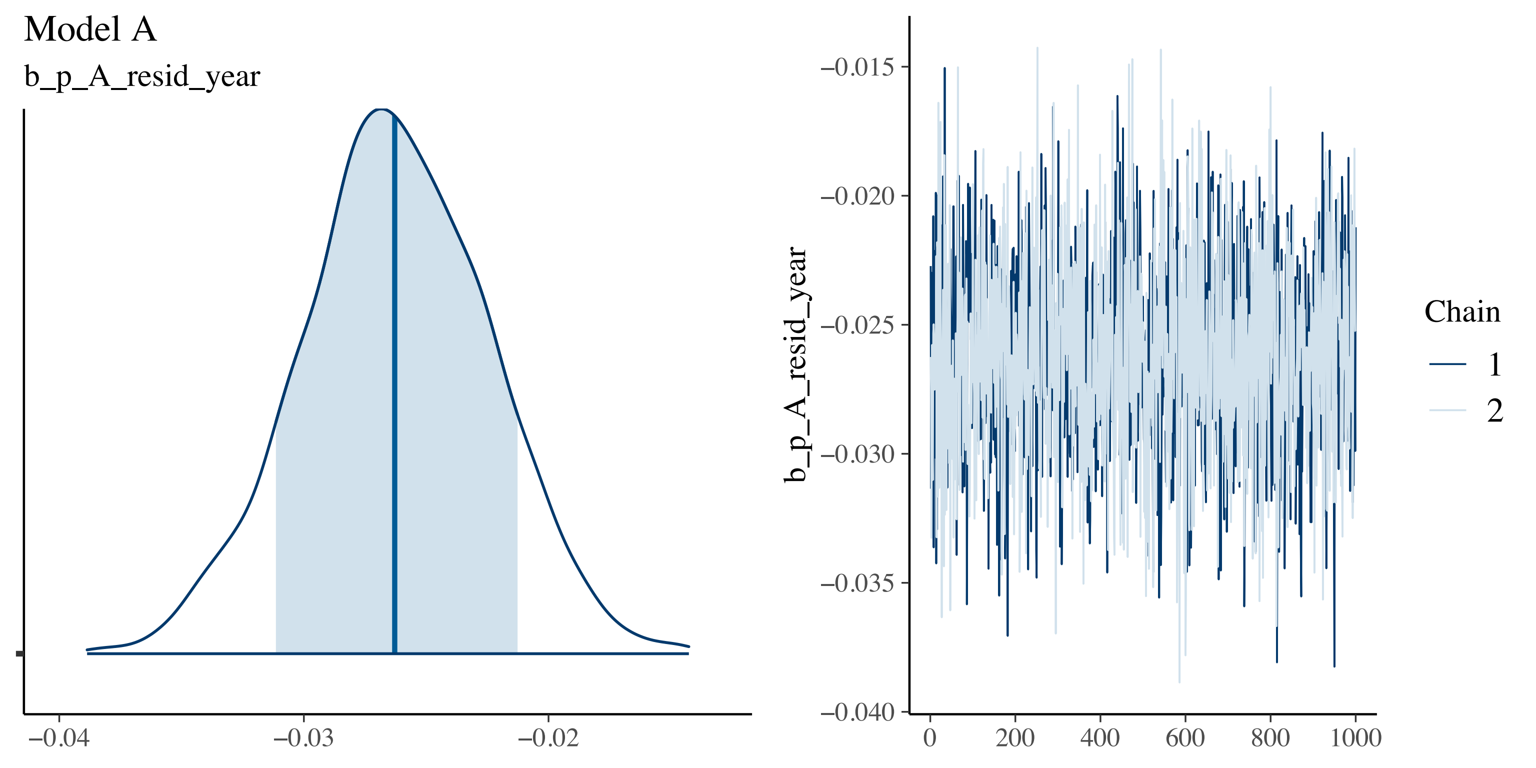

Model A
b_p_A_resid_year
−0.04
−0.03
−0.02
b_p_A_resid_year
−0.015
−0.020
−0.025
−0.030
−0.035
−0.040
0
200
400
600
800
1000
Chain
1
2

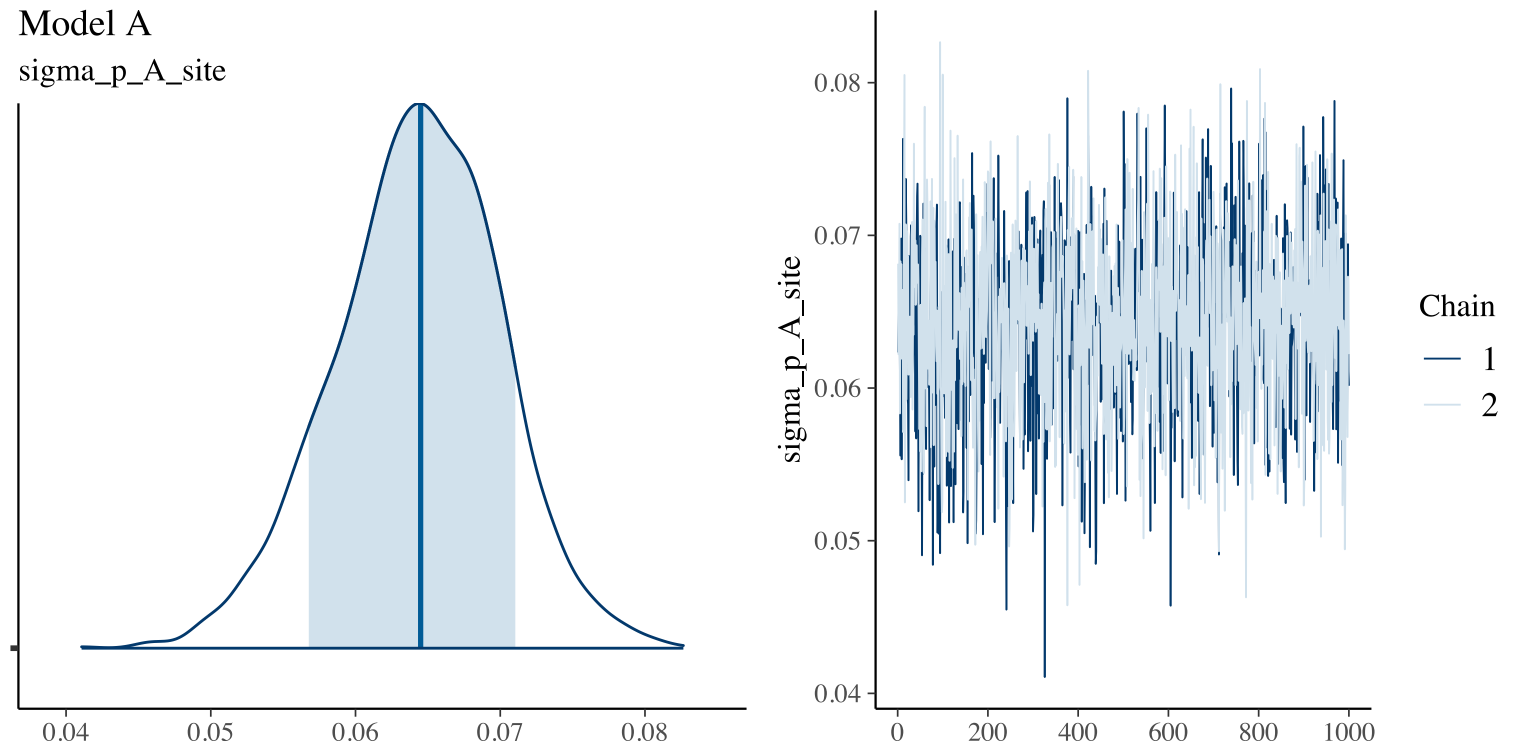

Model A
sigma_p_A_site
0.04
0.05
0.06
0.07
0.08
sigma_p_A_site
0
200
400
600
800
1000
Chain
1
2

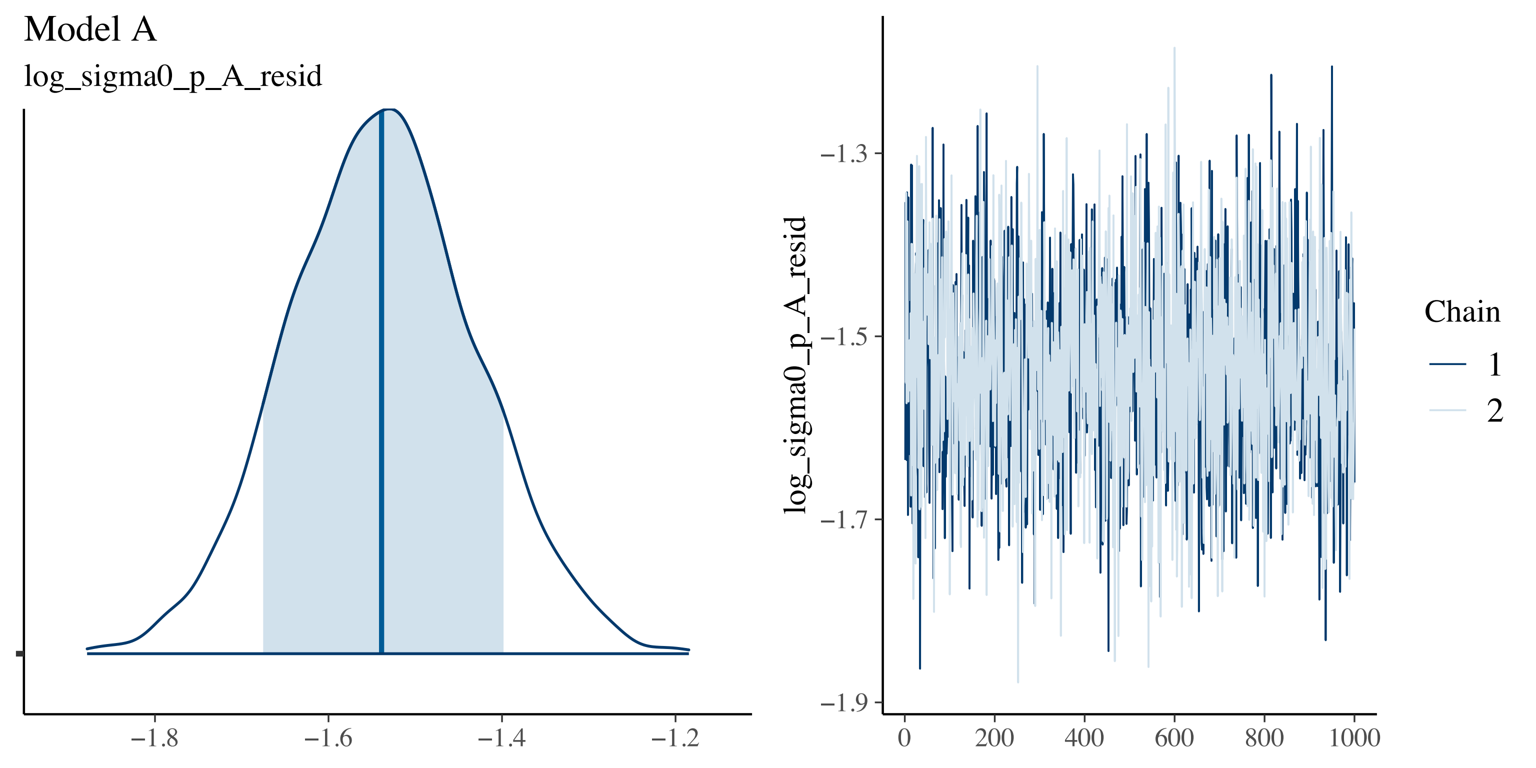
Model A
log_sigma0_p_A_resid
−1.8
−1.6
−1.4
−1.2
log_sigma0_p_A_resid
−1.3
−1.5
−1.7
−1.9
0
200
400
600
800
1000
Chain
1
2

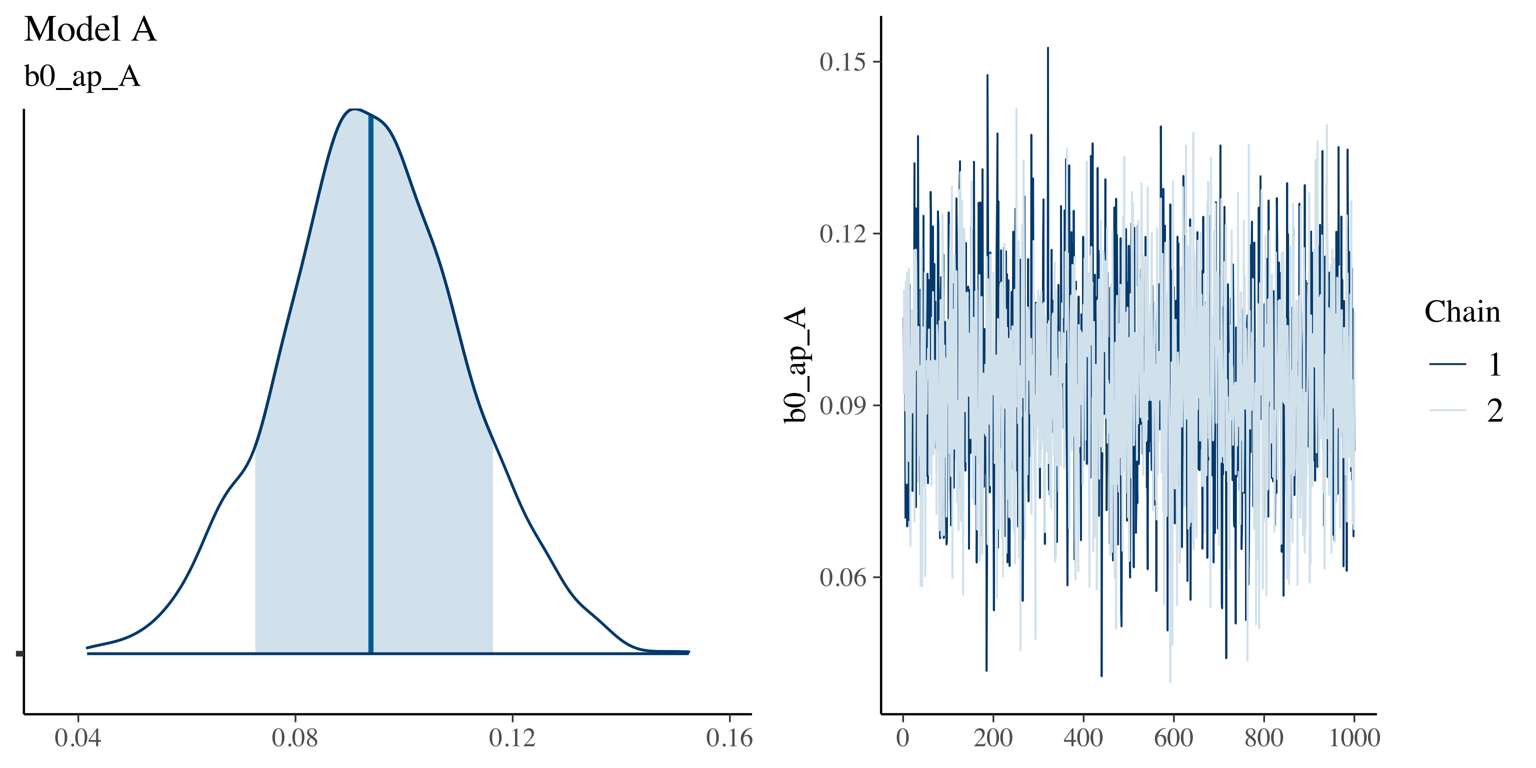

Model A
b0_ap_A
0.04
0.08
0.12
0.16
b0_ap_A
0.15
0.12
0.09
0.06
0
200
400
600
800
1000
Chain
1
2

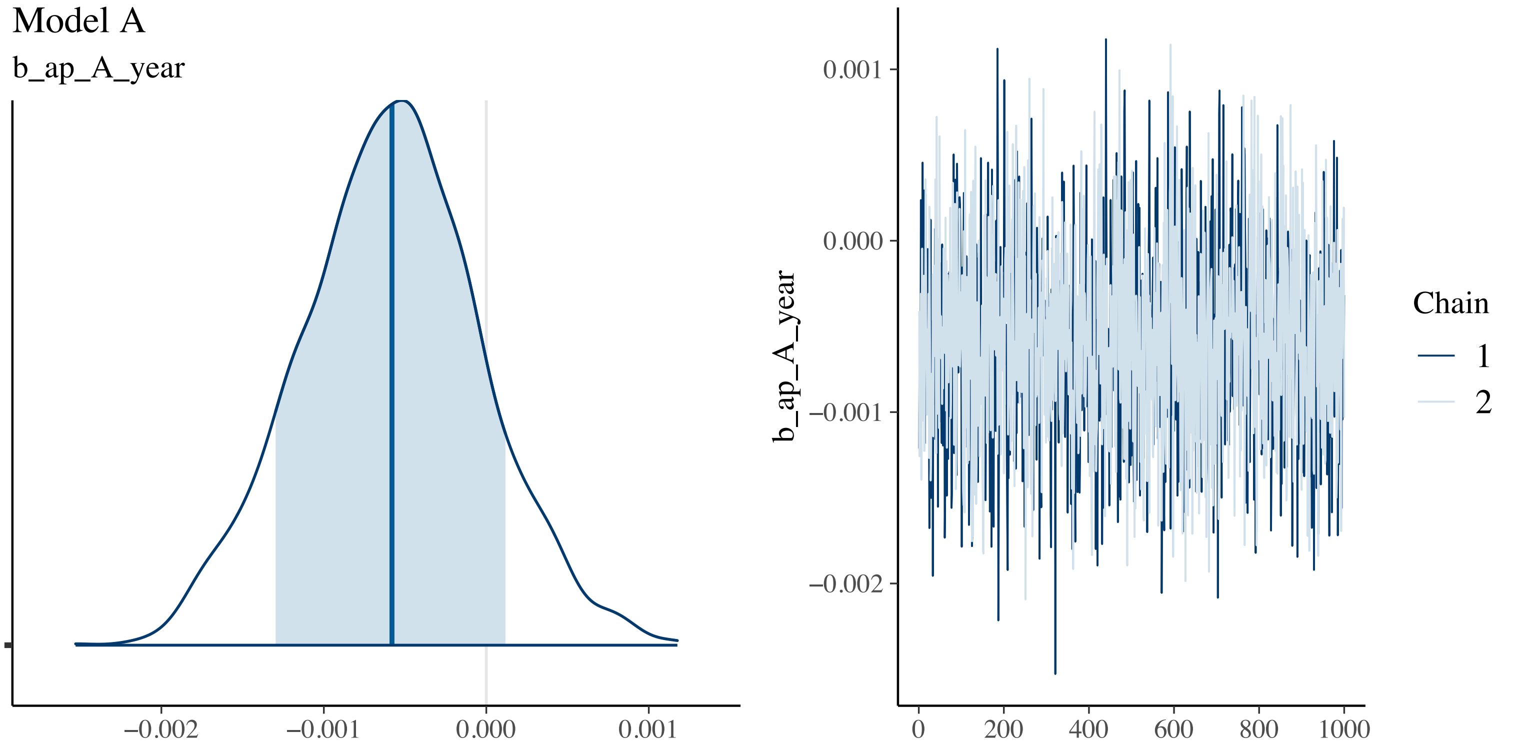

Model A
b_ap_A_year
−0.002
−0.001
0.000
0.001
b_ap_A_year
0.001
0.000
−0.001
−0.002
0
200
400
600
800
1000
Chain
1
2

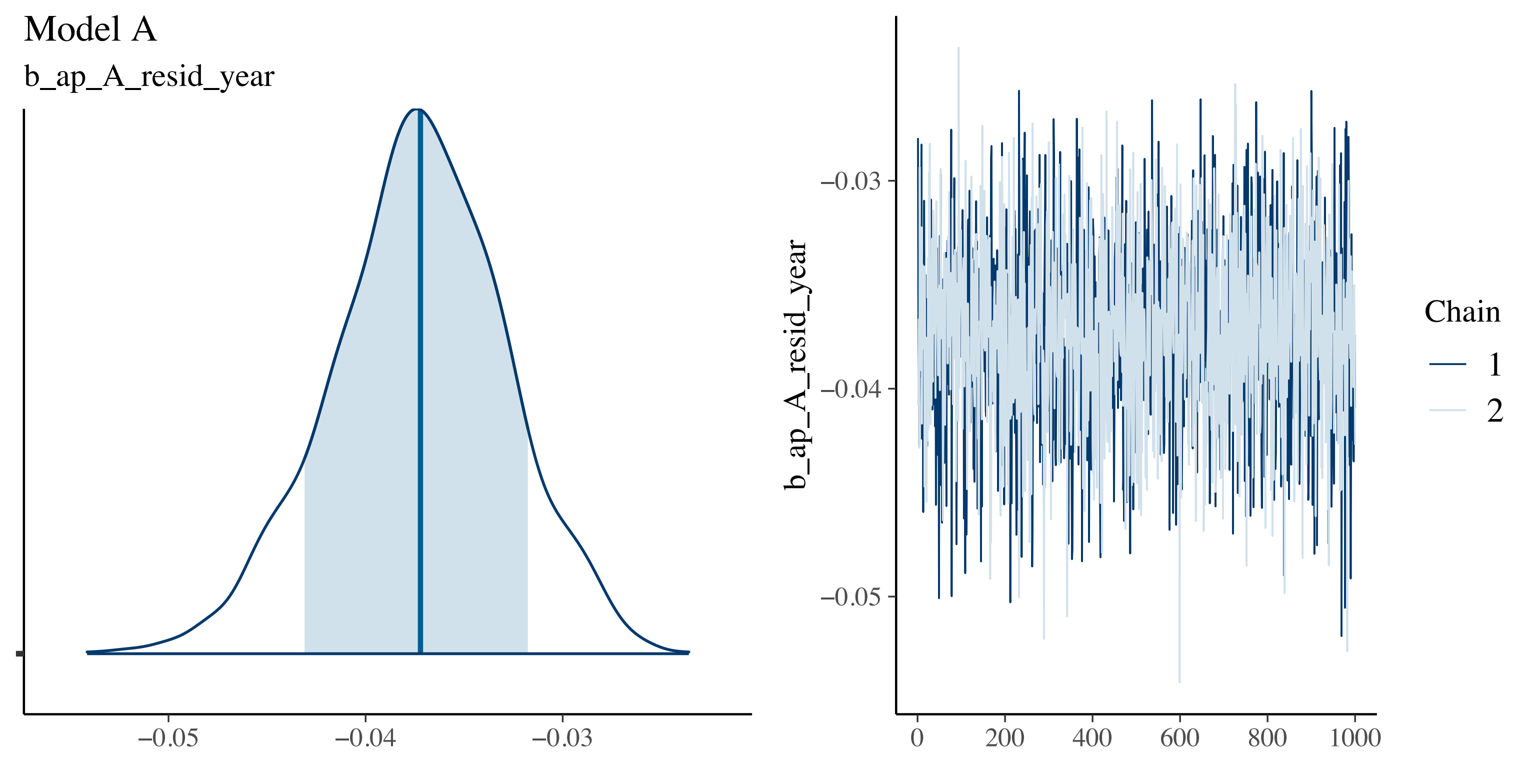

Model A
b_ap_A_resid_year
−0.05
−0.04
−0.03
b_ap_A_resid_year
−0.03
−0.04
−0.05
0
200
400
600
800
1000
Chain
1
2

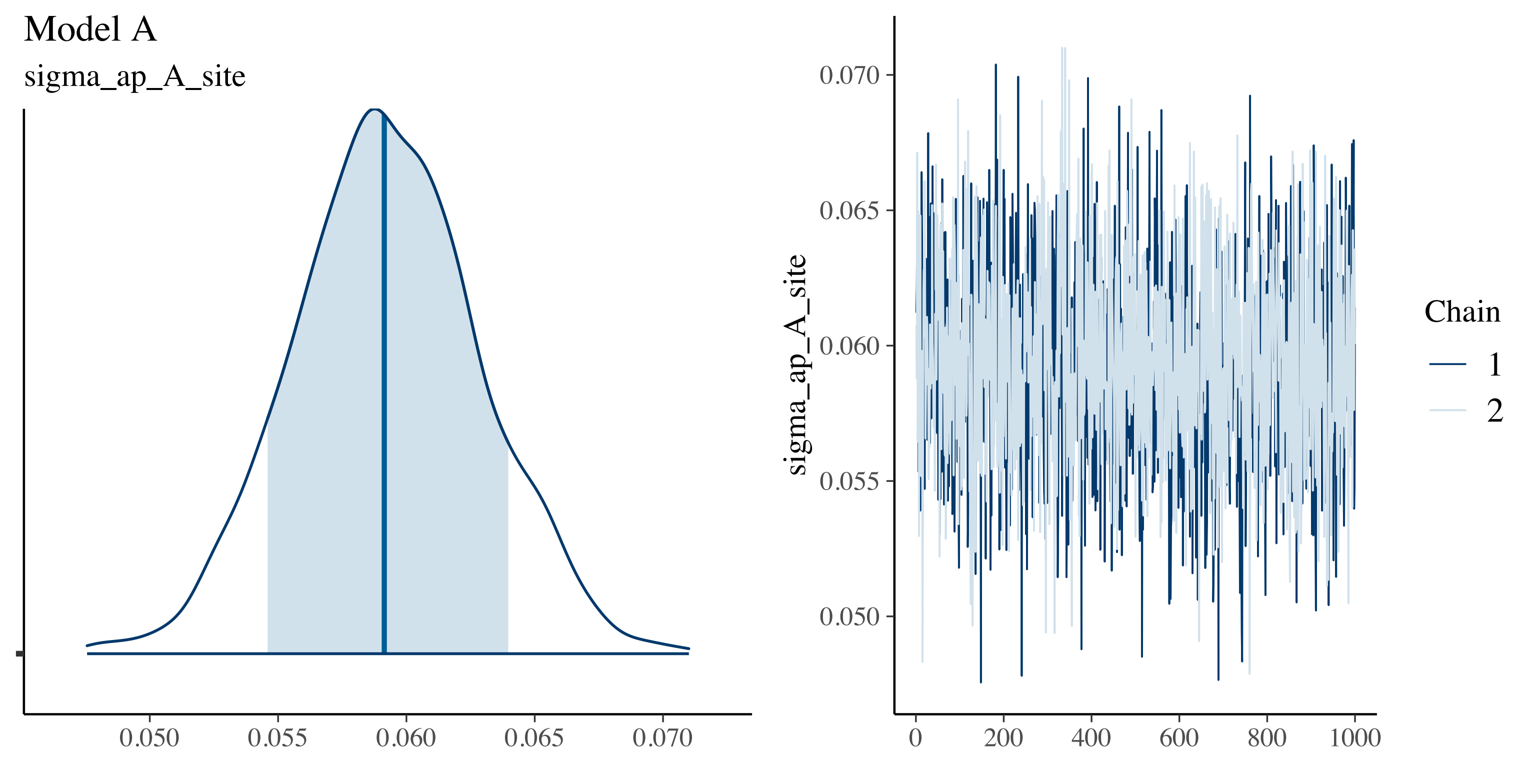

Model A
sigma_ap_A_site
0.050
0.055
0.060
0.065
0.070
sigma_ap_A_site
0.070
0.065
0.060
0.055
0.050
0
200
400
600
800
1000
Chain
1
2

Model A

log_sigma0_ap_A_resid

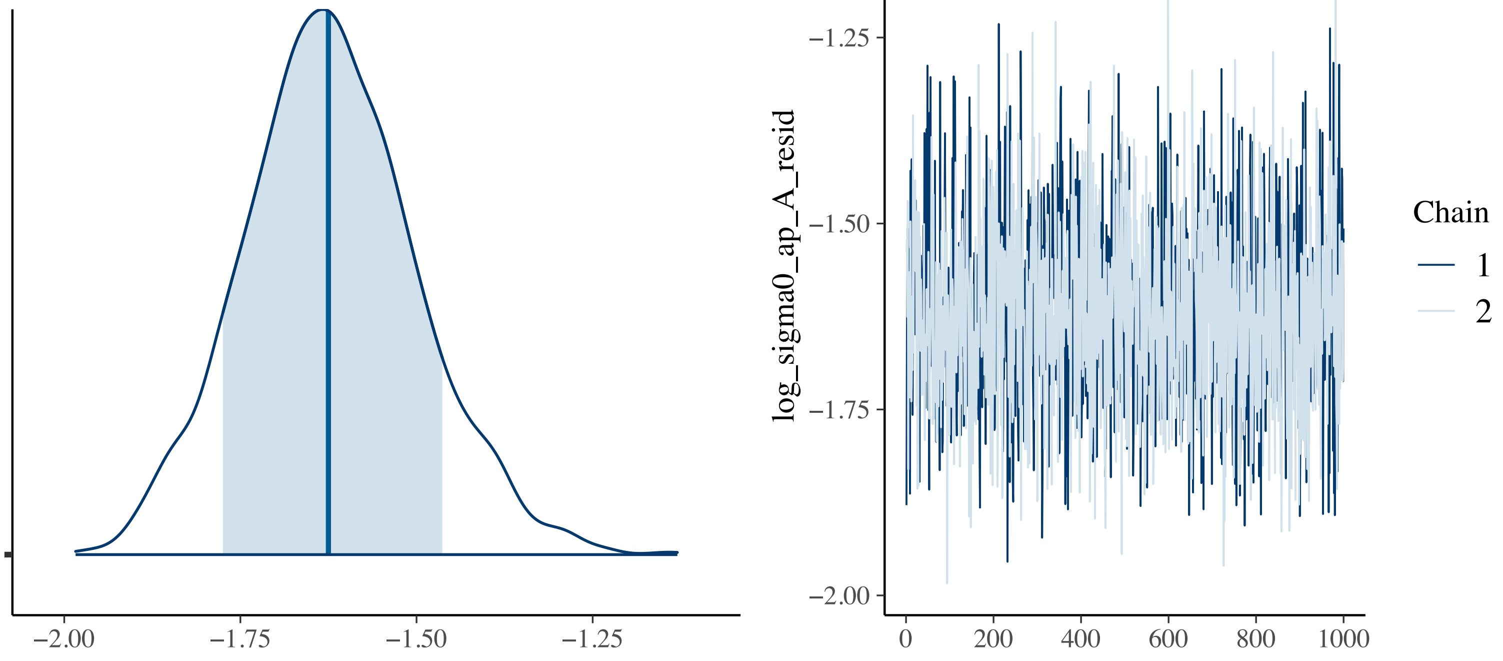

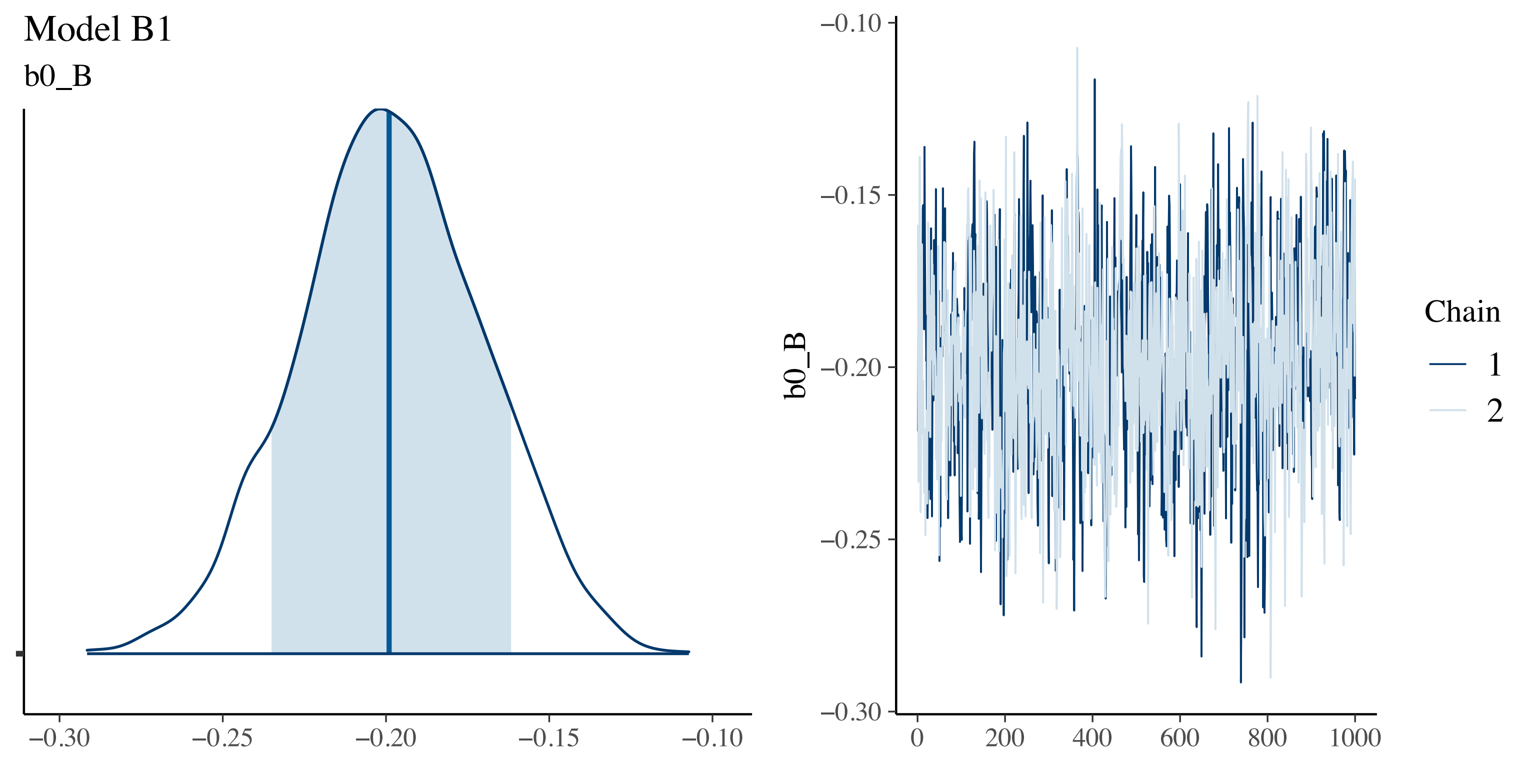

Model B1
b0_B
−0.30
−0.25
−0.20
−0.15
−0.10
b0_B
0
200
400
600
800
1000
Chain
1
2

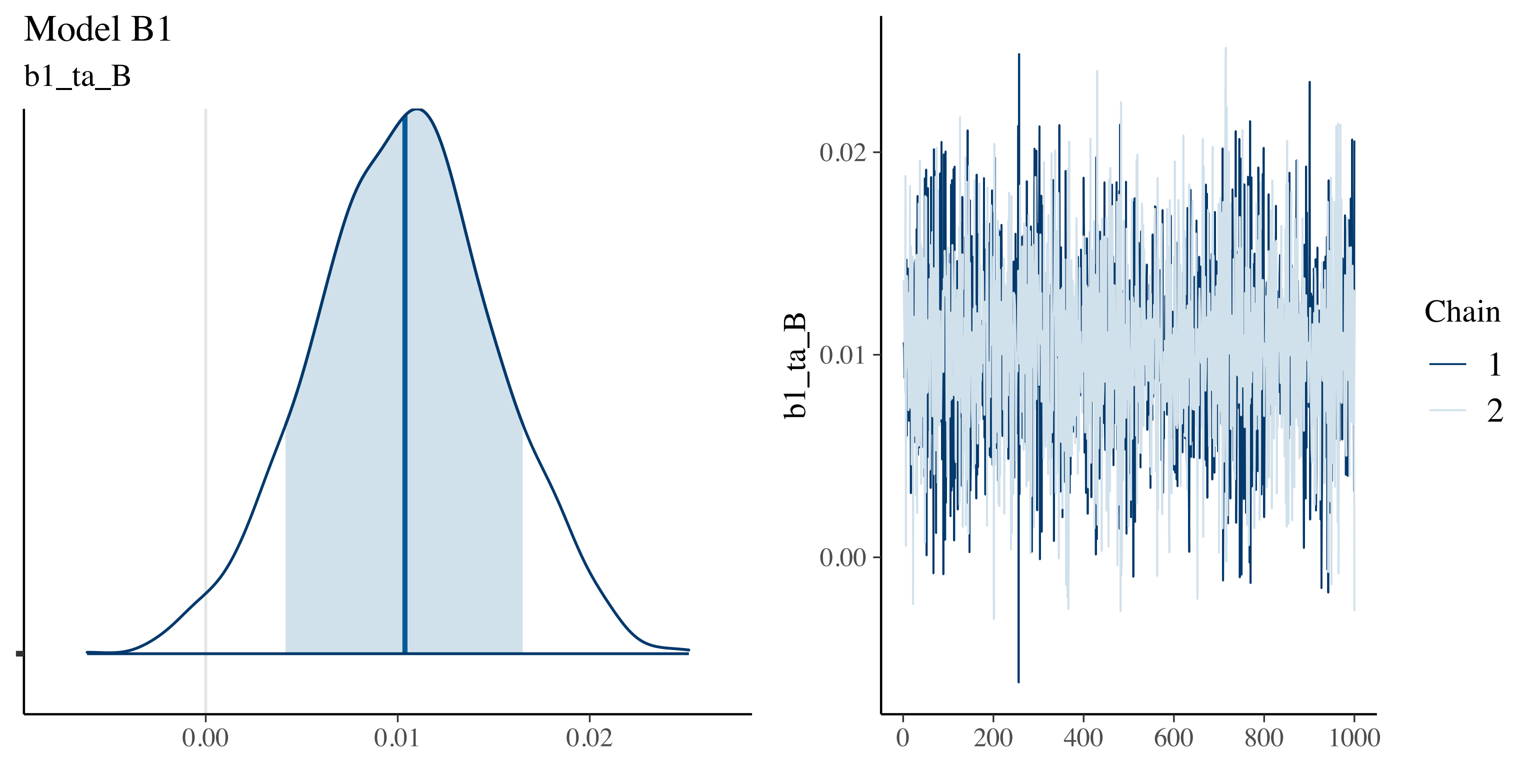
Model B1
b1_ta_B
0.00
0.01
0.02
b1_ta_B
0.02
0.01
0.00
0
200
400
600
800
1000
Chain
1
2

Model B1

b2_ta_B

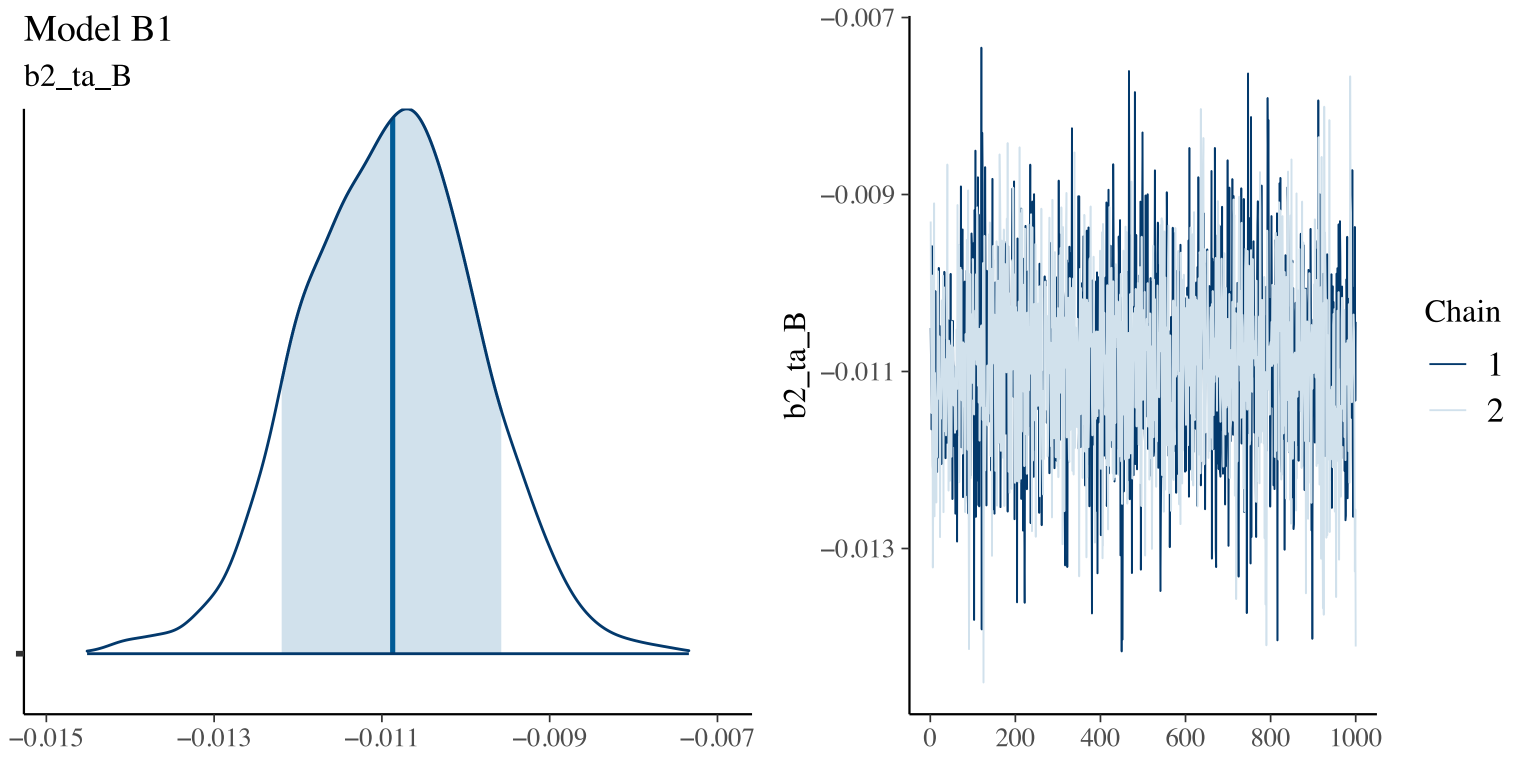

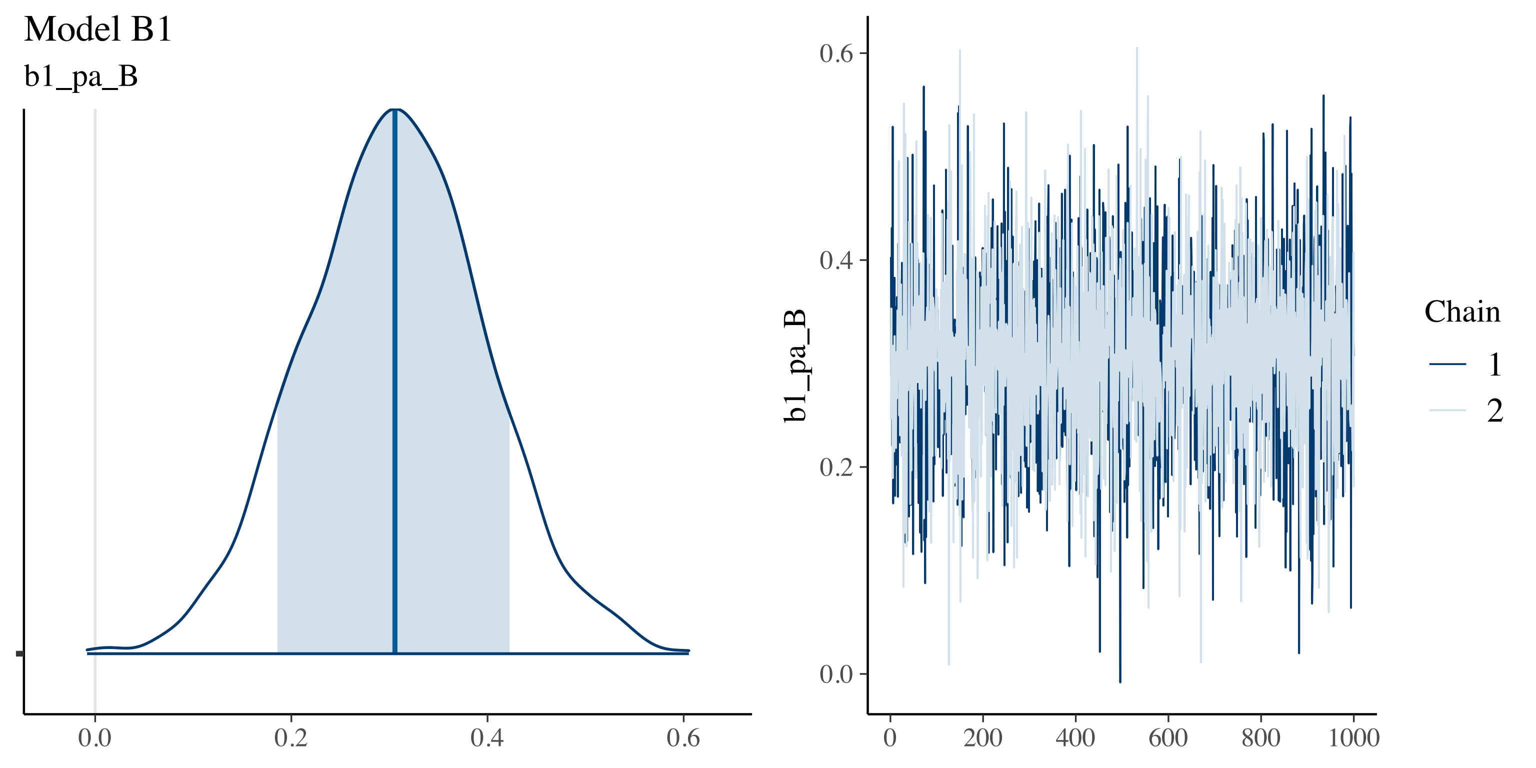
Model B1
b1_pa_B
0.0
0.2
0.4
0.6
b1_pa_B
0.0
0.2
0.4
0.6
0
200
400
600
800
1000
Chain
1
2

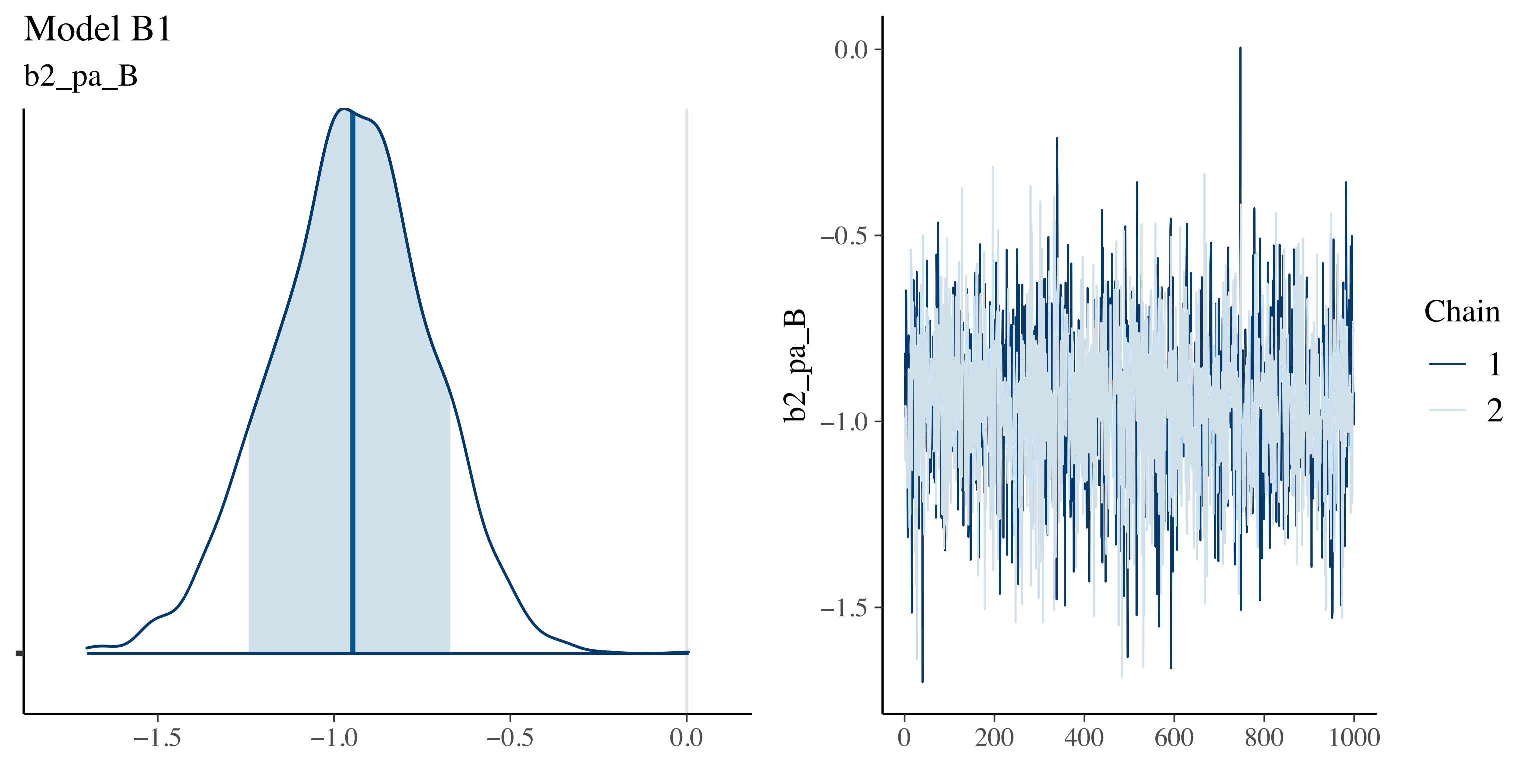
Model B1
b2_pa_B
−1.5
−1.0
−0.5
0.0
b2_pa_B
0.0
−0.5
−1.0
−1.5
0
200
400
600
800
1000
Chain
1
2

Model B1

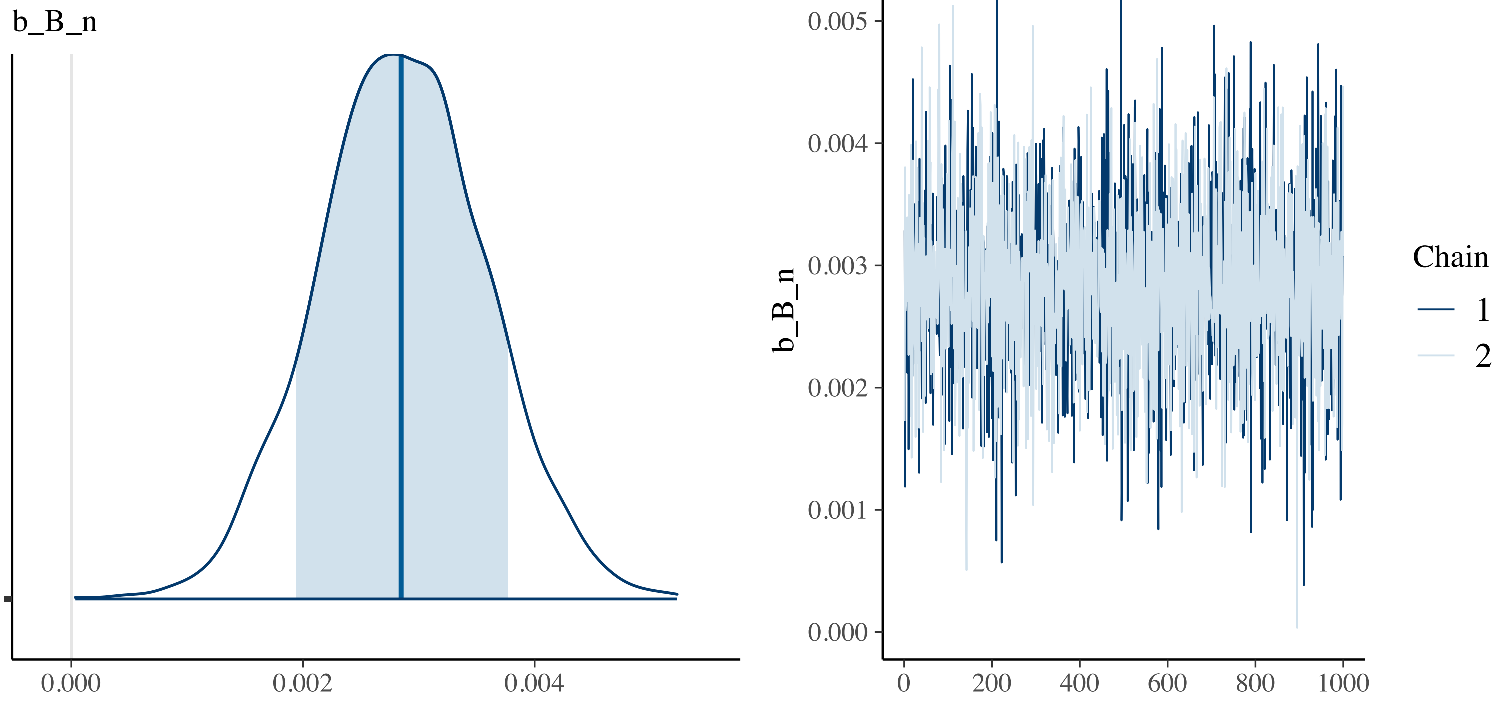
b_B_n
0.000
0.002
0.004
0.005
0.004
0.003
0.002
0.001
0.000
b_B_n
0
200
400
600
800
1000
Chain
1
2

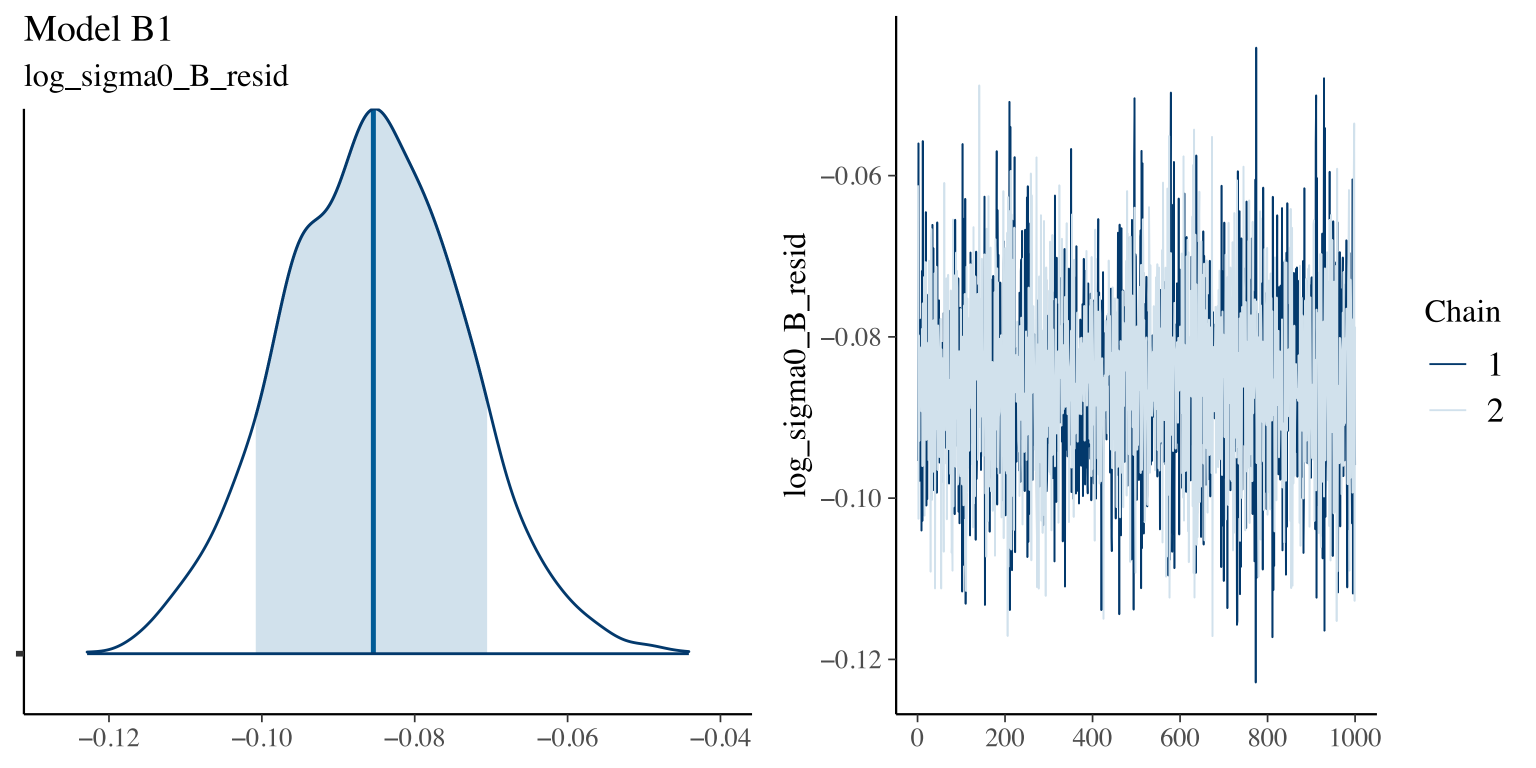
Model B1
log_sigma0_B_resid
−0.12
−0.10
−0.08
−0.06
−0.04
log_sigma0_B_resid
−0.06
−0.08
−0.10
−0.12
0
200
400
600
800
1000
Chain
1
2

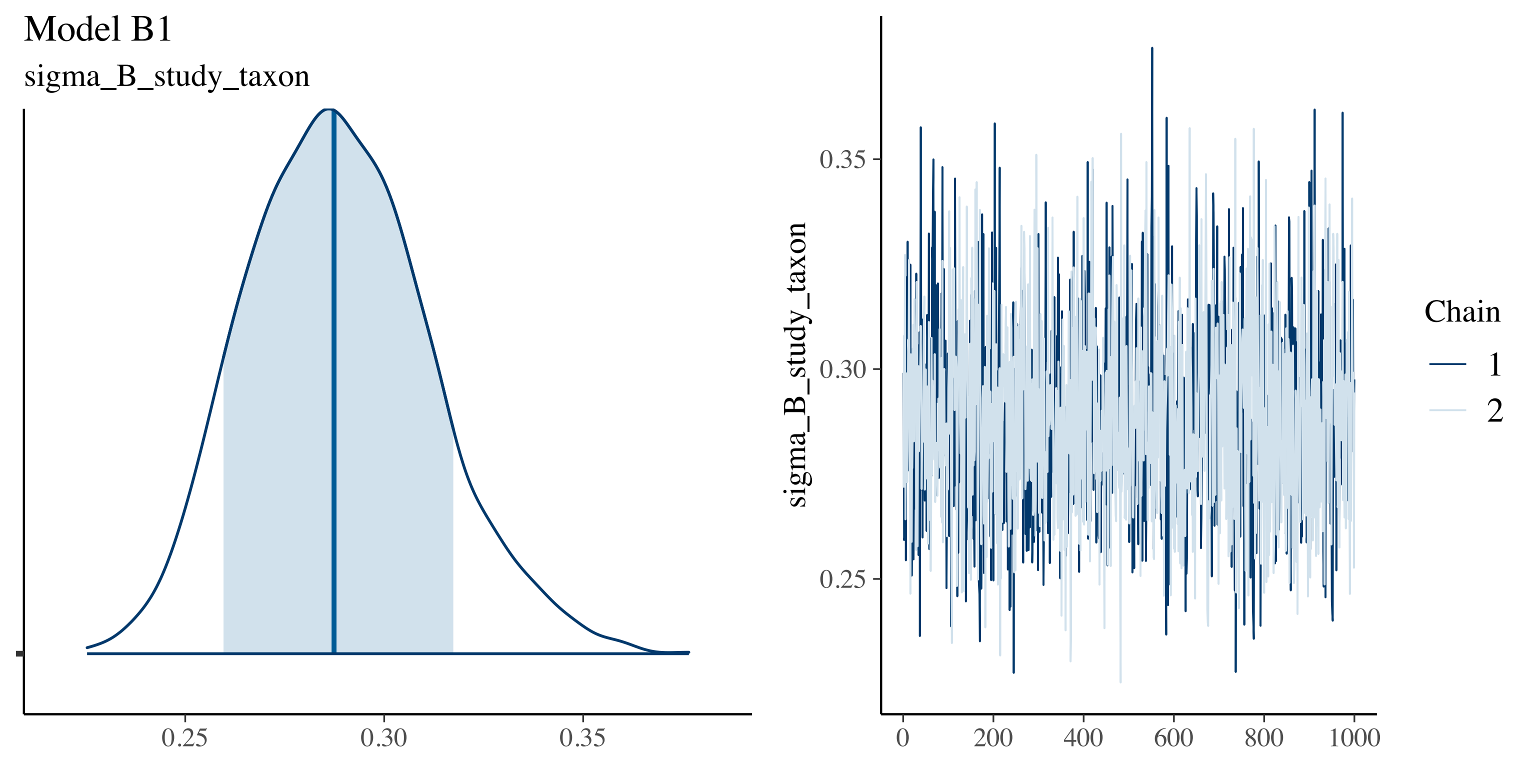

Model B1
sigma_B_study_taxon
0.25
0.30
0.35
sigma_B_study_taxon
0.35
0.30
0.25
0
200
400
600
800
1000
Chain
1
2

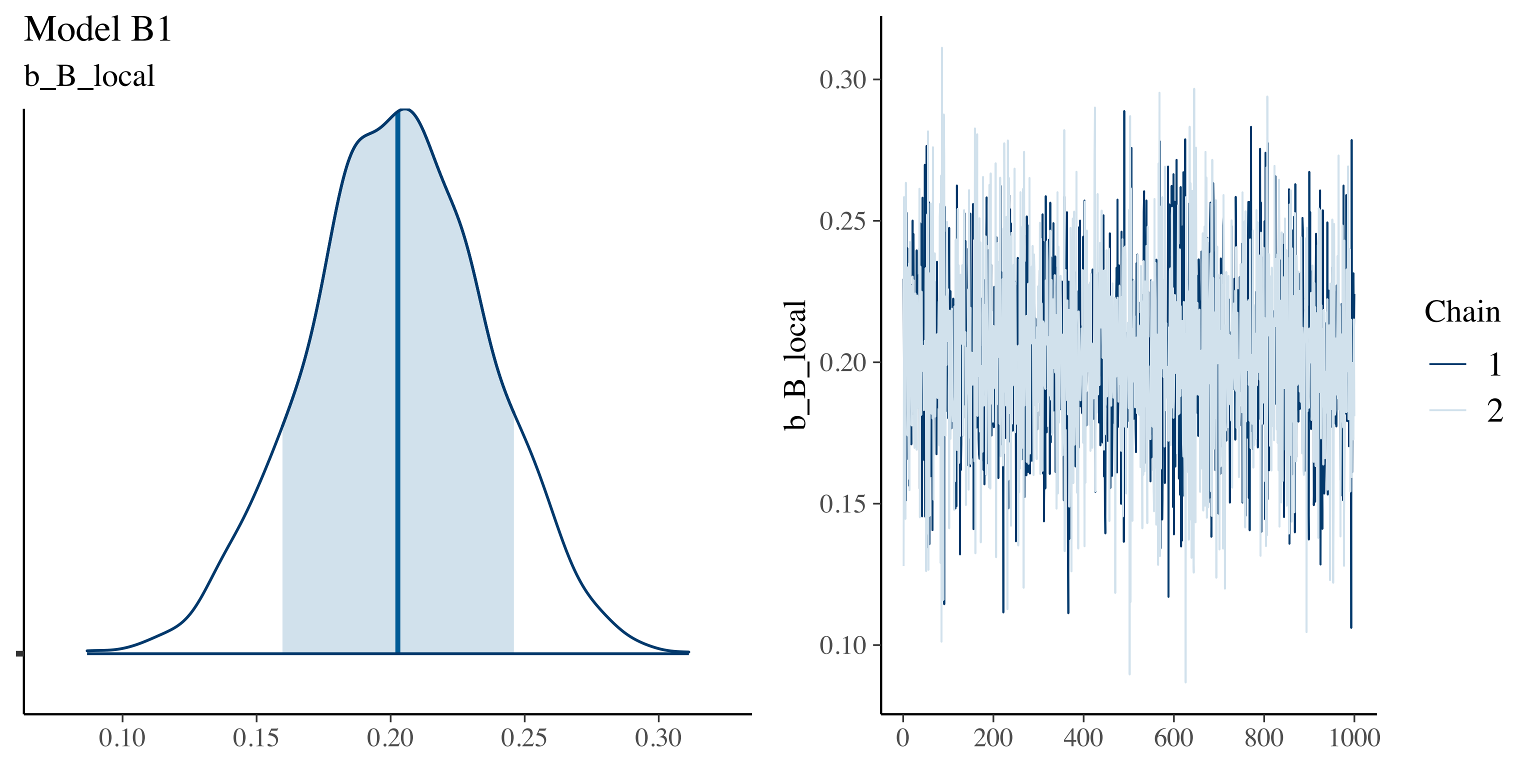
Model B1
b_B_local
0.10
0.15
0.20
0.25
0.30
b_B_local
0
200
400
600
800
1000
Chain
1
2

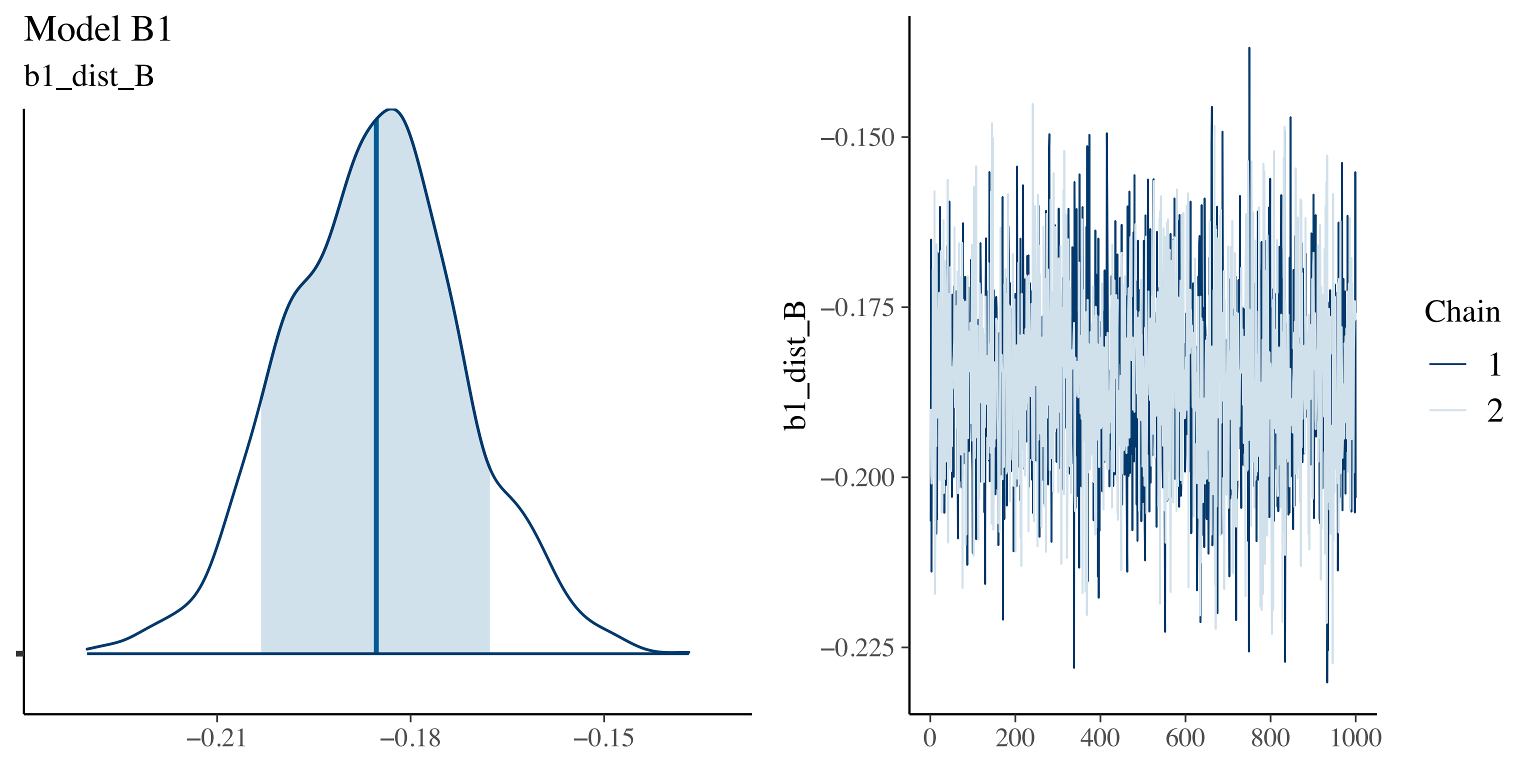
Model B1
b1_dist_B
−0.21
−0.18
−0.15
b1_dist_B
−0.150
−0.175
−0.200
−0.225
0
200
400
600
800
1000
Chain
1
2

Model B2

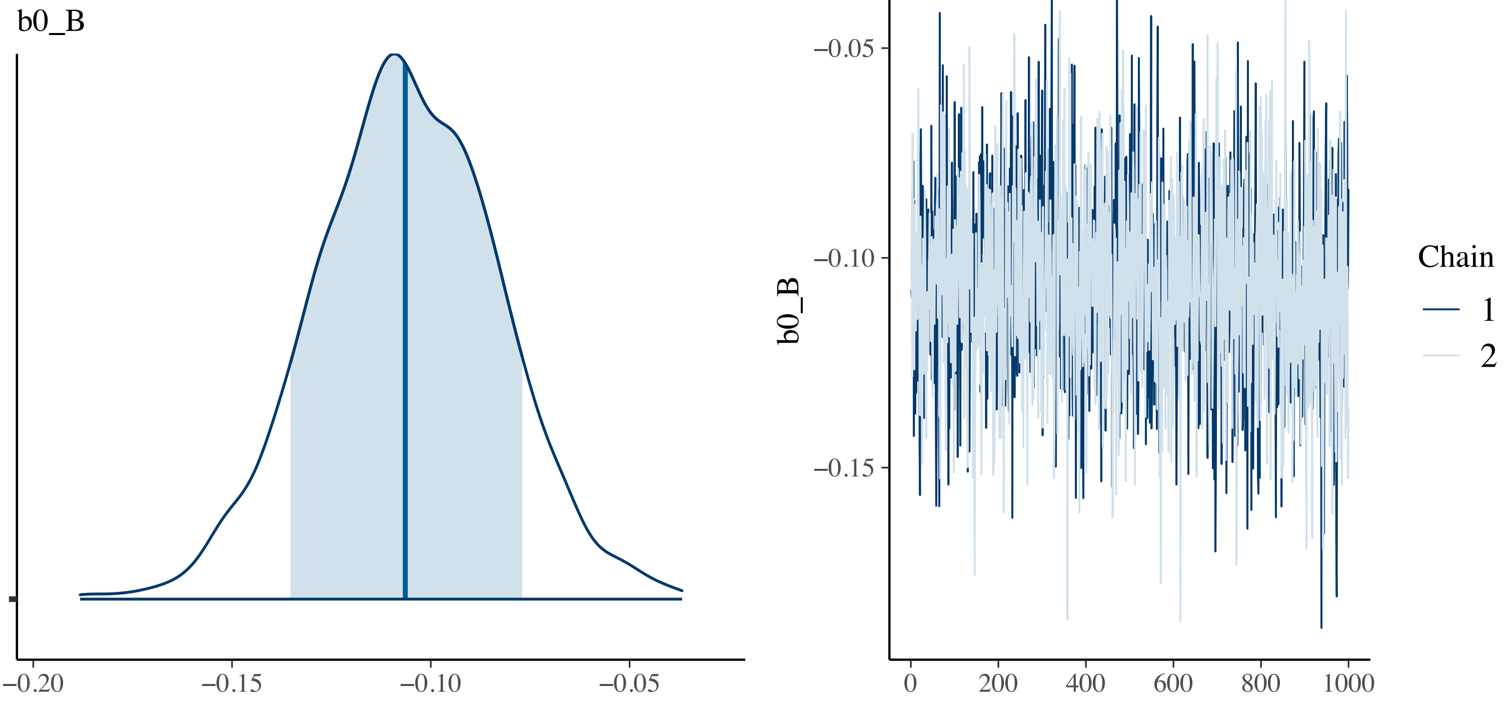
b0_B
−0.20
−0.15
−0.10
−0.05
−0.05
−0.10
−0.15
b0_B
0
200
400
600
800
1000
Chain
1
2

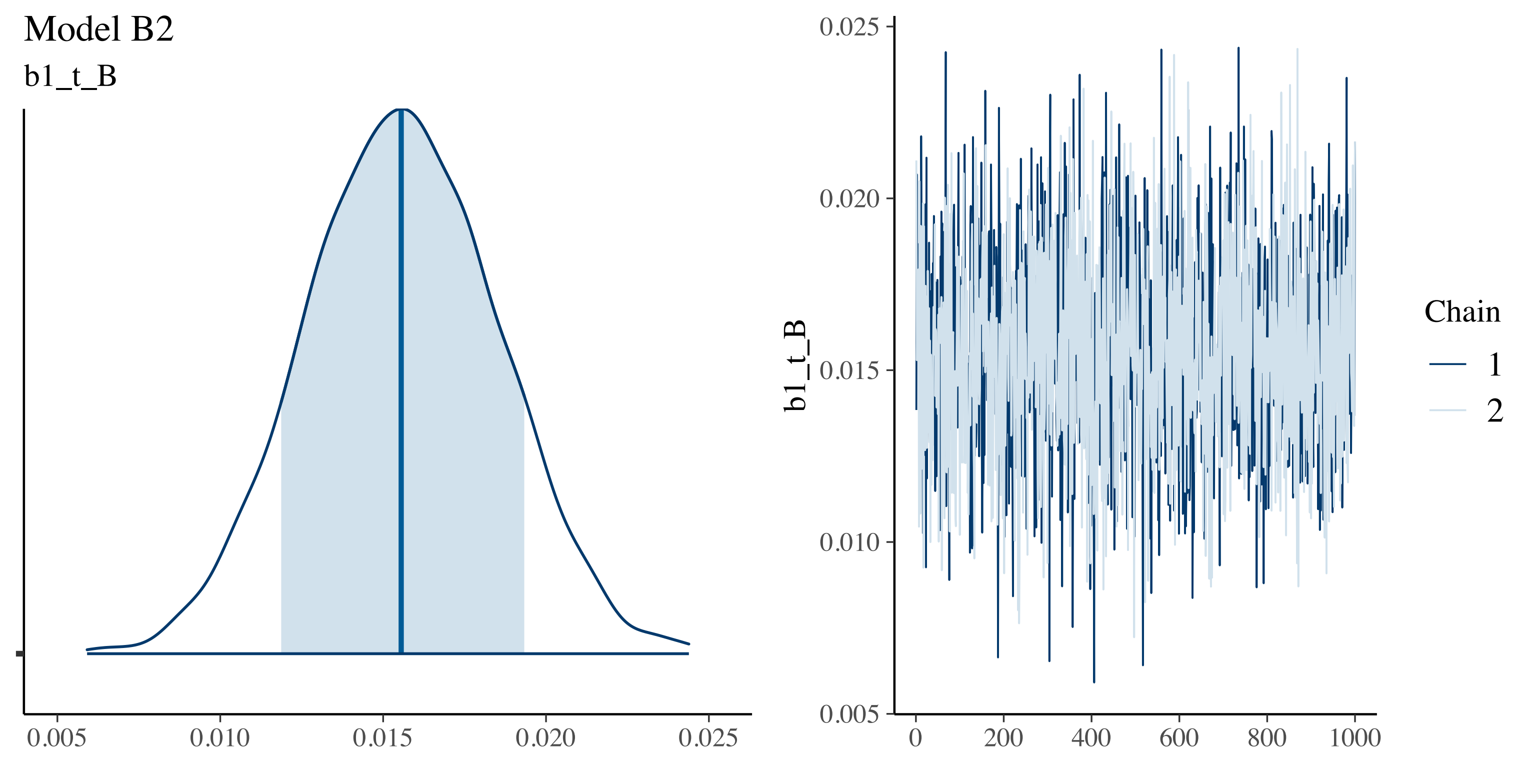

Model B2
b1_t_B
0.005
0.010
0.015
0.020
0.025
b1_t_B
0.025
0.020
0.015
0.010
0.005
0
200
400
600
800
1000
Chain
1
2

Model B2

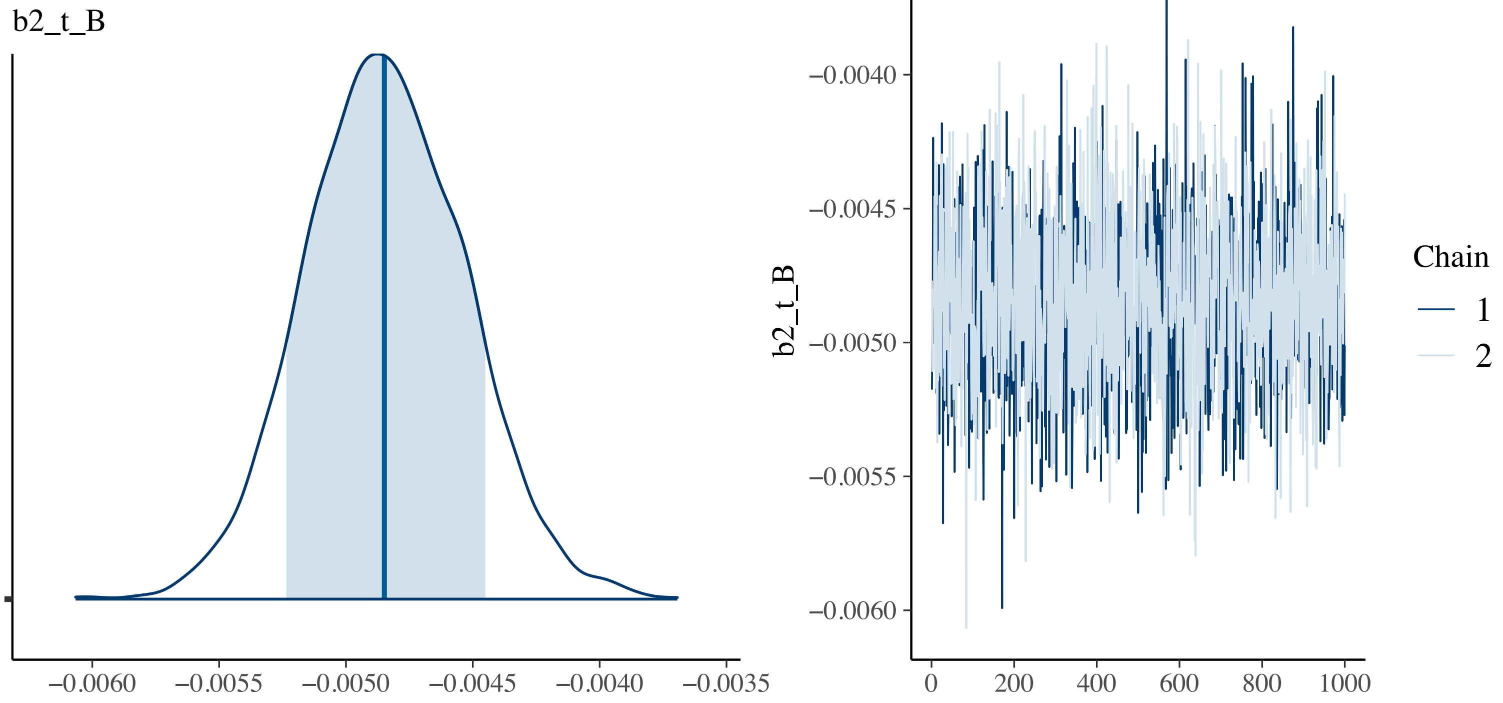
b2_t_B
−0.0060
−0.0055
−0.0050
−0.0045
−0.0040
−0.0035
b2_t_B
−0.0040
−0.0045
−0.0050
−0.0055
−0.0060
0
200
400
600
800
1000
Chain
1
2

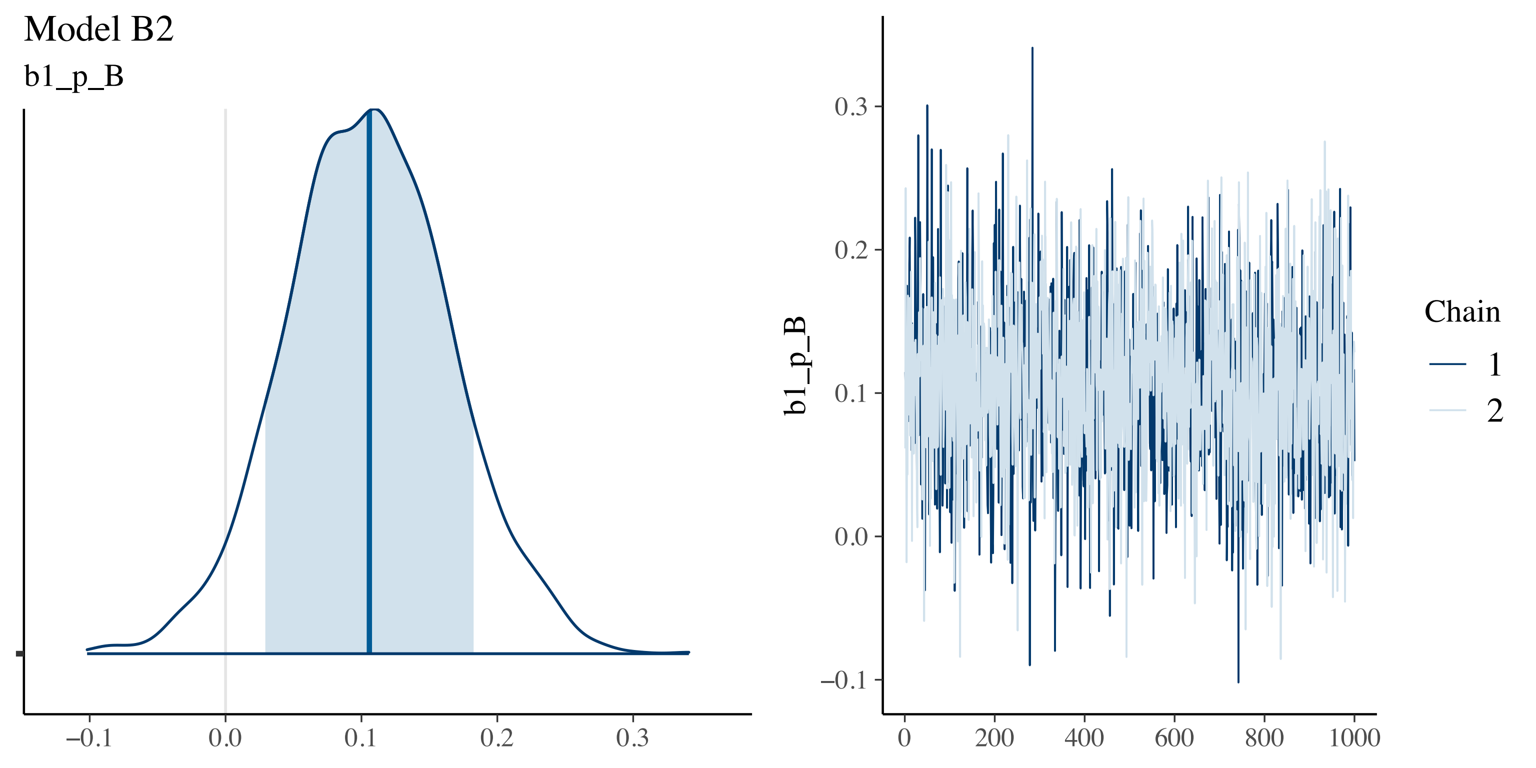
Model B2
b1_p_B
−0.1
0.0
0.1
0.2
0.3
b1_p_B
0.3
0.2
0.1
0.0
−0.1
0
200
400
600
800
1000
Chain
1
2

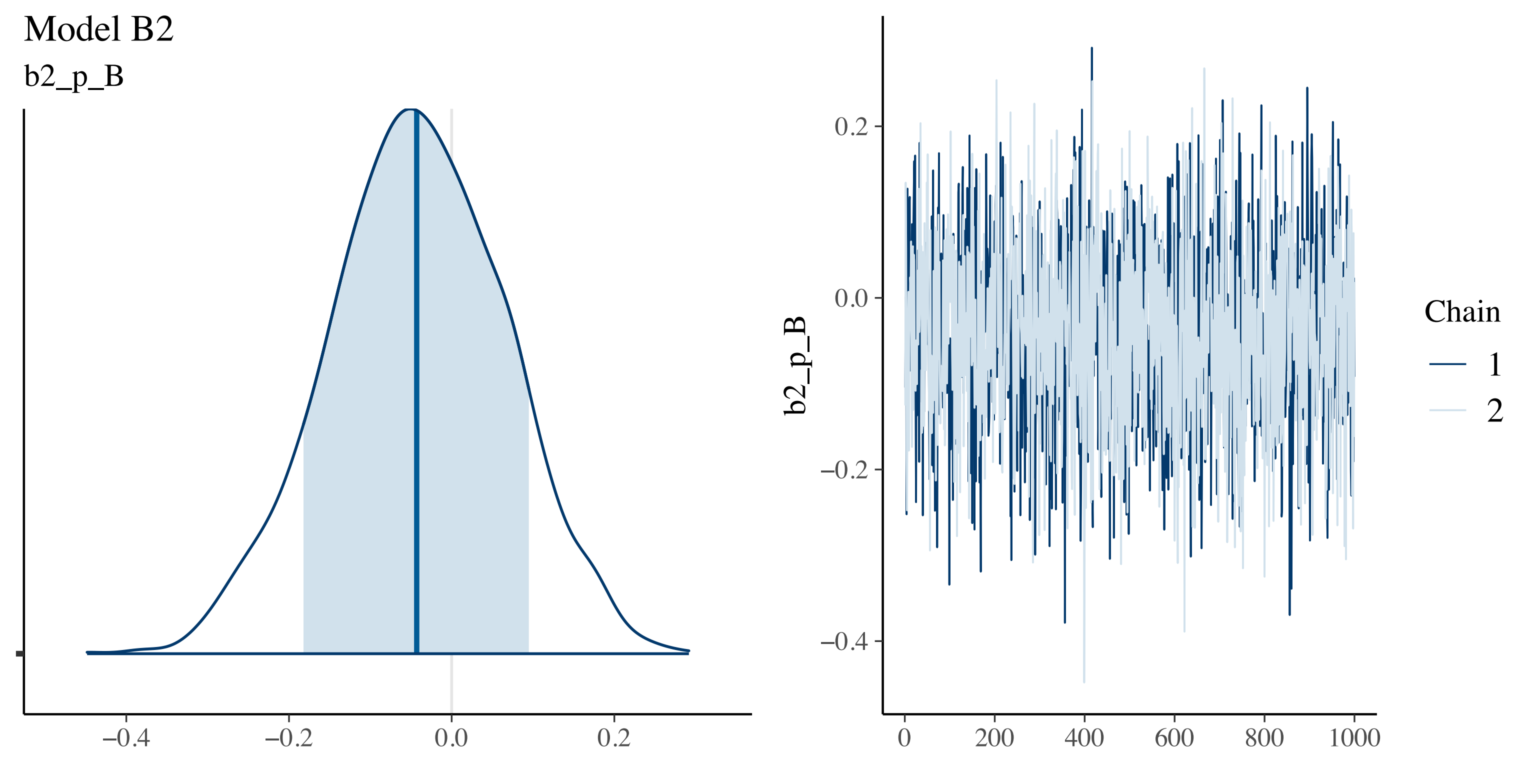
Model B2
b2_p_B
−0.4
−0.2
0.0
0.2
b2_p_B
0.2
0.0
−0.2
−0.4
0
200
400
600
800
1000
Chain
1
2

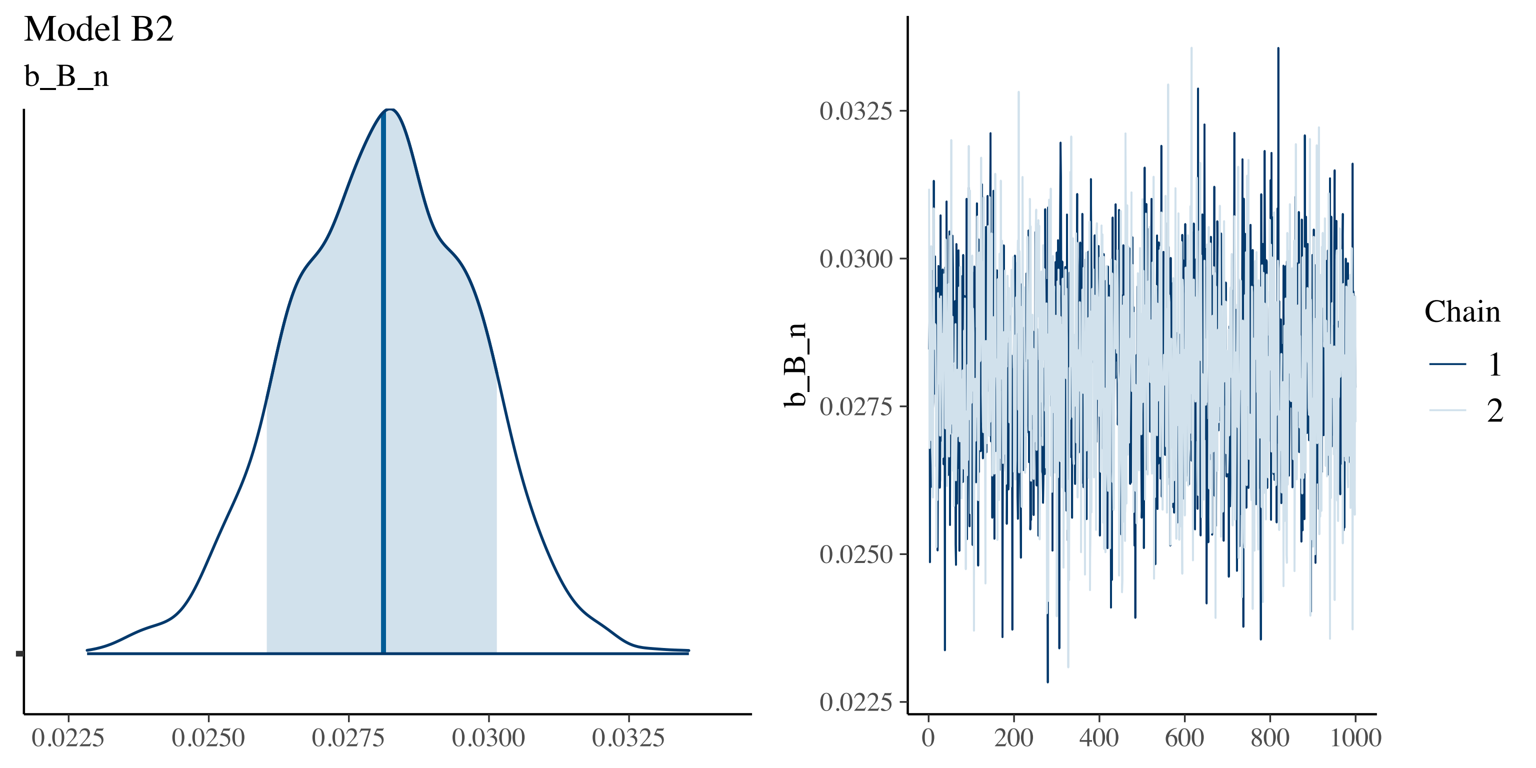
Model B2
b_B_n
0.0225
0.0250
0.0275
0.0300
0.0325
b_B_n
0
200
400
600
800
1000
Chain
1
2

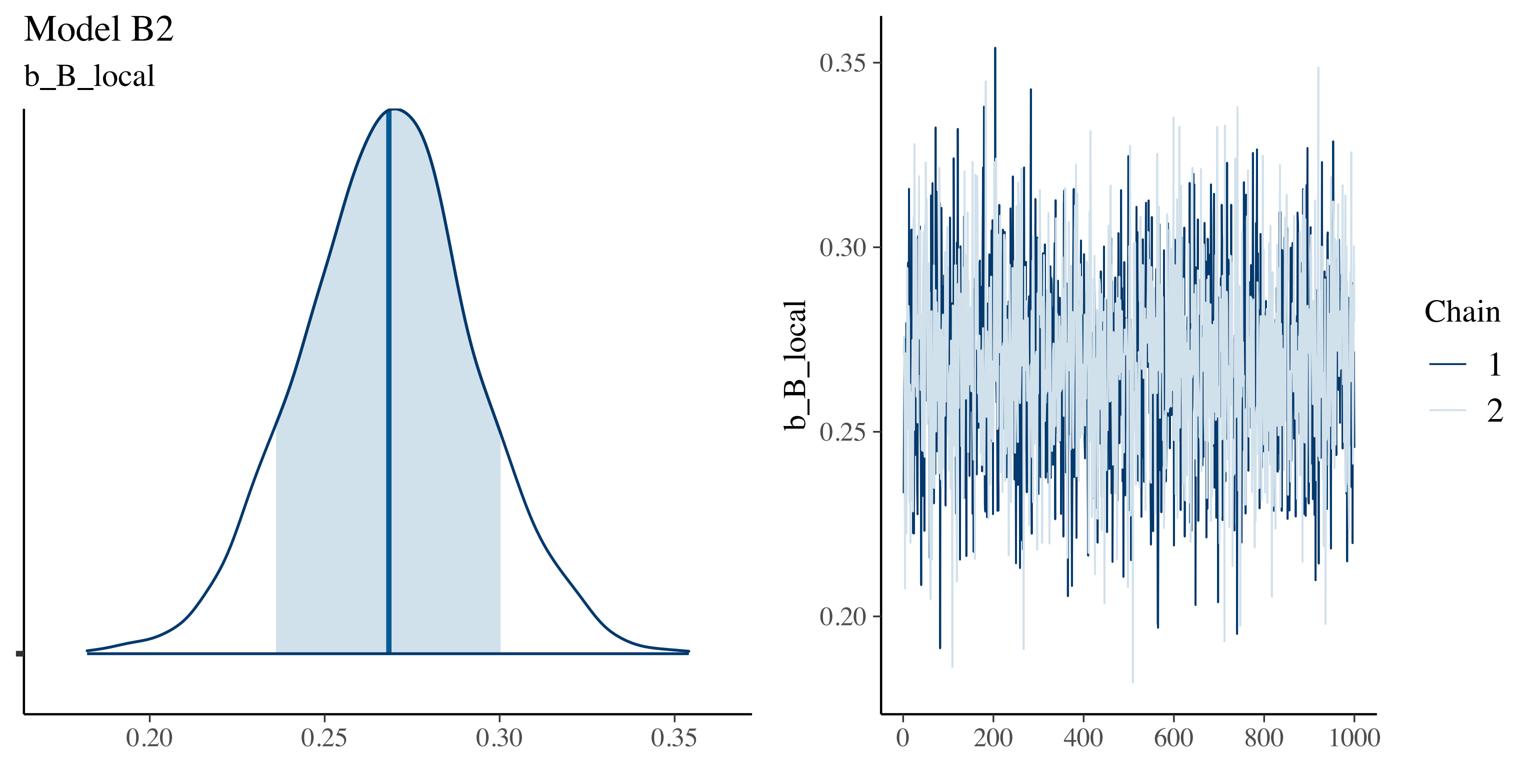
Model B2
b_B_local
0.20
0.25
0.30
0.35
b_B_local
0
200
400
600
800
1000
Chain
1
2

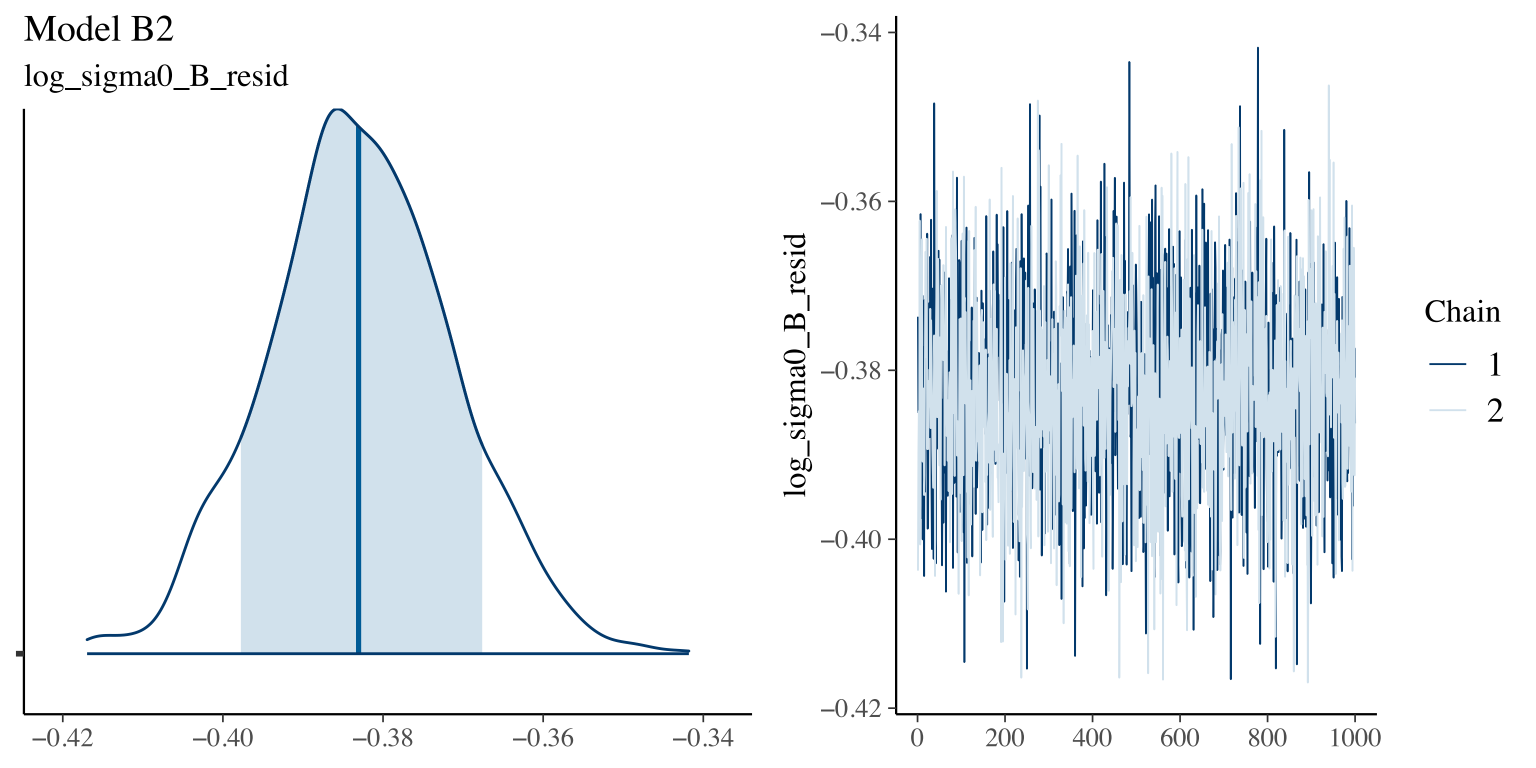

Model B2
log_sigma0_B_resid
−0.42
−0.40
−0.38
−0.36
−0.34
log_sigma0_B_resid
0
200
400
600
800
1000
Chain
1
2

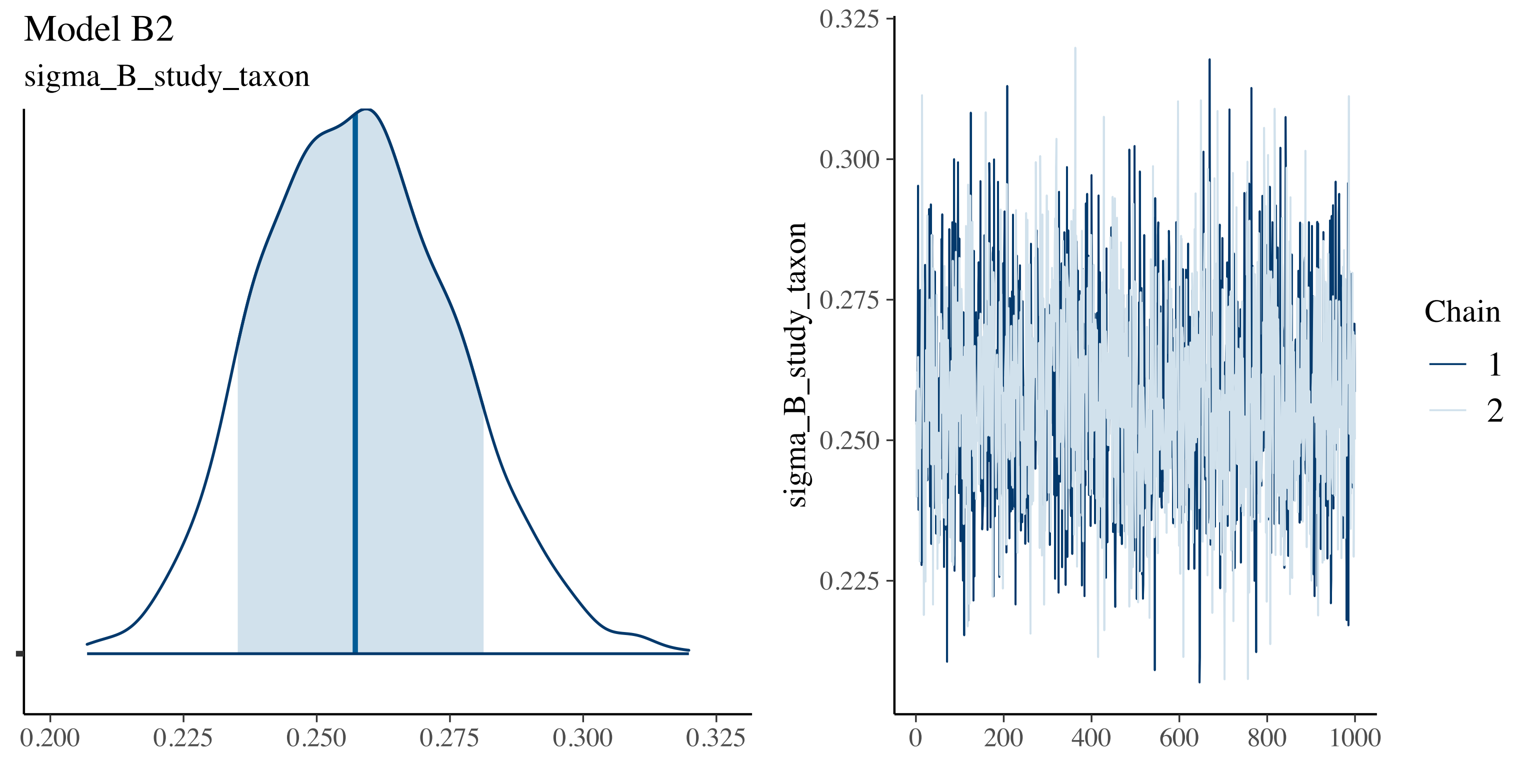
Model B2
sigma_B_study_taxon
0.200
0.225
0.250
0.275
0.300
0.325
sigma_B_study_taxon
0
200
400
600
800
1000
Chain
1
2

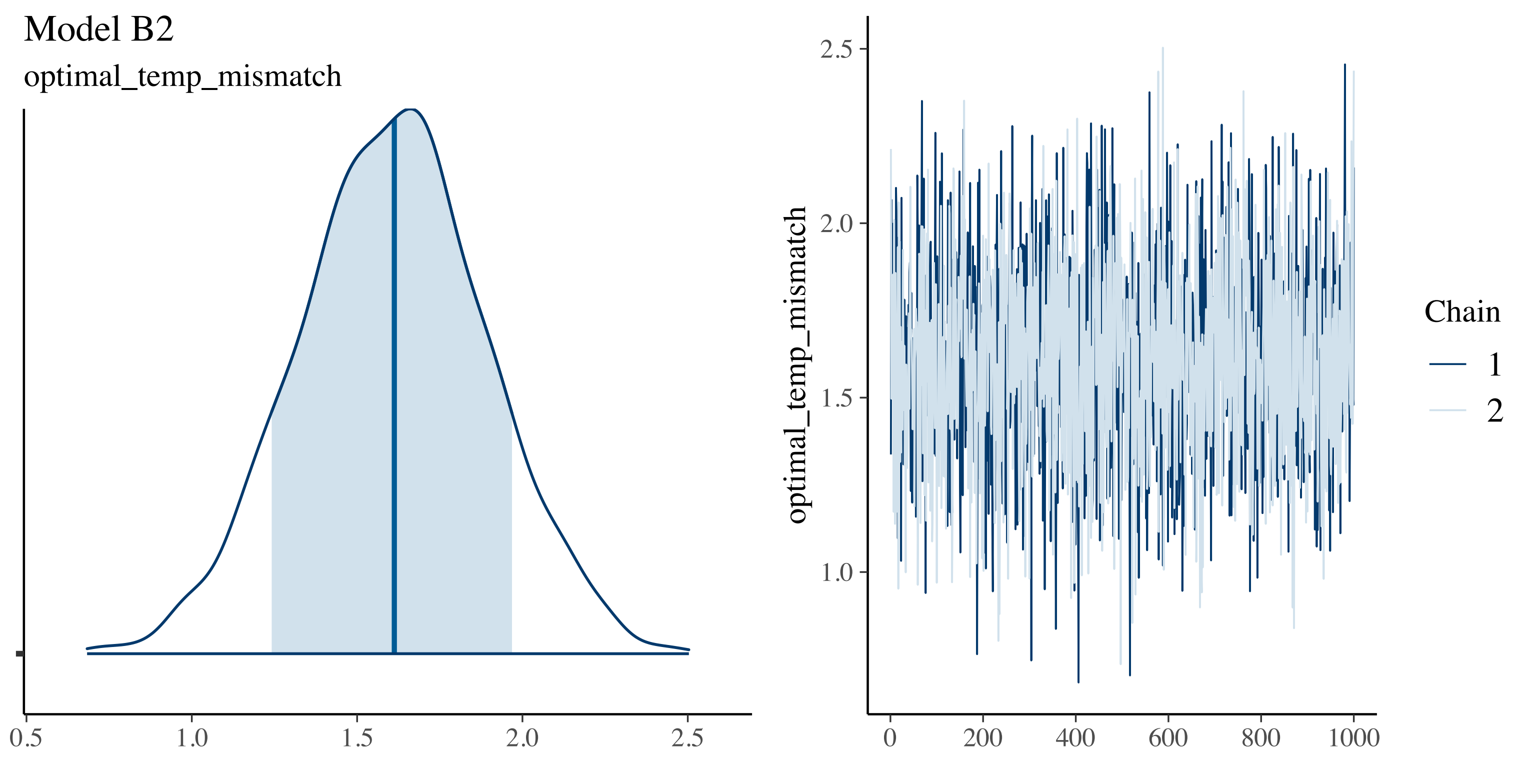

Model B2
optimal_temp_mismatch
0.5
1.0
1.5
2.0
2.5
optimal_temp_mismatch
0
200
400
600
800
1000
Chain
1
2

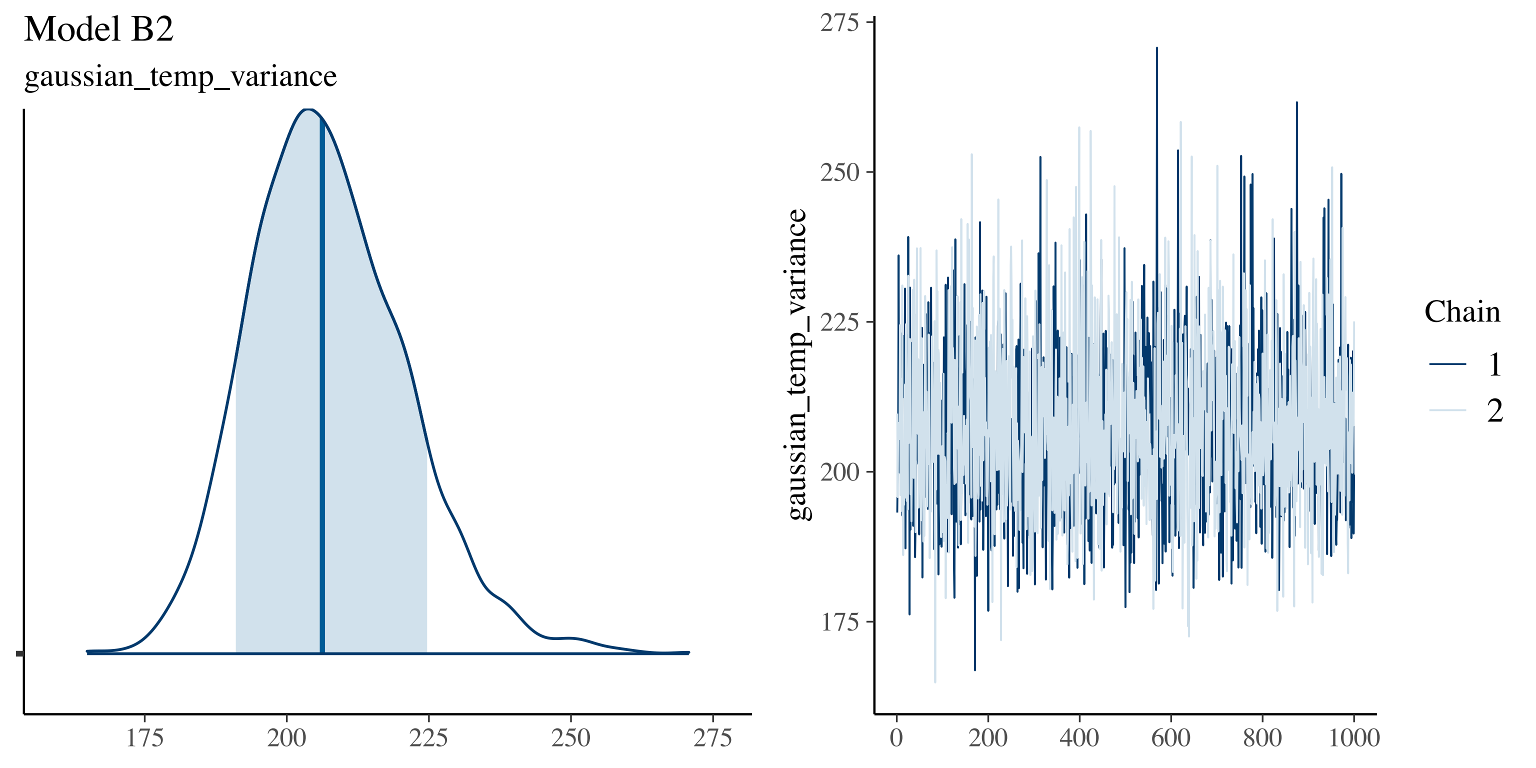

Model B2
gaussian_temp_variance
175
200
225
250
275
gaussian_temp_variance
0
200
400
600
800
1000
Chain
1
2

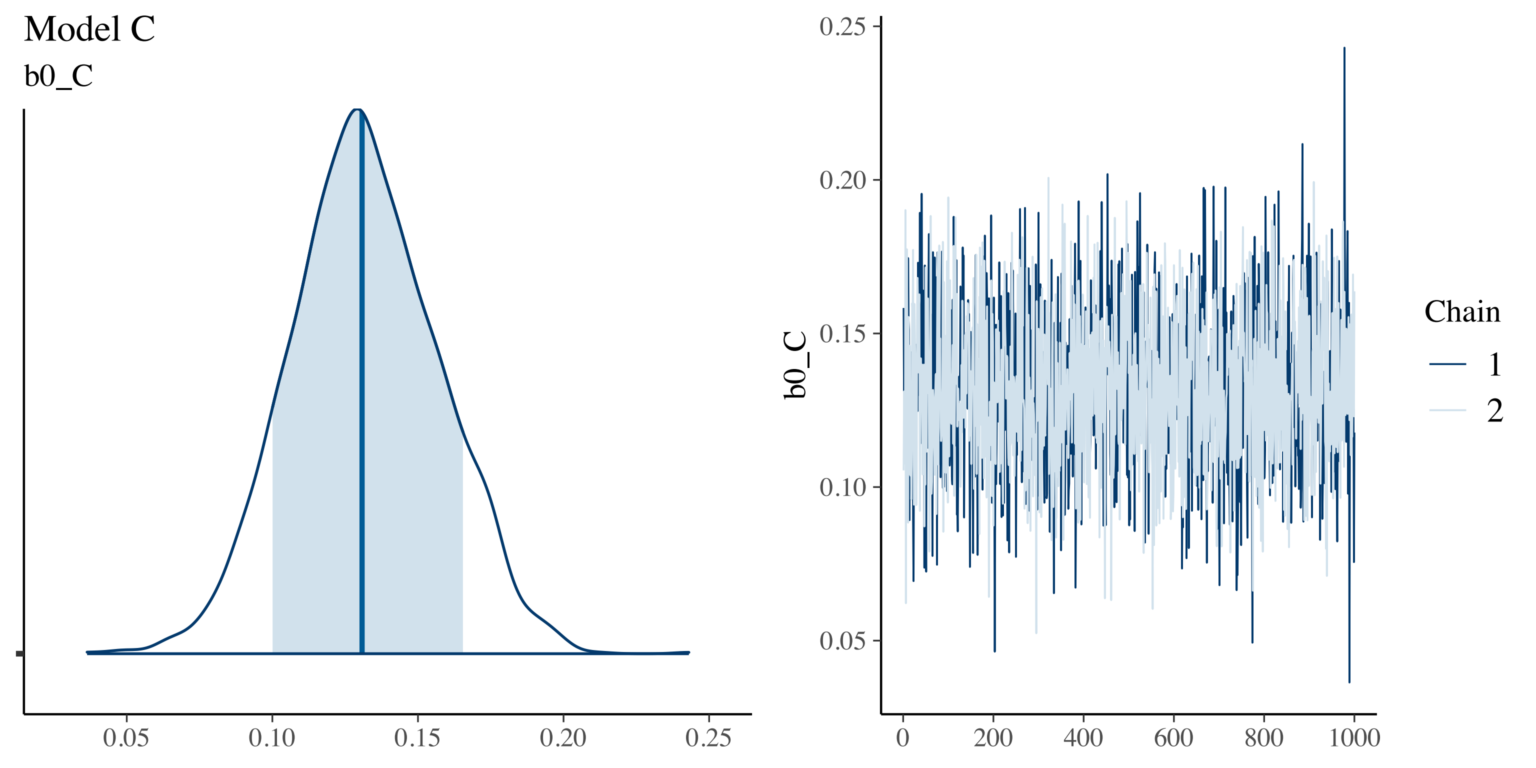

Model C
b0_C
0.05
0.10
0.15
0.20
0.25
b0_C
0
200
400
600
800
1000
Chain
1
2

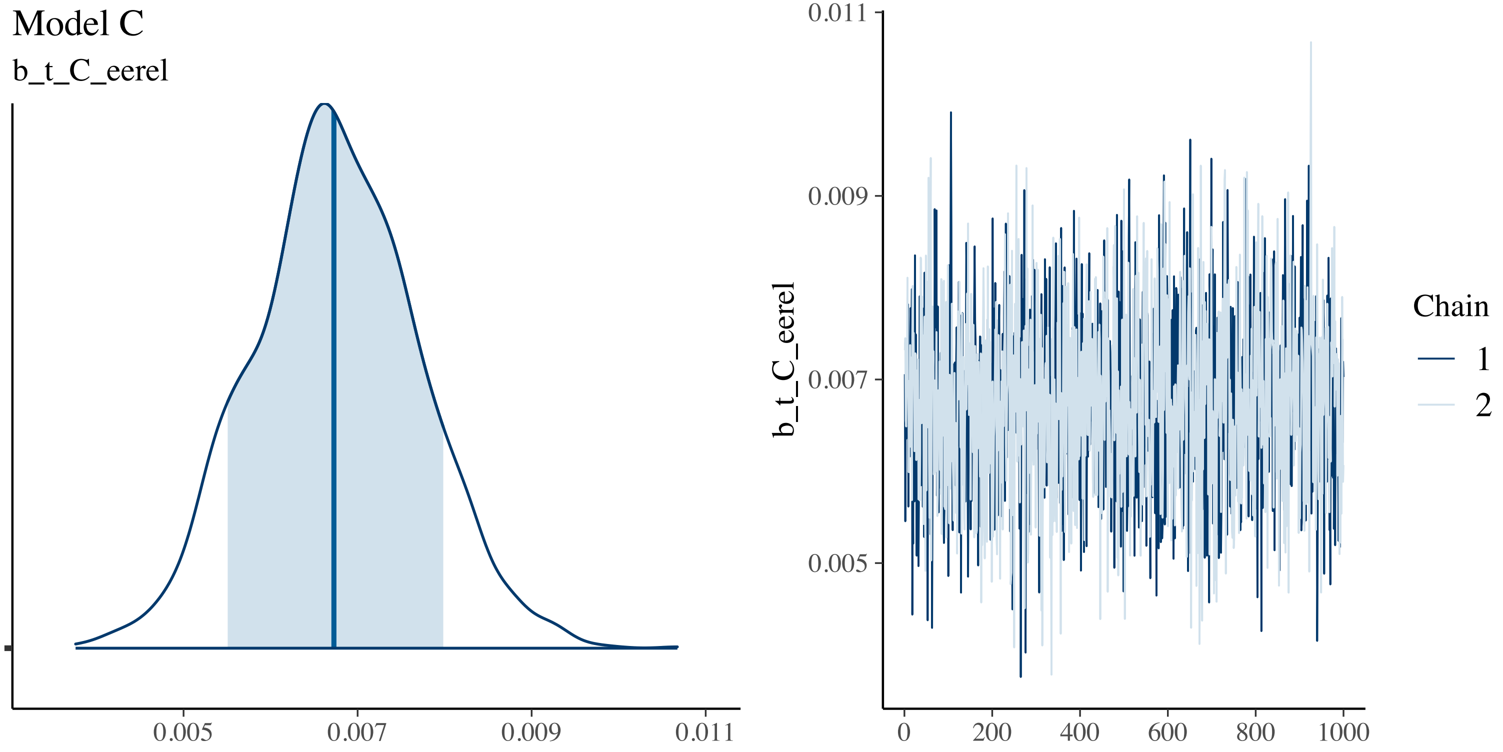
Model C
b_t_C_eerel
0.005
0.007
0.009
0.011
b_t_C_eerel
0.011
0.009
0.007
0.005
0
200
400
600
800
1000
Chain
1
2

Model C

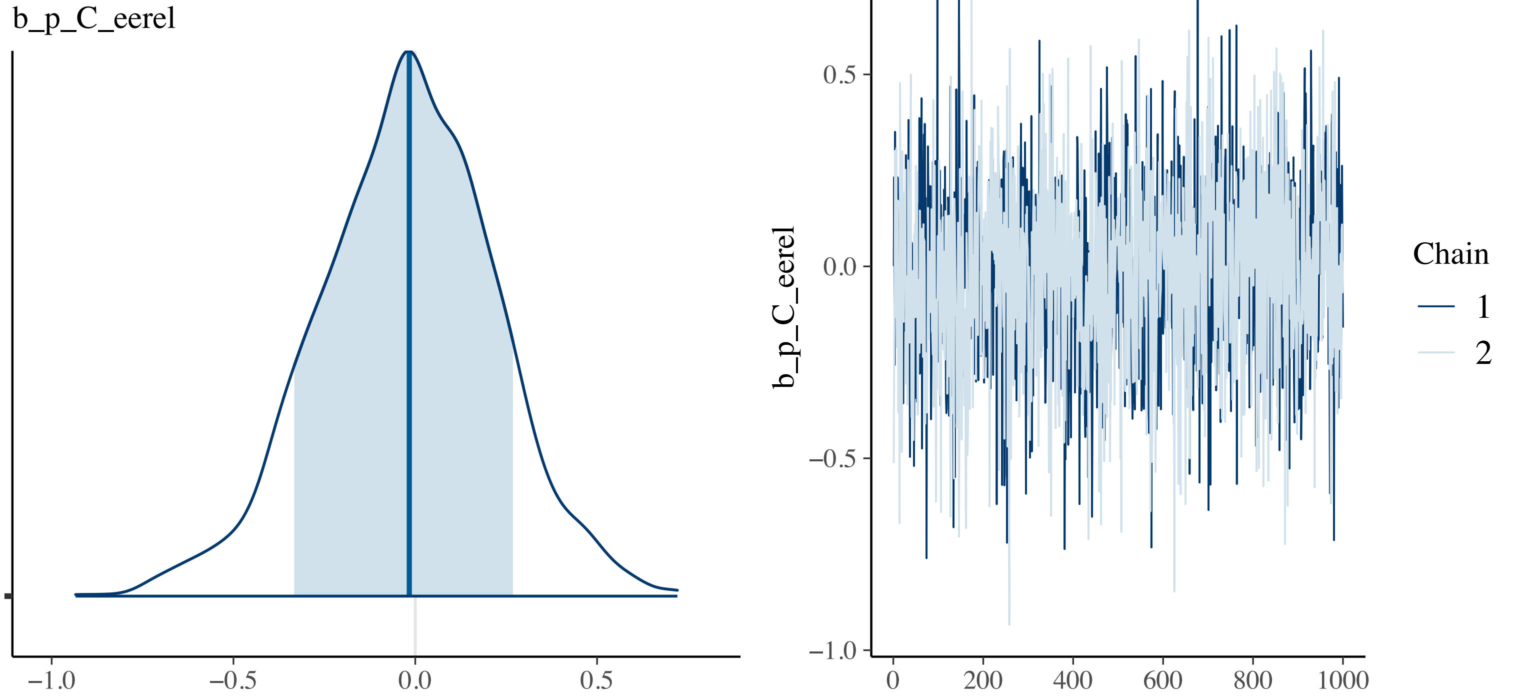

b_p_C_eerel
−1.0
−0.5
0.0
0.5
0.5
0.0
−0.5
−1.0
0
200
400
600
800
1000
b_p_C_eerel
Chain
1
2

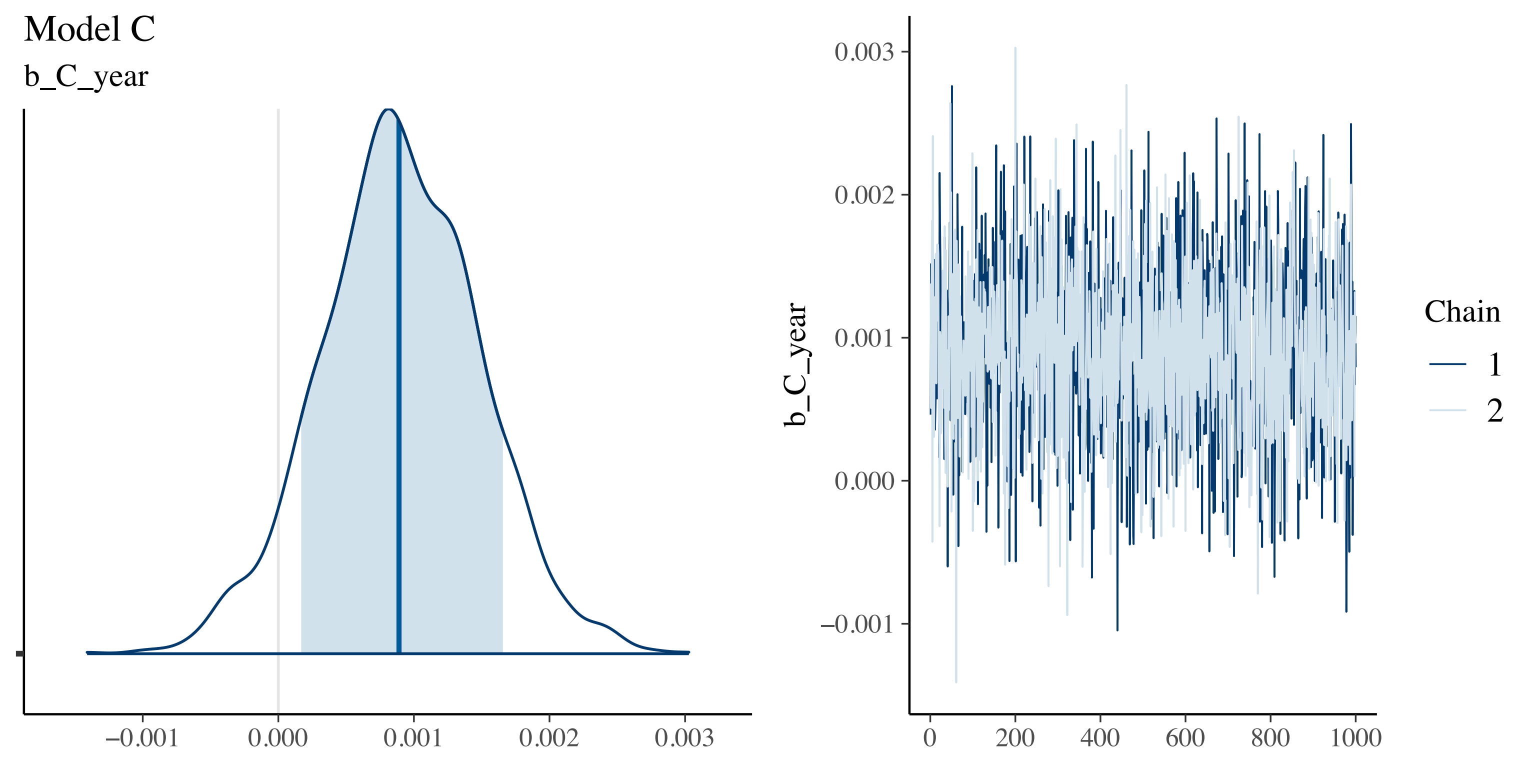
Model C
b_C_year
−0.001
0.000
0.001
0.002
0.003
b_C_year
0
200
400
600
800
1000
Chain
1
2

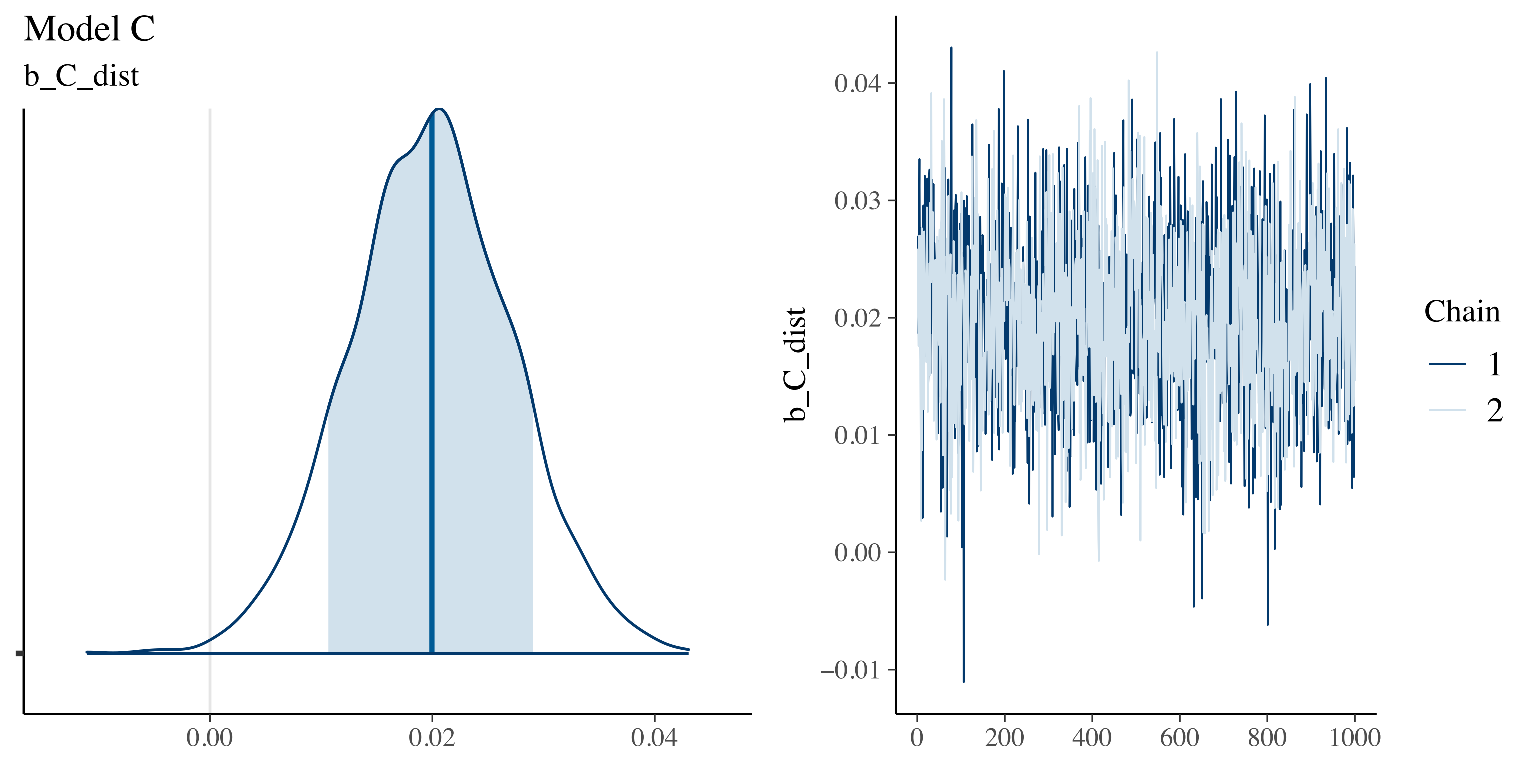

Model C
b_C_dist
0.00
0.02
0.04
b_C_dist
0.04
0.03
0.02
0.01
0.00
−0.01
0
200
400
600
800
1000
Chain
1
2

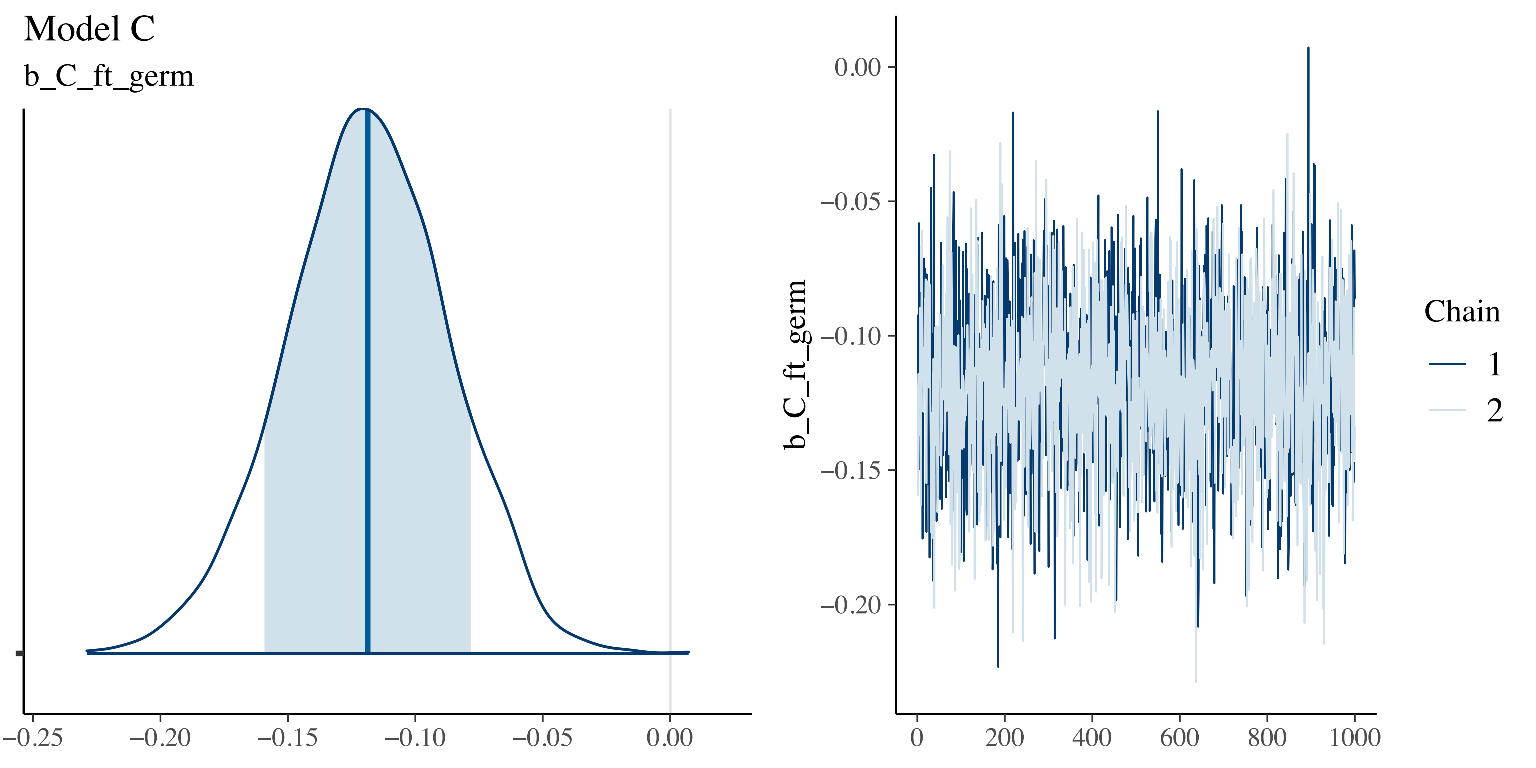

Model C
b_C_ft_germ
−0.25
−0.20
−0.15
−0.10
−0.05
0.00
b_C_ft_germ
0.00
−0.05
−0.10
−0.15
−0.20
0
200
400
600
800
1000
Chain
1
2

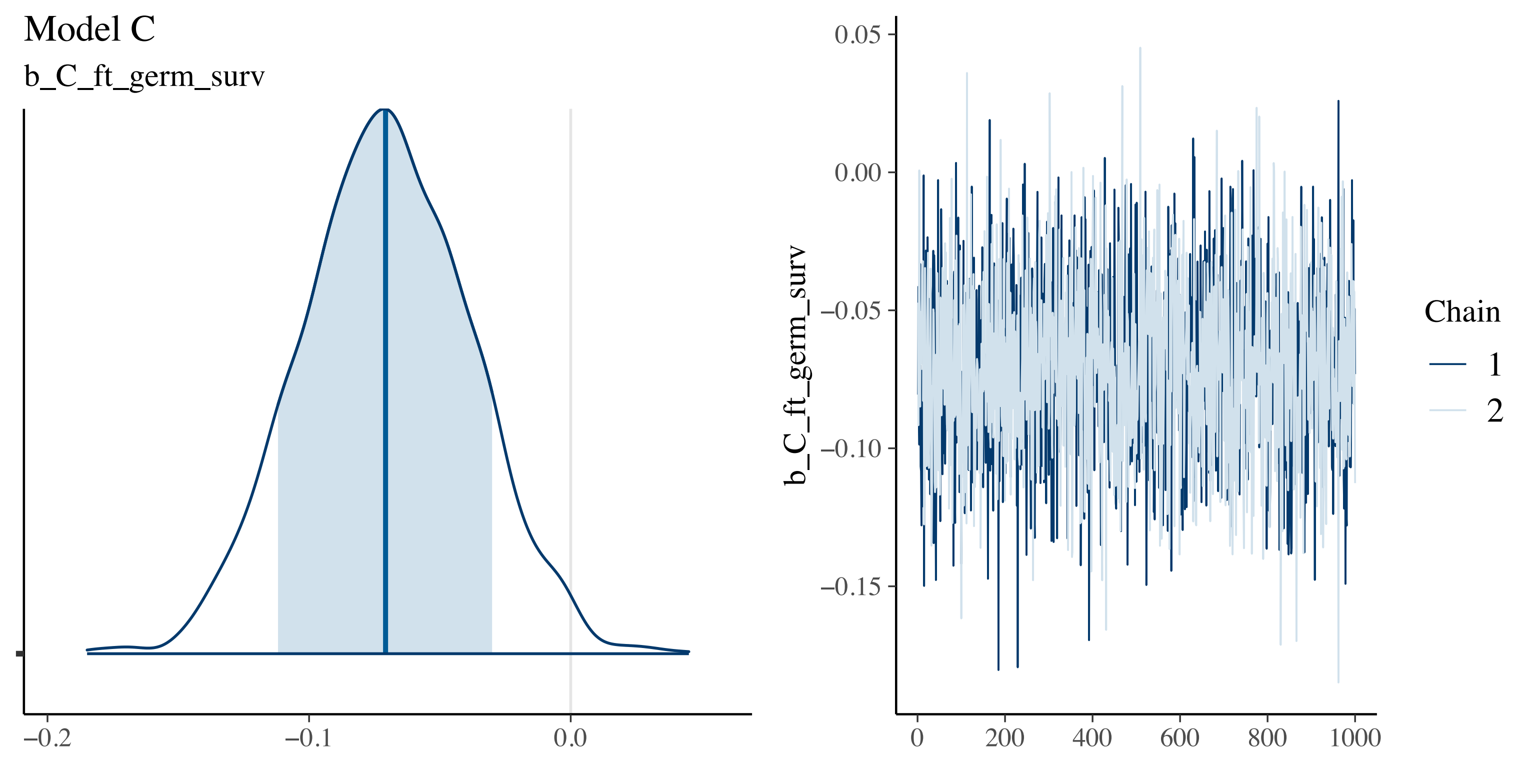

Model C
b_C_ft_germ_surv
−0.2
−0.1
0.0
0.05
0.00
−0.05
−0.10
−0.15
b_C_ft_germ_surv
0
200
400
600
800
1000
Chain
1
2

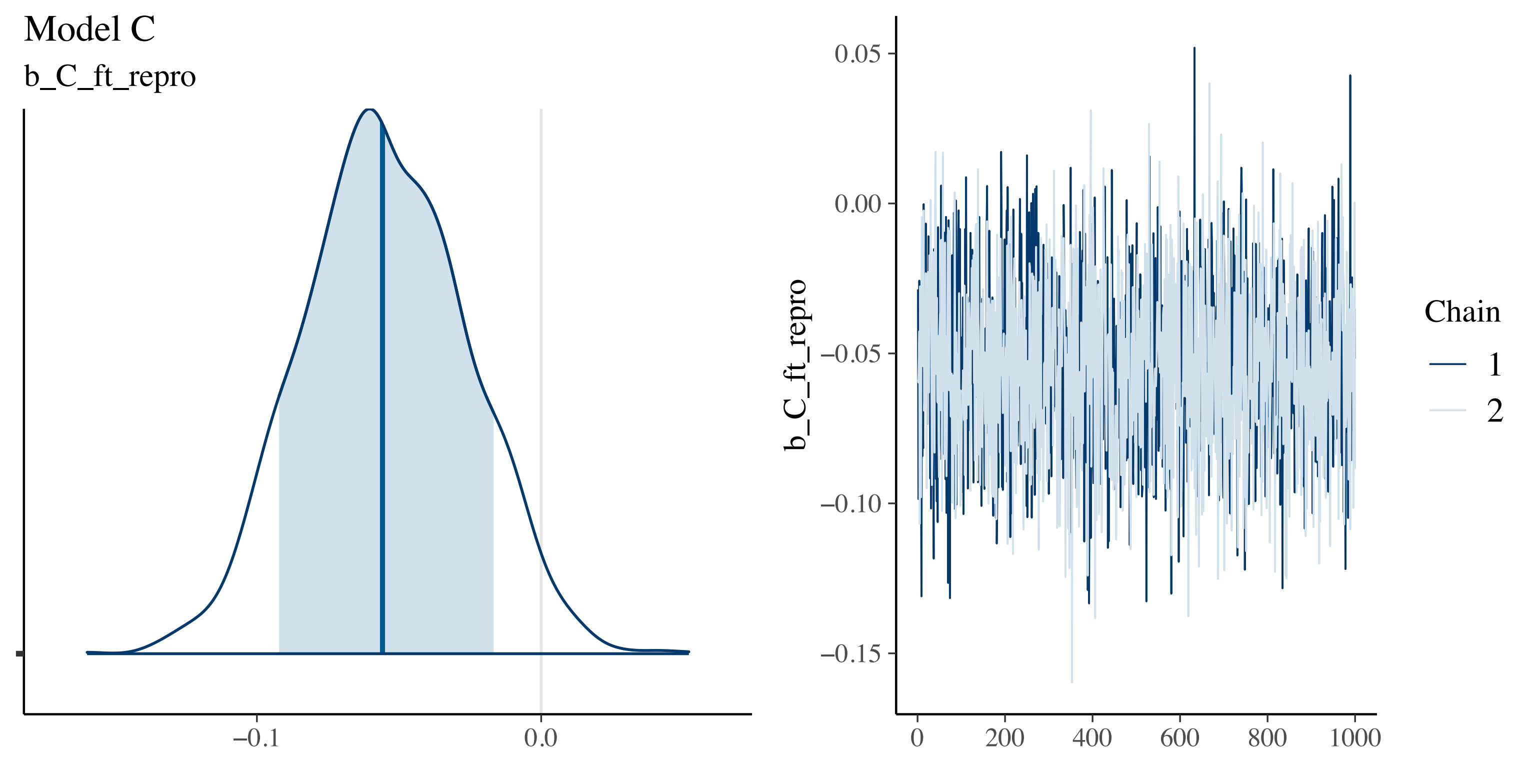

Model C
b_C_ft_repro
−0.1
0.0
0.05
0.00
−0.05
−0.10
−0.15
b_C_ft_repro
0
200
400
600
800
1000
Chain
1
2

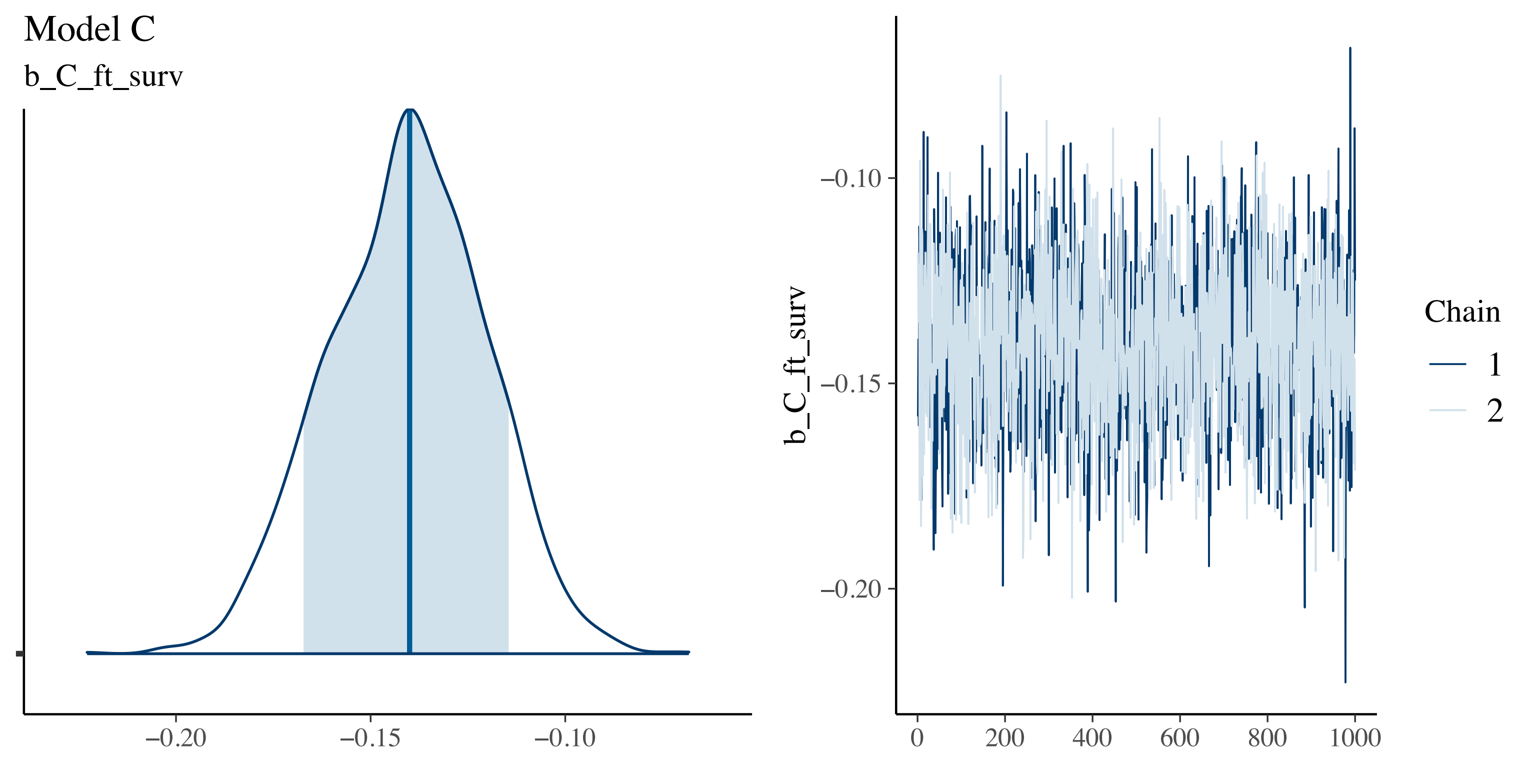

Model C
b_C_ft_surv
−0.20
−0.15
−0.10
b_C_ft_surv
−0.10
−0.15
−0.20
0
200
400
600
800
1000
Chain
1
2

#### Model C

log_sigma0_C_resid

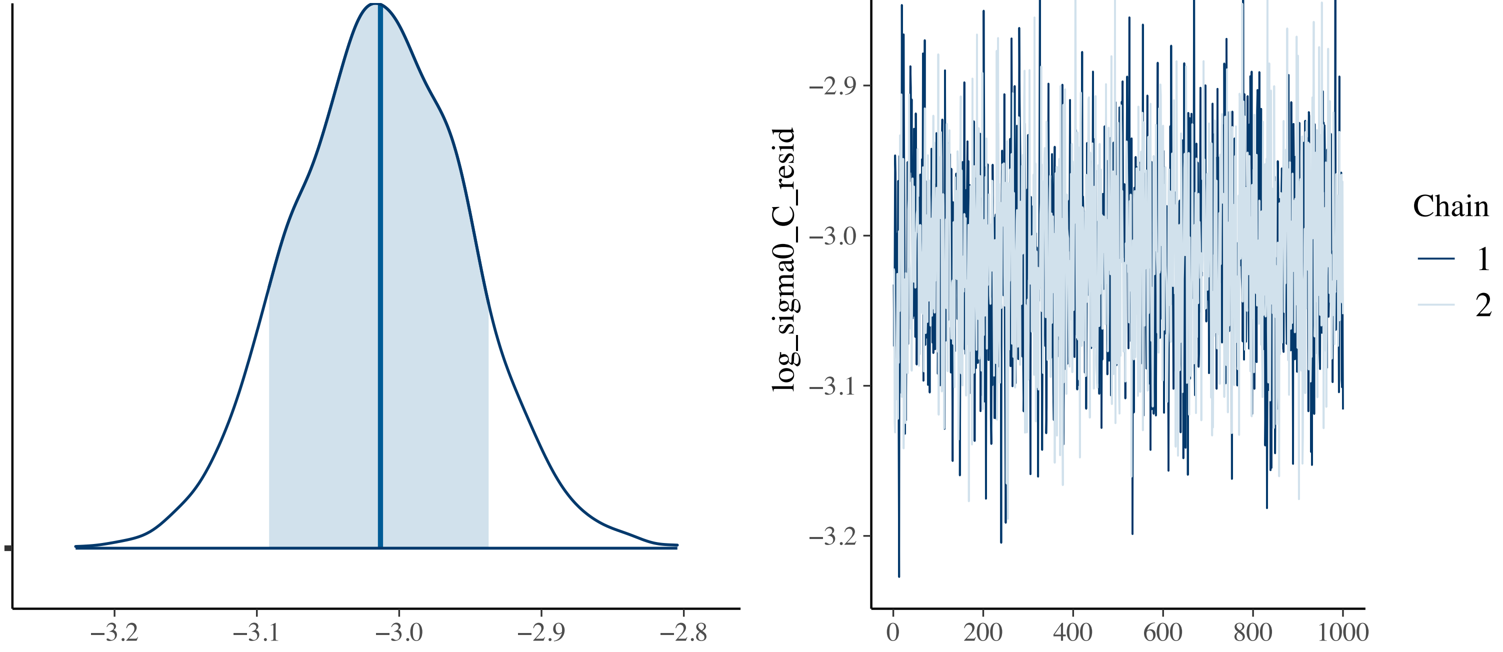

Model C

b_C_resid_year

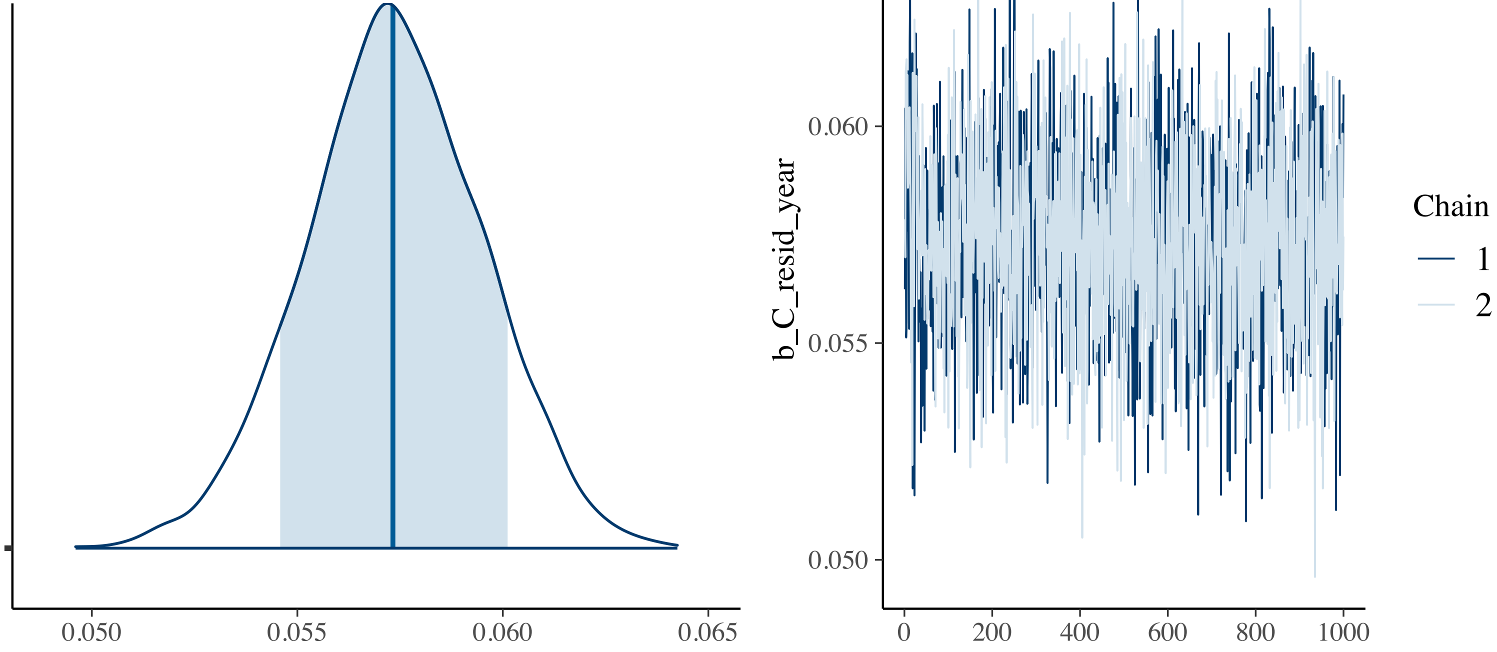

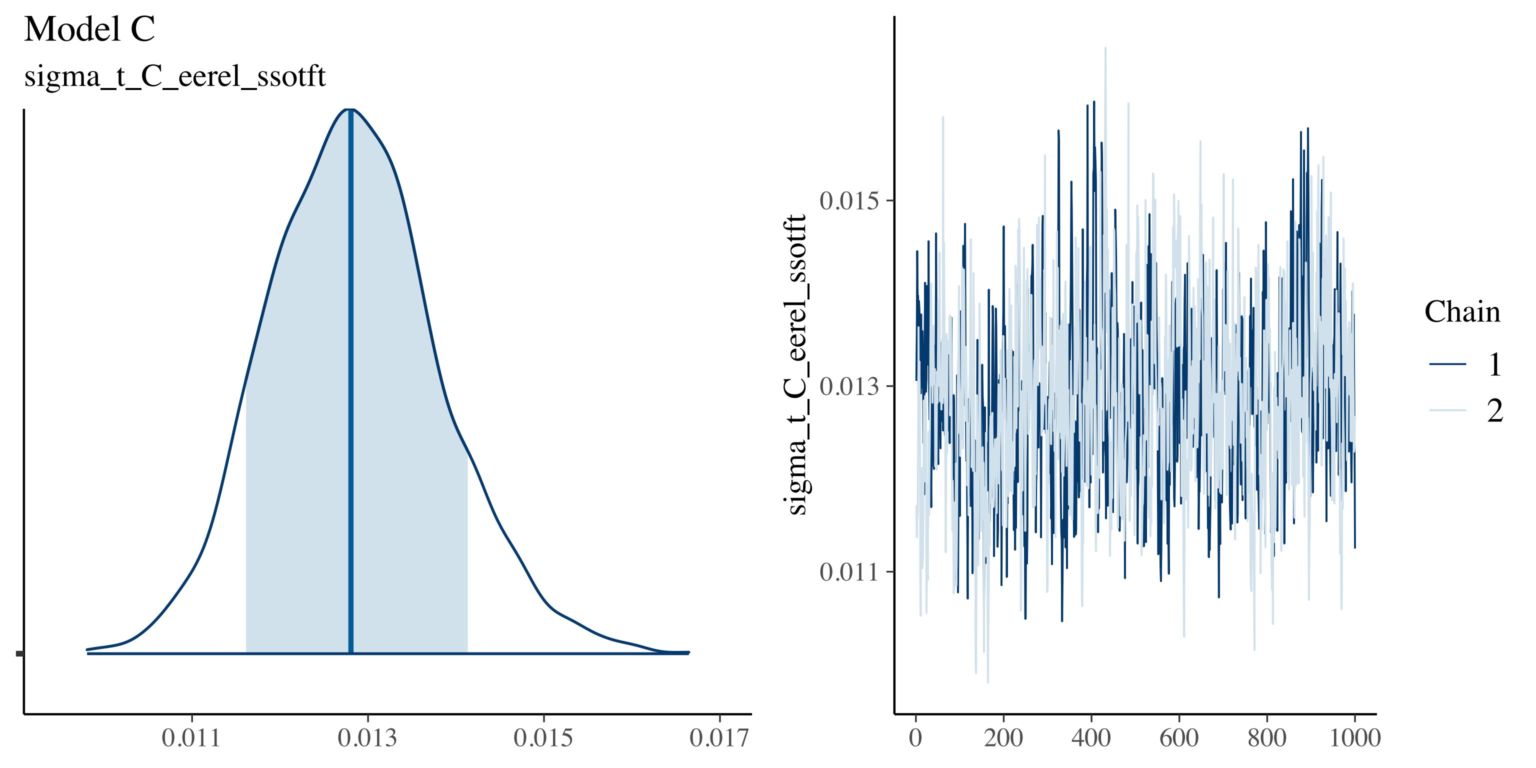

Model C
sigma_t_C_eerel_ssotft
0.011
0.013
0.015
0.017
sigma_t_C_eerel_ssotft
0.015
0.013
0.011
0
200
400
600
800
1000
Chain
1
2

#### Model C

sigma_p_C_eerel_ssotft

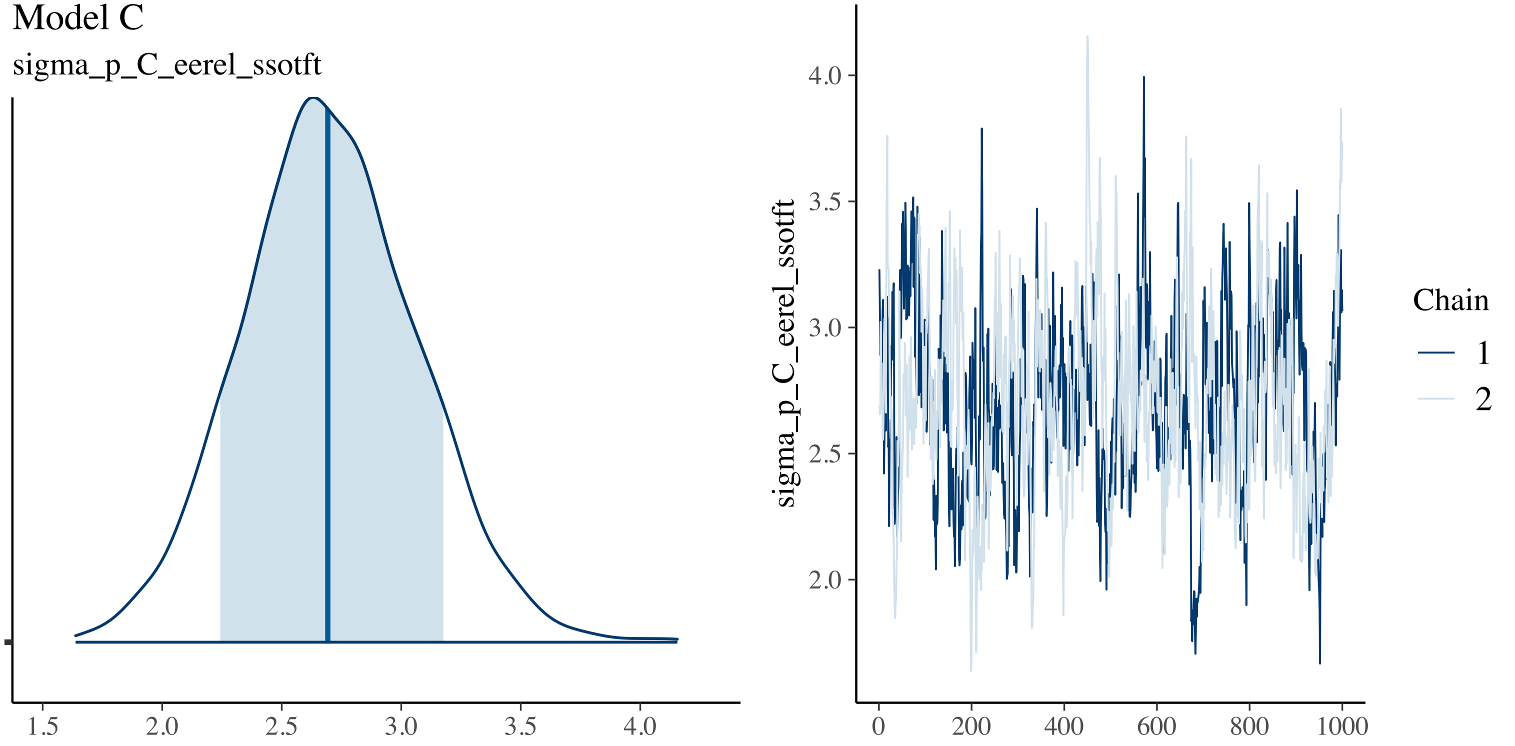

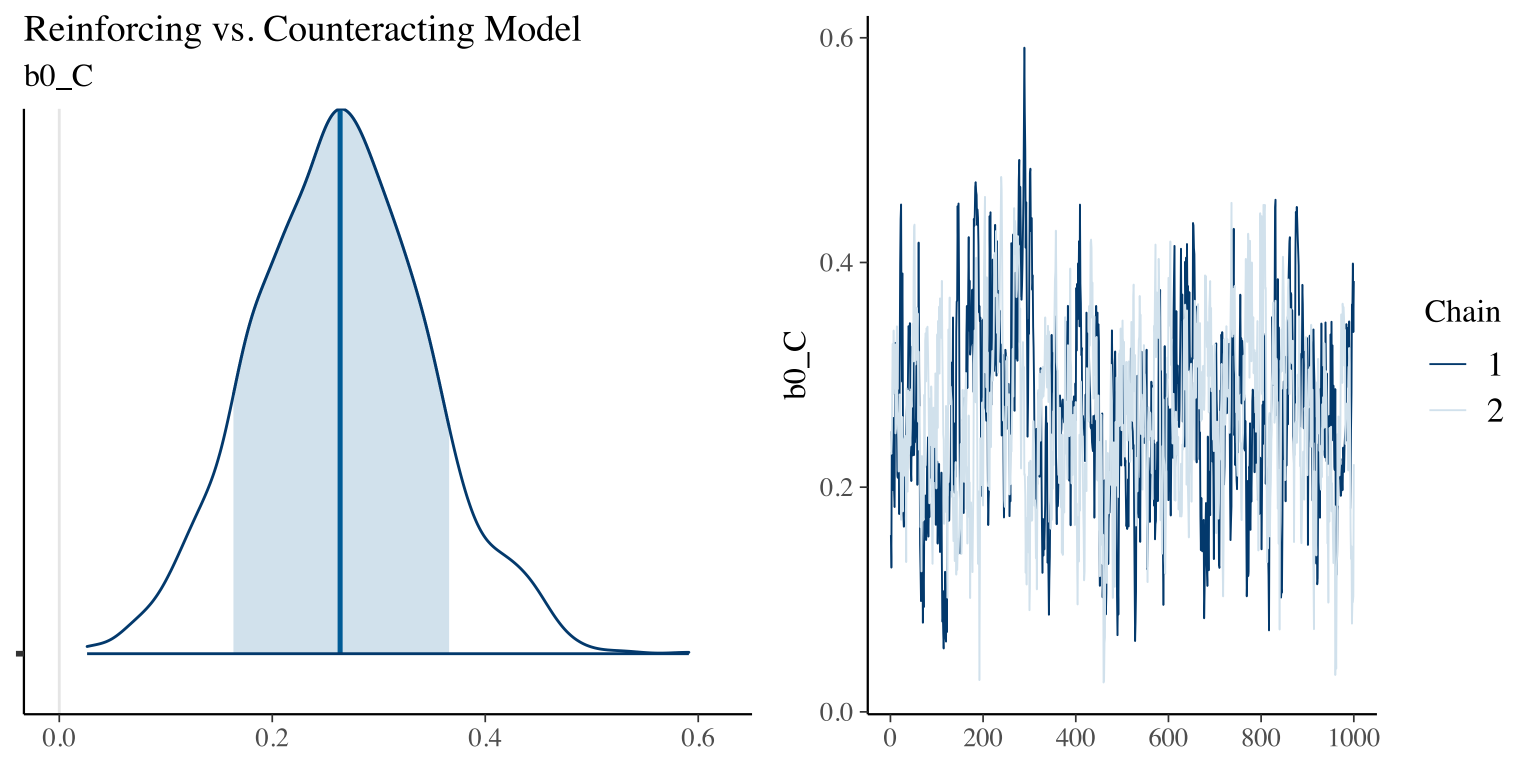
Reinforcing vs. Counteracting Model
b0_C
0.0
0.2
0.4
0.6
b0_C
0.0
0.2
0.4
0.6
0
200
400
600
800
1000
Chain
1
2

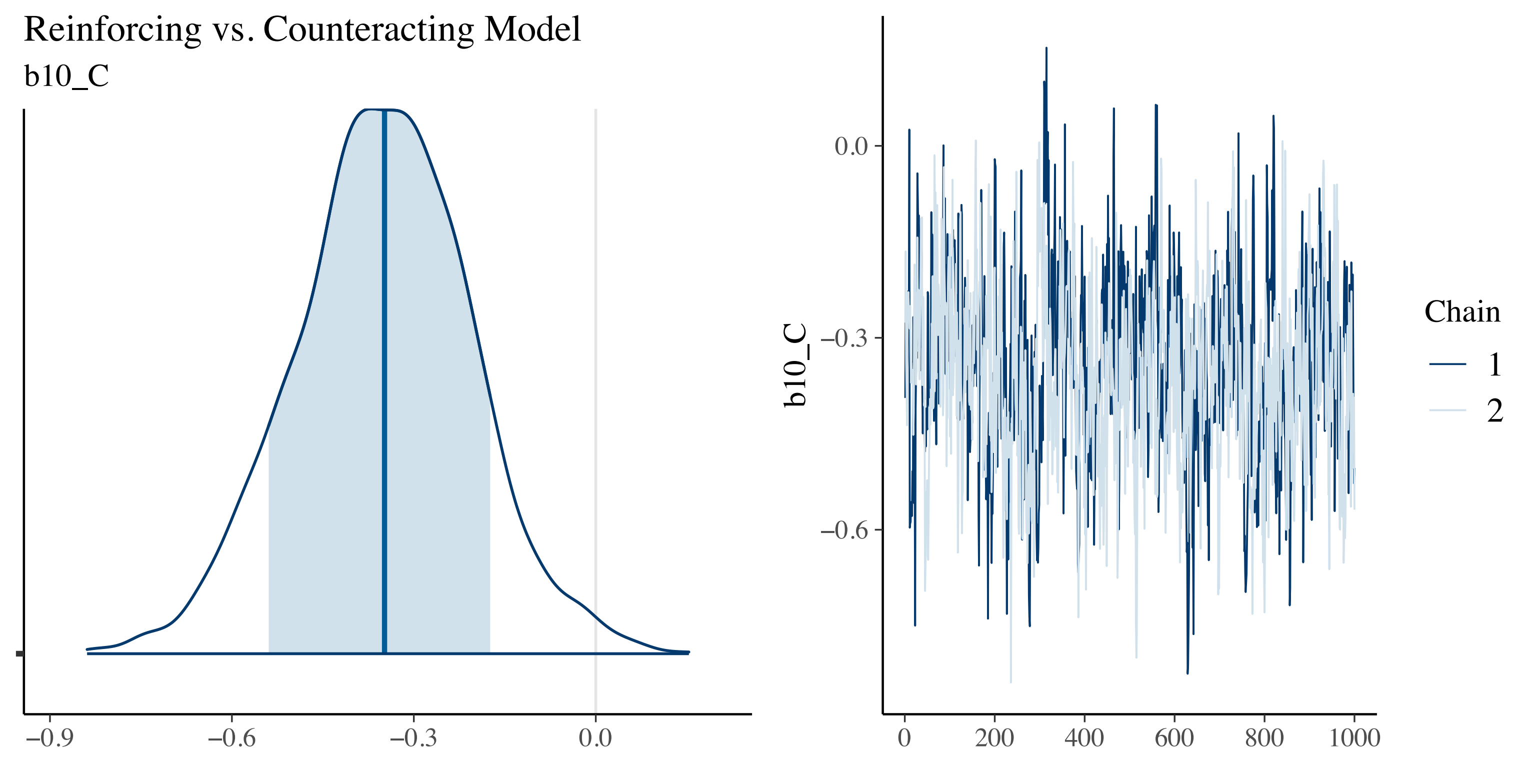
Reinforcing vs. Counteracting Model
b10_C
−0.9
−0.6
−0.3
0.0
b10_C
0.0
−0.3
−0.6
0
200
400
600
800
1000
Chain
1
2

Reinforcing vs. Counteracting Model

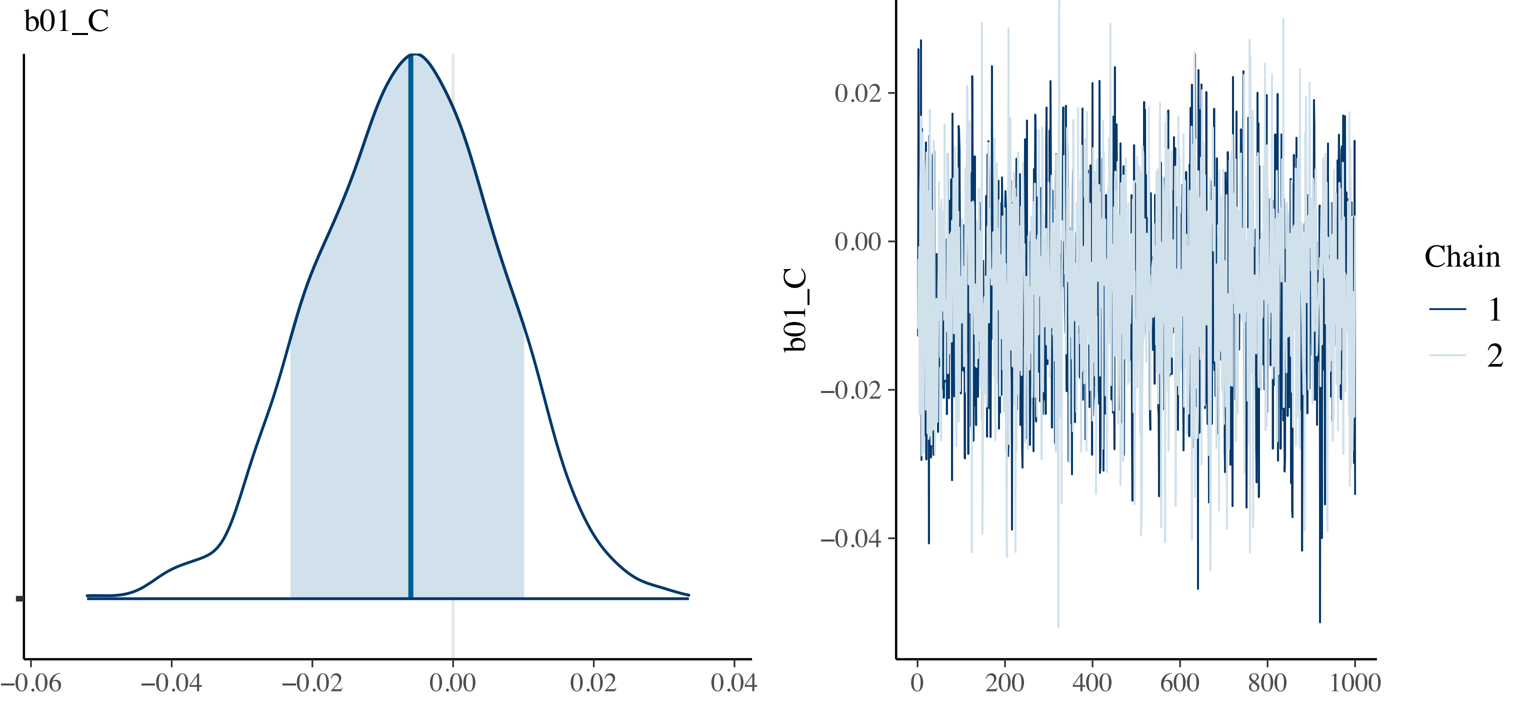

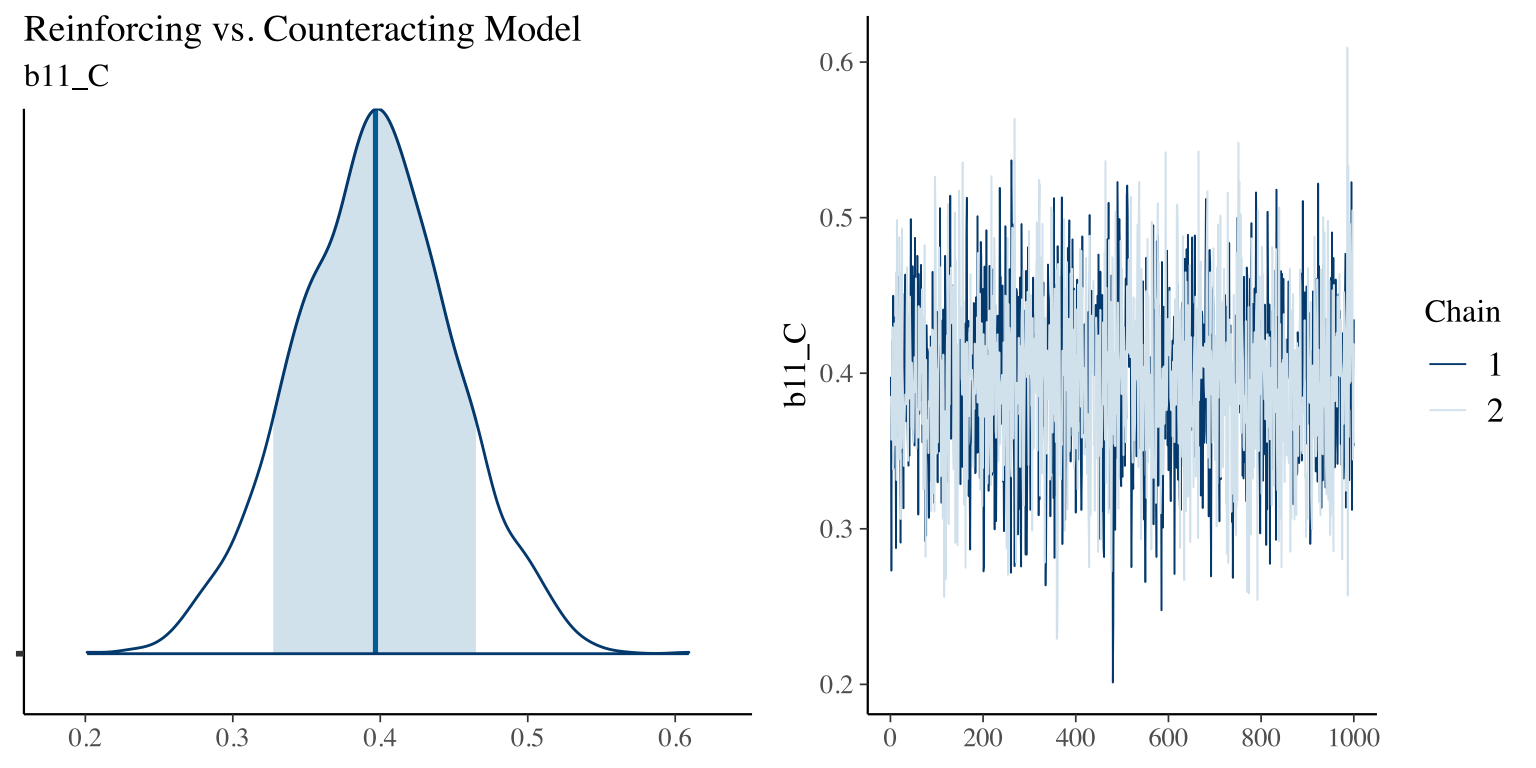

Reinforcing vs. Counteracting Model
b11_C
0.2
0.3
0.4
0.5
0.6
b11_C
0.2
0.3
0.4
0.5
0.6
0
200
400
600
800
1000
Chain
1
2

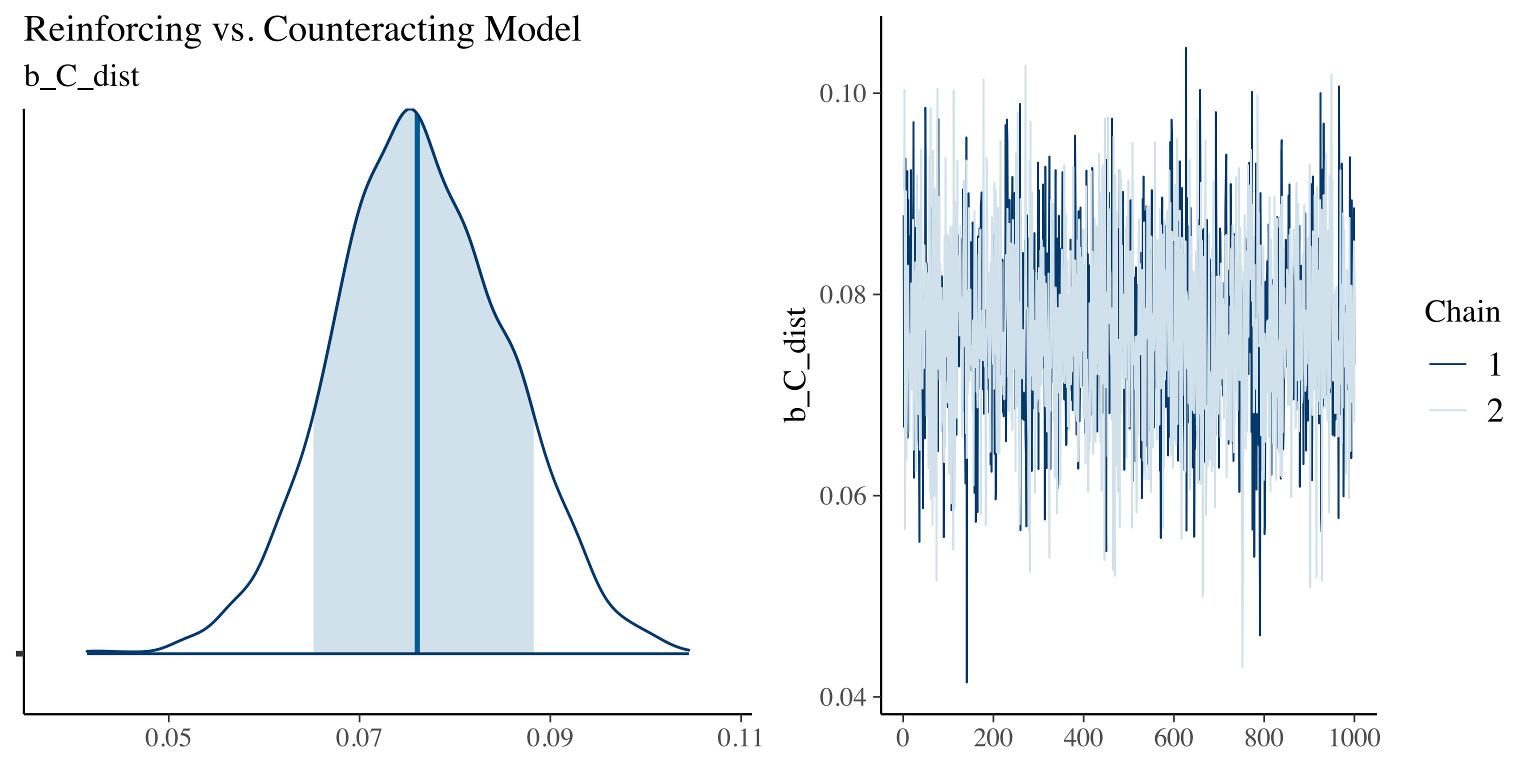

Reinforcing vs. Counteracting Model
b_C_dist
0.05
0.07
0.09
0.11
b_C_dist
0.10
0.08
0.06
0.04
0
200
400
600
800
1000
Chain
1
2

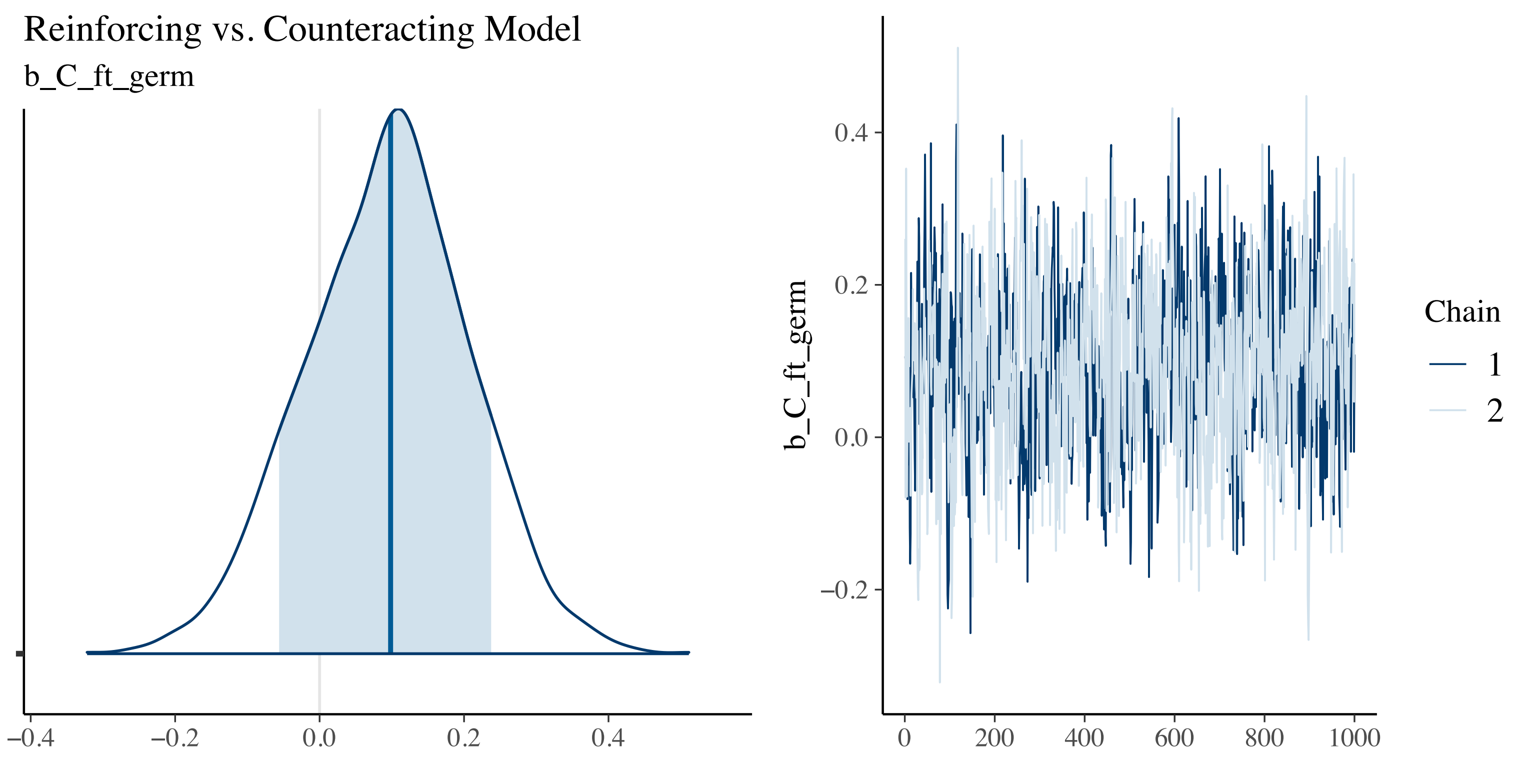

Reinforcing vs. Counteracting Model
b_C_ft_germ
−0.4
−0.2
0.0
0.2
0.4
b_C_ft_germ
0.4
0.2
0.0
−0.2
0
200
400
600
800
1000
Chain
1
2

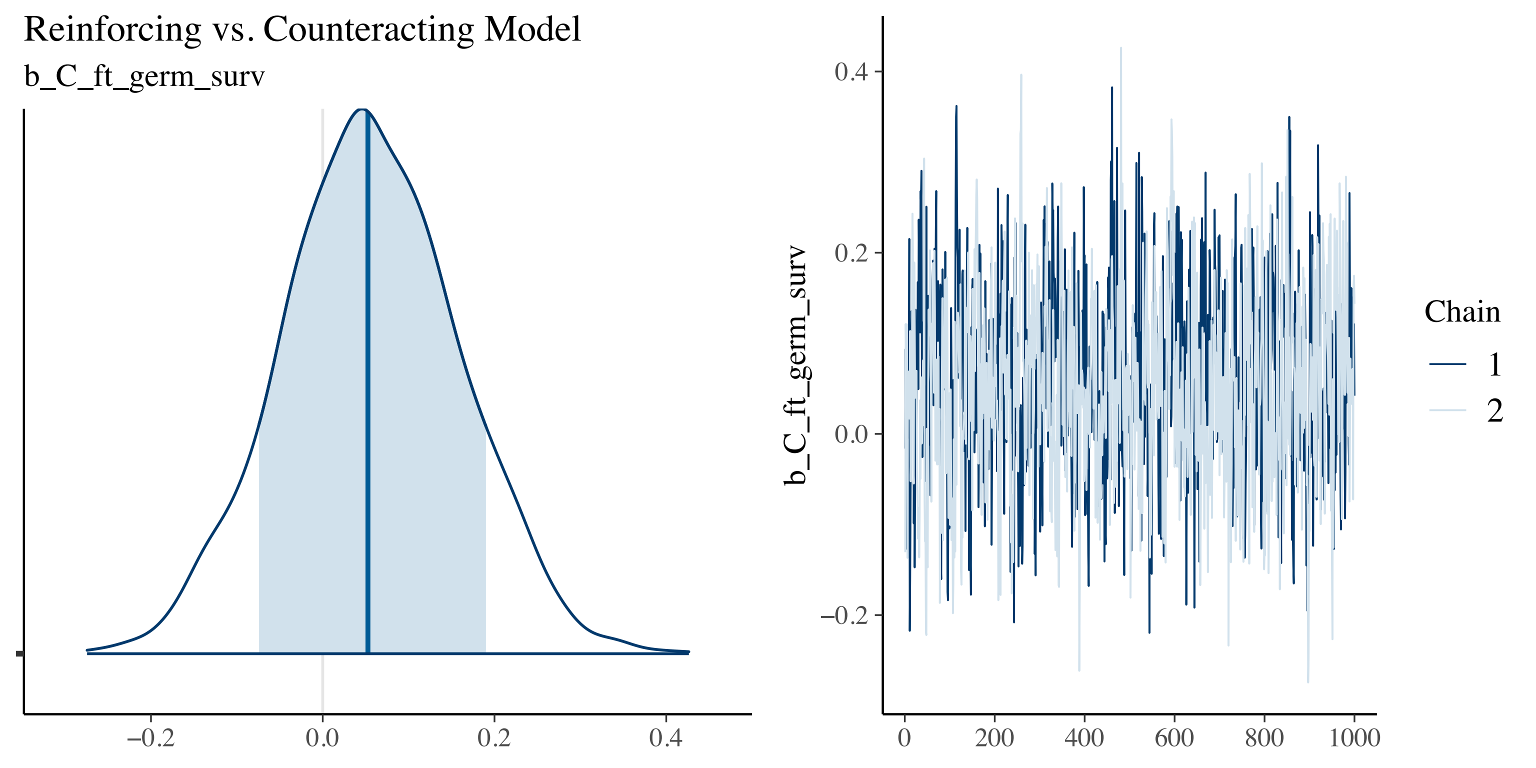
Reinforcing vs. Counteracting Model
b_C_ft_germ_surv
−0.2
0.0
0.2
0.4
b_C_ft_germ_surv
0.4
0.2
0.0
−0.2
0
200
400
600
800
1000
Chain
1
2

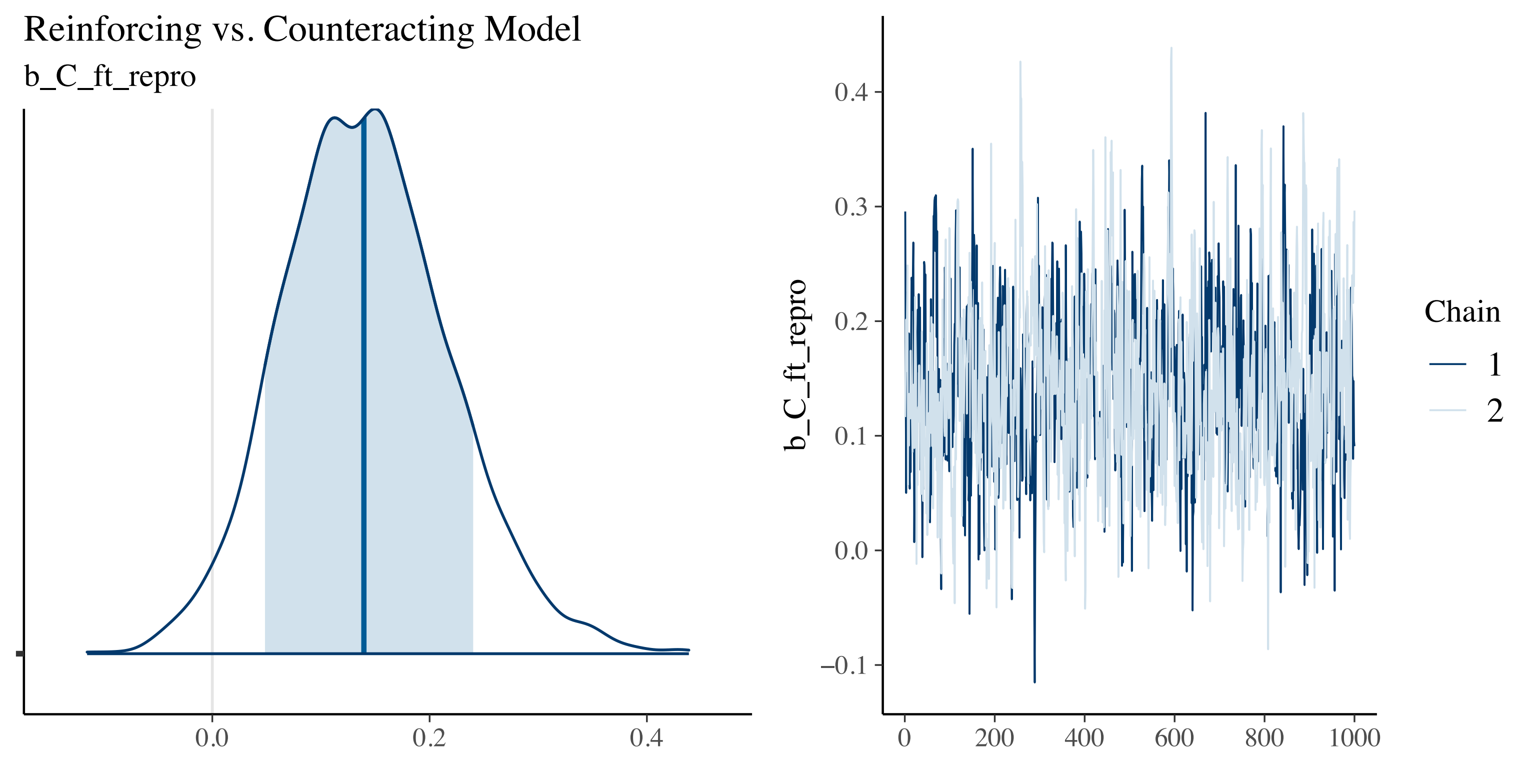
Reinforcing vs. Counteracting Model
b_C_ft_repro
0.0
0.2
0.4
b_C_ft_repro
0.4
0.3
0.2
0.1
0.0
−0.1
0
200
400
600
800
1000
Chain
1
2

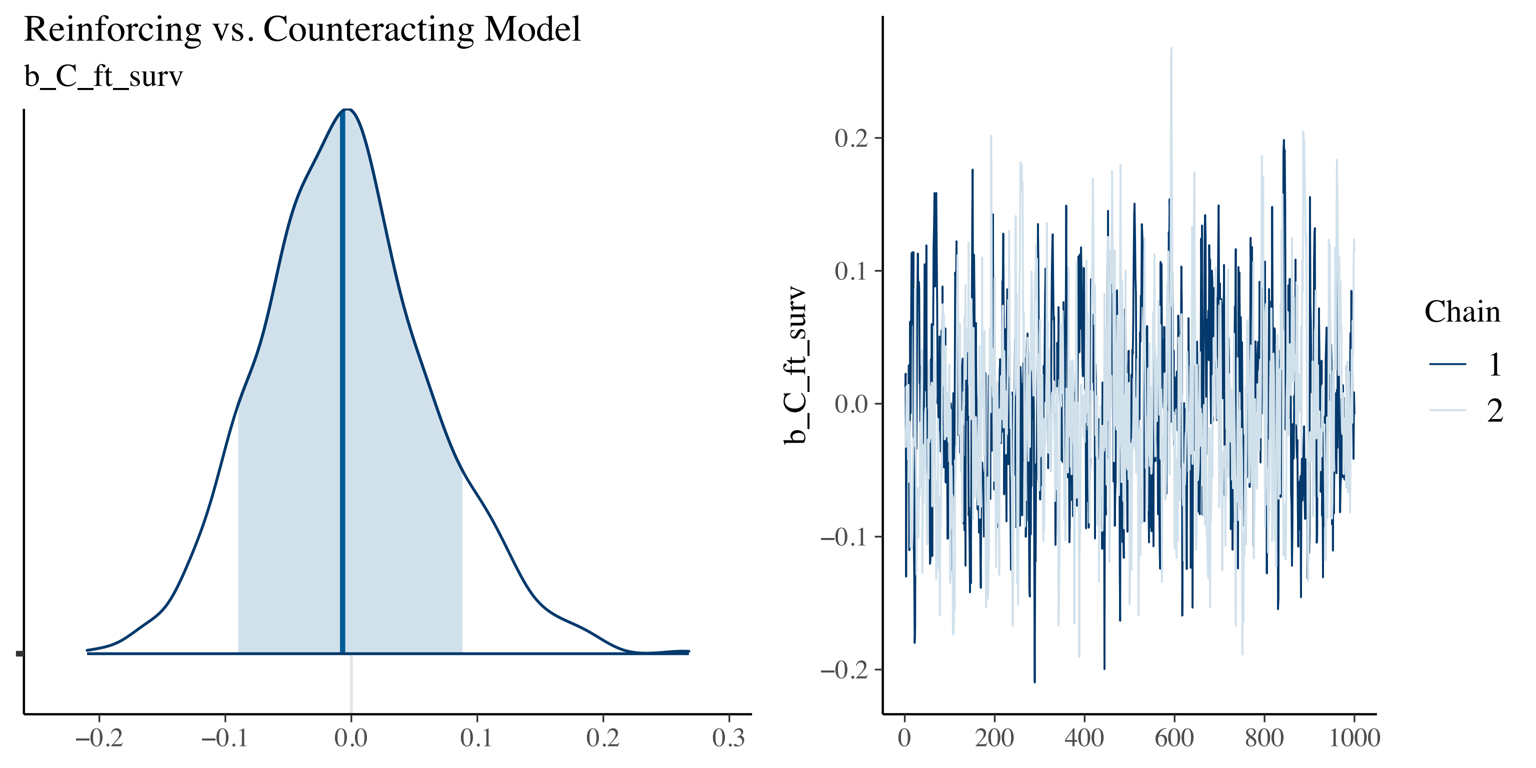

Reinforcing vs. Counteracting Model
b_C_ft_surv
−0.2
−0.1
0.0
0.1
0.2
0.3
b_C_ft_surv
0.2
0.1
0.0
−0.1
−0.2
0
200
400
600
800
1000
Chain
1
2

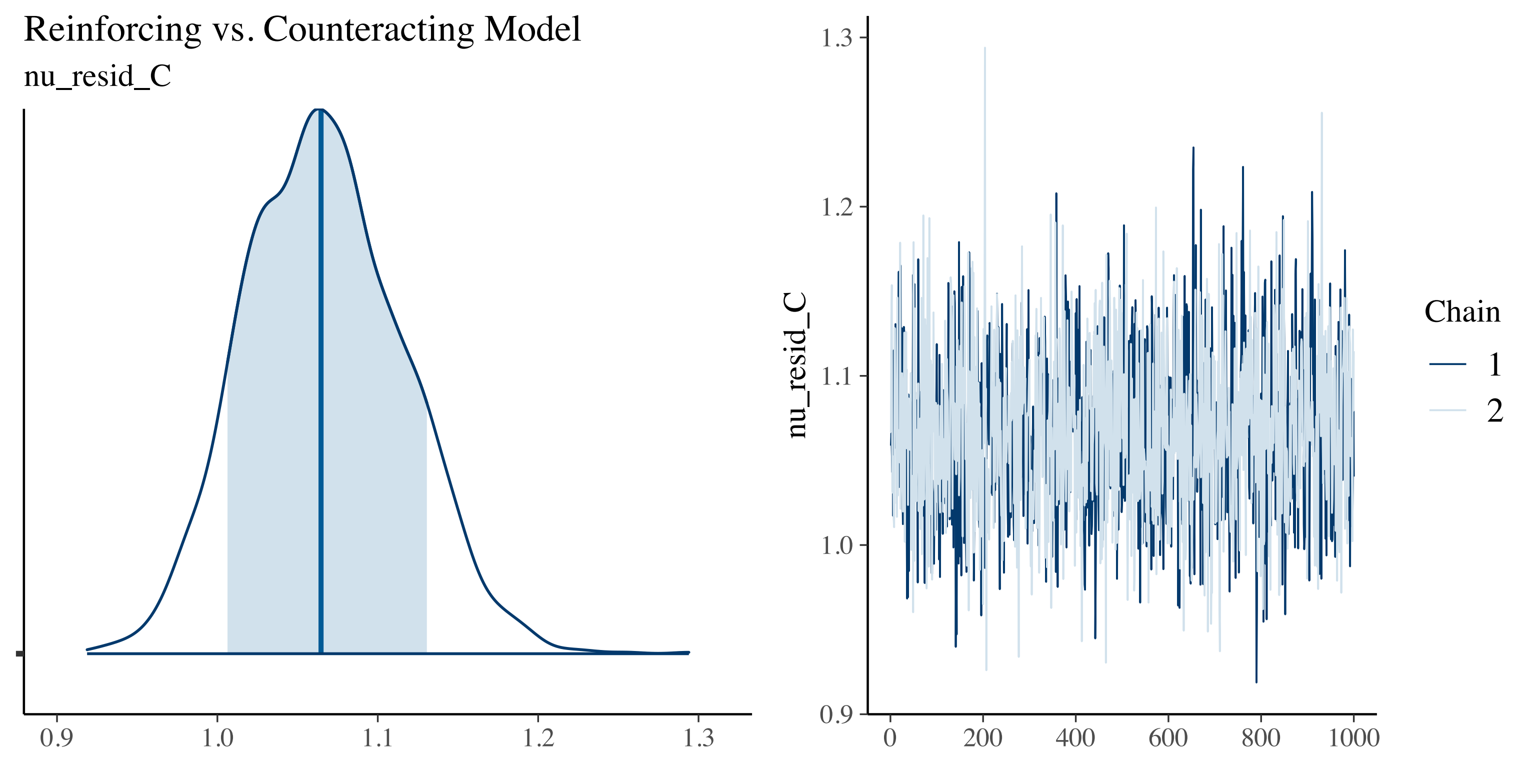

Reinforcing vs. Counteracting Model
nu_resid_C
0.9
1.0
1.1
1.2
1.3
nu_resid_C
0
200
400
600
800
1000
Chain
1
2

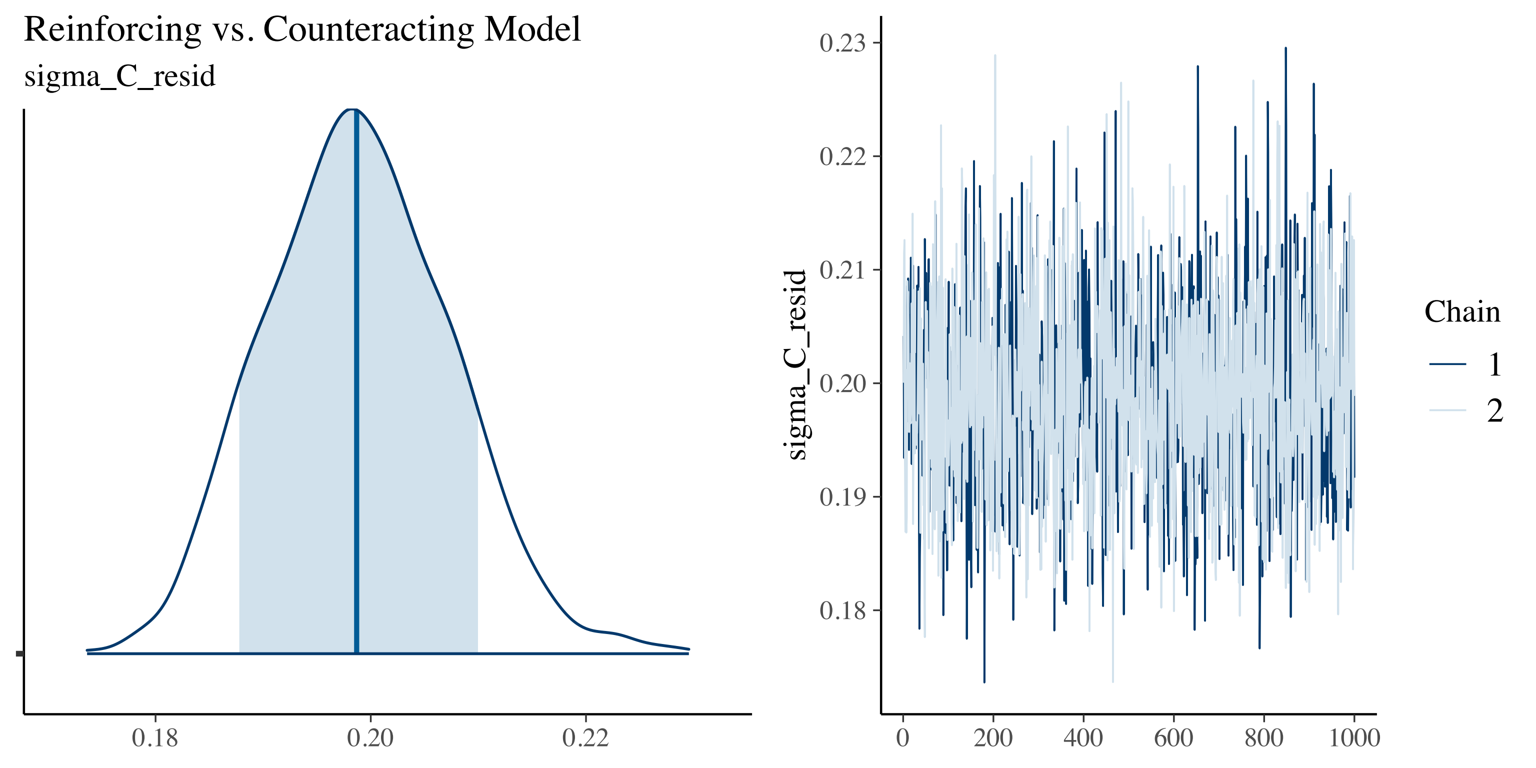

Reinforcing vs. Counteracting Model
sigma_C_resid
0.18
0.20
0.22
0.23
0.22
0.21
0.20
0.19
0.18
sigma_C_resid
0
200
400
600
800
1000
Chain
1
2

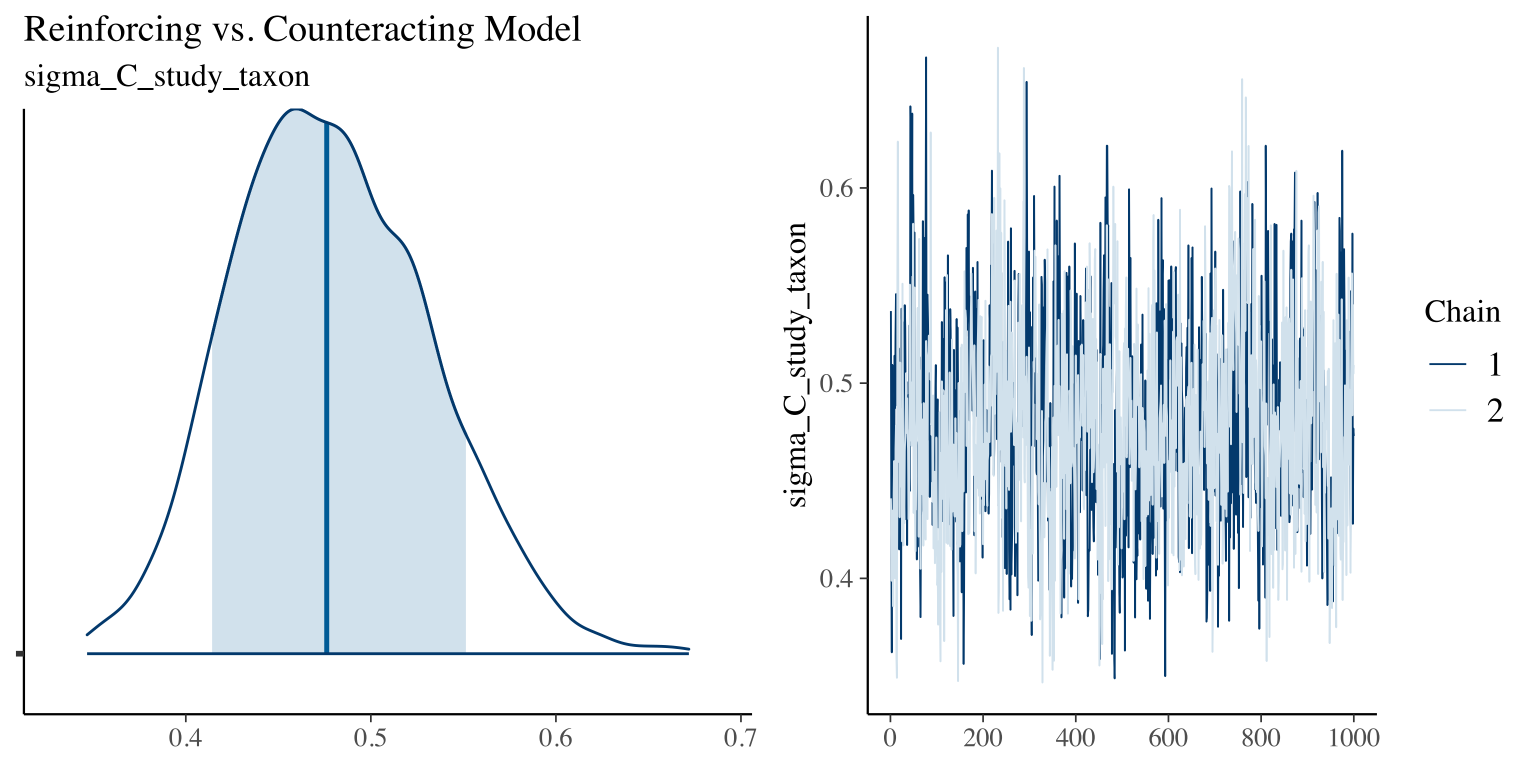
Reinforcing vs. Counteracting Model
sigma_C_study_taxon
0.4
0.5
0.6
0.7
sigma_C_study_taxon
0.6
0.5
0.4
0
200
400
600
800
1000
Chain
1
2

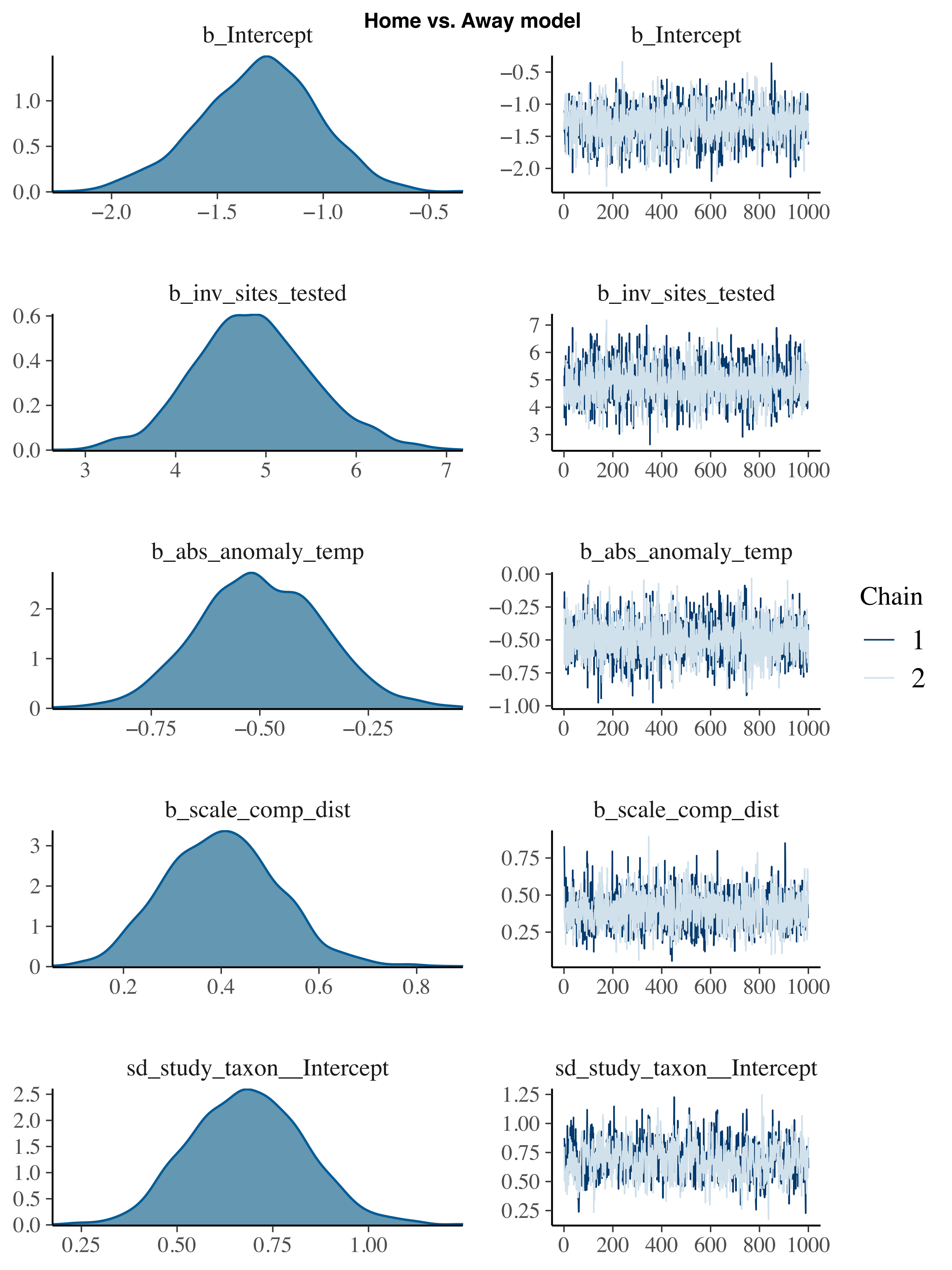
Home vs. Away model
b_Intercept
b_Intercept
b_inv_sites_tested
b_inv_sites_tested
b_abs_anomaly_temp
b_abs_anomaly_temp
b_scale_comp_dist
b_scale_comp_dist
sd_study_taxon__Intercept
sd_study_taxon__Intercept
Chain
1
2

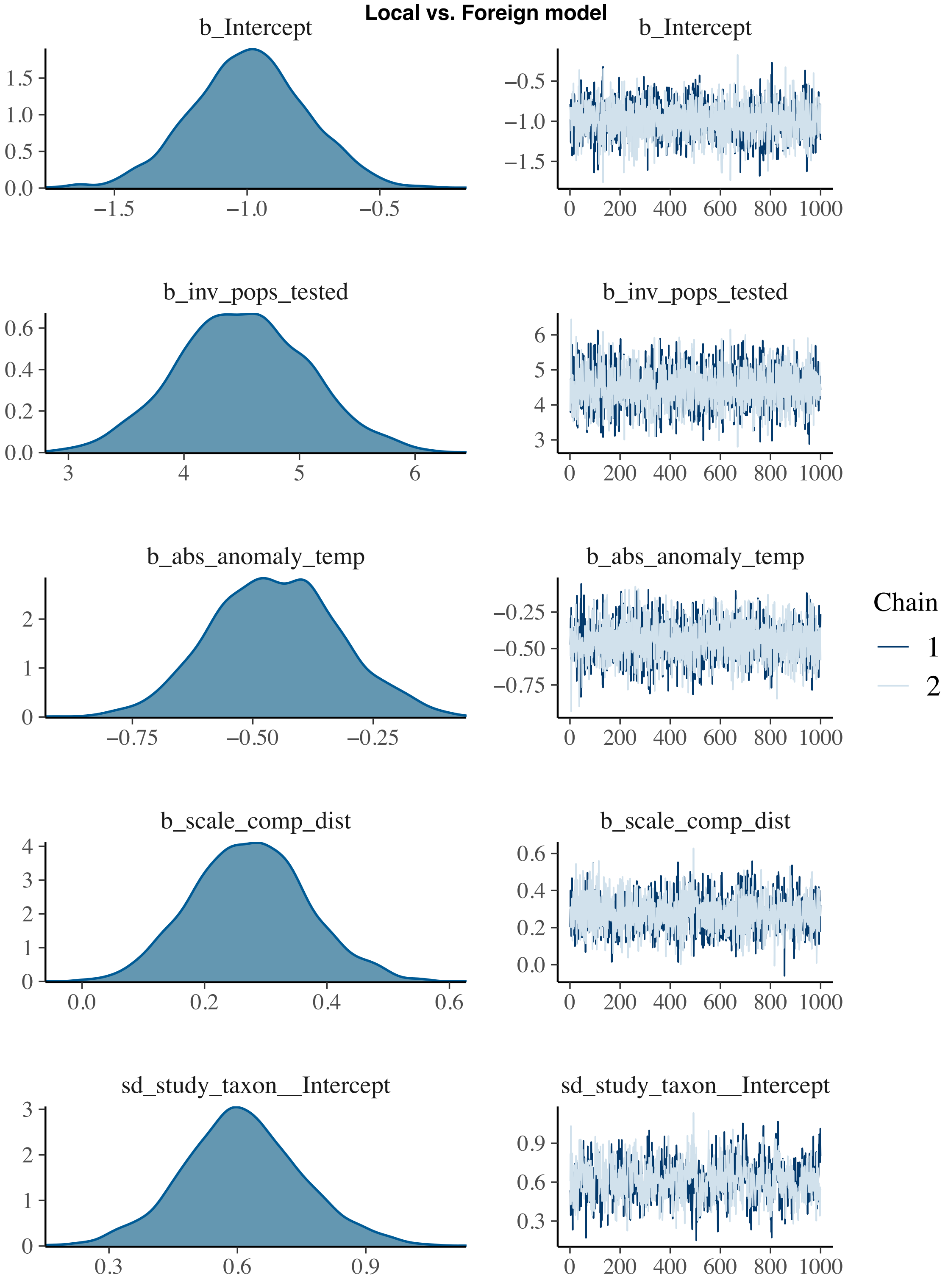

Local vs. Foreign model
b_Intercept
b_Intercept
b_inv_pops_tested
b_inv_pops_tested
b_abs_anomaly_temp
b_abs_anomaly_temp
Chain
1
2
b_scale_comp_dist
b_scale_comp_dist
sd_study_taxon__Intercept
sd_study_taxon__Intercept
